# Early establishment acts as a selective filter shaping climate-associated genomic variation in European beech

**DOI:** 10.64898/2026.09.02.748467

**Authors:** Marieke Lenga, Lars Opgenoorth, Katrin Heer, Christian Lampei, Mona Schreiber

**Author notes:** Mona Schreiber and Christian Lampei should be considered joint senior authors.

## Abstract

Climate change is increasing drought and heat stress in European forests, raising concerns about the capacity of long-lived tree species to respond to rapidly changing environmental conditions. While local adaptation has been documented in many forest trees, it remains unclear whether newly established seedlings, which form the forests of the future, are able to persist and adapt to these new climatic conditions. Here, we investigated genomic differences between naturally regenerated seedlings and trees of European beech (*Fagus sylvatica*) across the three regions of the German Biodiversity Exploratories using low-coverage whole-genome sequencing (∼5x) of 1,032 individuals. Population structure was primarily driven by geographic region, whereas genetic diversity was similar across life stages. Despite this genome-wide similarity, we detected allele frequency shifts between trees and seedlings, concentrated in narrow genomic windows. These shifts were strongest in surviving seedlings, suggesting that environmental filtering during early establishment may contribute to shaping the genetic composition of regenerating populations. The strongest signals were observed within the Swabian Alb, where sampled seedlings were 2-years old and had experienced a longer period of potential filtering prior to sampling. Genotype–environment association analyses identified loci associated with climatic variables, and subsequent GO enrichment analyses of genes linked to these loci revealed significantly more enriched GO terms in seedlings than in trees, suggesting stronger environmental filtering by the current climate in seedlings. In particular, we found associations with maximum air temperature, relative humidity, soil moisture, and precipitation, affecting genes involved in stress responses, growth, metabolism, and developmental processes. Together, our results demonstrate that young cohorts of European beech differ genetically from trees and reveal genomic patterns consistent with life-stage-dependent environmental filtering. These findings suggest that the genetic composition of early life-stages is already altered by current environmental conditions, possibly contributing to adaptation to new climatic conditions.

## 1 Introduction

Extreme climate events, such as the 2018/2019 drought and heat wave in Europe, have caused widespread growth declines, elevated tree mortality, and shifts in forest structure and function with impacts expected to intensify under continued climate change (Martinez del Castillo et al., 2022; Senf & Seidl, 2021). Consequently, there is a growing urgency to better understand the processes that enable forest persistence under rapid climate change. Long-lived temperate broad-leaved tree species are generally considered genetically resilient due to their large effective population sizes, extensive gene flow, polygenic architectures, and high levels of standing genetic variation, all of which can facilitate adaptive responses to environmental change (Kremer et al., 2025). Accordingly, models incorporating adaptive capacity suggest that forest trees may partly mitigate climate change impacts through adaptation, phenotypic plasticity, and gene flow (Benito Garzón et al., 2011, 2019; Gárate-Escamilla et al., 2019; Valladares et al., 2014), whereas projections based on species distribution models, which generally do not account for adaptive evolution, phenotypic plasticity, or adaptive gene flow, often predict greater climate-driven range shifts (Dyderski et al., 2018; Hanewinkel et al., 2013). However, whether ongoing selection is already altering the genetic composition of natural populations remains poorly understood.

In Central Europe, *Fagus sylvatica* L. (European beech, hereafter beech) is a dominant forest tree species that until recently was regarded as relatively resistant to rapidly changing climate. However, recent drought and heat events have revealed substantial vulnerability, particularly in older stands, resulting in growth decline, crown damage, and increased mortality (BMEL, 2023; Leuschner, 2020; Miranda et al., 2022; Rohner et al., 2021; Schuldt et al., 2020). This raises the question of whether beech is undergoing a general decline in Central Europe or whether the observed loss of vitality reflects selection imposed by a changing climate.

During the 2018 drought, individual beech trees within the same stands showed strongly contrasting responses, with severely drought-damaged trees occurring alongside apparently healthy individuals. The identification of numerous drought-associated SNPs across the genome (Pfenninger et al., 2021), together with evidence for rapid genome-wide selection across generations (Eberhardt et al., 2026), suggests that drought resilience in beech has a polygenic basis, providing potential for adaptive responses. Consistent with this, previous studies have reported evidence for local adaptation and high phenotypic plasticity in European beech (Gárate-Escamilla et al., 2019; Lazic et al., 2024). However, whether these mechanisms are sufficient to keep pace with ongoing climate change remains uncertain. The polygenic architecture of adaptive traits and the strong population structure across its distribution range, complicate the detection of climate-associated loci, indicating that current genomic approaches capture only part of the adaptive variation (Lazic et al., 2024). Many landscape genomic studies sample numerous populations but few individuals per population, making demographic differences difficult to distinguish from adaptive genetic variation. Increasing sampling effort within populations, can reduce the relative influence of demographic differences among populations and thereby lower the risk of false-positive associations with climate variables (Slavov et al., 2025).

Further, comparing life stages within local populations in long-lived trees, minimizes the effects of population structure while enabling the detection of ongoing selection, which is expected to be strongest during early establishment (Petit & Hampe, 2006). Heat and drought are expected to alter selective pressures, shifting selection from traits associated with competitive growth towards traits enhancing stress tolerance (Collet & Le Moguedec, 2007; Grossnickle, 2012). However, most genomic studies in forest trees still focus on mature individuals or already established juvenile cohorts. Studies incorporating juvenile cohorts, for example in Pinus cembra, suggest that although alleles beneficial under future climates are already present, shifts in allele frequencies may be too slow to keep pace with ongoing warming (Dauphin et al., 2021). Yet, this study focuses on already established cohorts (13-15 years old) that have passed early-life selection, leaving the earliest stages of establishment poorly understood.

Here, we investigate how selection shapes genomic variation across life stages in European beech by comparing naturally regenerated seedlings, and mature trees across environmental gradients within the Biodiversity Exploratories (https://www.biodiversity-exploratories.de). Within a total of 95 forest plots, we genotyped adult trees and seedlings and monitored the survival of seedlings from one year to another. This allowed us to assess allele frequency shifts between cohorts and between surviving and non-surviving seedlings and thereby assess selection during early establishment. Further, we tested how climatic conditions influence genomic variation across life stages. This approach allows us to evaluate the genomic basis of adaptive responses during establishment and to assess whether these responses are concentrated in specific genomic regions or distributed across the genome.

We hypothesize that (1) selection during early establishment results in allele frequency differences between mature trees and surviving seedlings, and (2) genomic associations with climatic conditions differ between life stages.

## 2 Methods

### 2.1 Experimental Design

We conducted this study within the Biodiversity Exploratories, a long-term research platform comprising one-hectare forest plots across three regions in Germany with a wide elevational and climatic range: Schorfheide-Chorin (SCH), Hainich-Dün (HAI), and the Swabian Alb (ALB). Environmental data at the plot level were obtained through BExIS, the data platform of the Biodiversity Exploratories (Fischer et al., 2010). We focused on European beech (*Fagus sylvatica*) and sampled six mature trees and six naturally regenerated seedlings per plot in a paired design across 95 experimental plots (EPs). By comparing trees and neighboring seedlings within the same plots, our design reduces differences in demographic history and population structure, and thereby weakens confounding neutral divergence (Lotterhos & Whitlock, 2015). In total, we sampled leaves from 1,080 individuals. The leaves were dried in silica gel. In HAI and SCH, we sampled seedlings that established in 2023. Due to low regeneration density in ALB, we included two-year-old seedlings (established in 2021) and determined their age based on bud scale scars (Caccianiga & Compostella, 2012). We selected seedlings randomly where possible or based on availability in EPs with low regeneration density. Each seedling was paired with a nearby adult tree within 5 m (Table S1).

For each seedling, we recorded stem diameter, chlorophyll content, leaf area, canopy openness (DIFN), and the height of the apical, first scar and cotyledons. To assess early-life selection, seedling survival was recorded one year after sampling. Individuals that were dead or could not be found were classified as non-surviving.

### 2.2 DNA Extraction and sequencing

For DNA extraction, dried leaf material was homogenized in a Retsch mill (Retsch MM 301). Genomic DNA was extracted following the ATMAB protocol described by (Bruegmann et al., 2022) with minor modifications (Supplementary Methods S1).

Library preparation and sequencing were performed by BGI Genomics (Hong Kong) using the DNBSEQ-T7 platform. Low-coverage whole-genome sequencing (lcWGS) was conducted using 150-bp paired-end reads (PE150), targeting approximately 5× genome coverage per sample. Of the 1,080 sampled individuals, 1,032 passed library preparation and were successfully sequenced. 48 samples were excluded due to insufficient DNA quality.

### 2.3 Read Processing, Variant Calling, and Filtering

Following sequencing, raw reads were filtered by BGI using SOAPnuke v2.1.6 (Chen et al., 2018). Filtering was carried out with the parameters -n 0.001 -l 10 -q 0.5 --adaMR 0.25 --polyX 50 --minReadLen 150, corresponding to the exclusion of reads with more than 0.1% ambiguous bases (N), reads shorter than 150 base pairs, and reads in which 50% or more of the bases had a Phred quality score below 10. Reads matching ≥25% of the adapter sequence (allowing up to two mismatches) were removed, along with reads containing polyX stretches of 50 bases or more. High-quality clean reads were obtained, with quality values reported using the Phred+33 scale and used for all downstream analyses.

We aligned reads to the chromosome-level *Fagus sylvatica* reference genome Bhaga (Mishra et al., 2022) using BWA-MEM v0.7.17 (Li, 2013). Using SAMtools mpileup and bcftools call v1.17 (Danecek et al., 2021), we performed SNP calling (minimum mapping quality 30, minimum base quality 20), resulting in a final VCF comprising 1,032 individuals. Detailed processing steps and software parameters are provided in Supplementary Methods S2.

We filtered variants using bcftools v1.17 and VCFtools v0.1.13 (Danecek et al., 2011, 2021). After masking low-confidence genotypes (DP < 3 or GQ < 5), we filtered variants based on quality, allele frequency, missing data, heterozygosity, and mean depth (QUAL ≥ 40, AC/AN ≥ 0.01, ≤ 90% missing genotypes, ≤ 90% heterozygous genotypes, mean depth ≤ 100). We then retained high confidence biallelic SNPs (DP ≥ 3, GQ ≥ 20), removed singleton and monomorphic variants (MAC > 1), excluded individuals with genotype missingness exceeding 60%. Detailed filtering thresholds and commands are provided in Supplementary Methods S3. We used the resulting VCF as the target dataset for genotype imputation.

### 2.4 Phasing and imputation

Because low-coverage whole-genome sequencing resulted in substantial missing data (Fig. S1), stringent variant filtering reduced the resolution of population structure. To mitigate this loss of information, while retaining internal haplotype structure, we performed phasing and imputation using Beagle v5.5 (Browning et al., 2018, 2021), which performs well on low-coverage sequencing data (Pasaniuc et al., 2012; Rubinacci et al., 2021). To generate internal reference panels, we filtered the unfiltered VCF for missingness (> 60% missing genotypes per individual, > 20% missing data per variant) and separated the resulting dataset by geographic region (ALB, HAI, SCH). We phased each regional reference panel and then imputed the filtered target dataset (impute=true, gp=true) using the corresponding region-specific reference panel. For downstream analysis we retained variants with imputation quality DR² ≥ 0.8. Principal component analyses of the imputed dataset closely resembled those of the unfiltered dataset, indicating that imputation restored genotype completeness without introducing artefacts in population structure.

Final dataset

After filtering and imputation, the final dataset comprised 3,161,583 high-quality biallelic SNPs across 1,006 individuals (492 trees and 514 seedlings). We used this dataset for all downstream analyses.

### 2.5 Data analysis

For all following data analyses, we used R (R Core Team, **2023**).

We summarized seedling survival as the proportion of individuals surviving one year after sampling within each location. We compared morphological traits between surviving and non-surviving seedlings using linear mixed-effects models with survival status as a fixed effect and plots nested within location as random effects. We tested residuals for departure from normality. Repeating the analyses without 2-year-old seedlings from ALB yielded similar results; therefore, we retained ALB in the final analysis.

#### 2.5.1 Genetic Distances and Diversity

To place signals of allele frequency shifts and genotype-environment associations into the context of neutral population structure and demographic history, we first characterized genetic diversity, differentiation and connectivity among regions and life stages.

To assess population genetic structure, we used principal component analysis (PCA) implemented in EIGENSOFT SmartPCA (Patterson et al., 2006). We ran PCA without outlier removal (numoutlieriter = 0, outliersigmathresh = 6) and retained the first 10 principal components. Using polynomial regression (2^nd^ order) models, we related PCs 1 and 2 to latitude and longitude, downloaded from the Biodiversity Exploratories Information System (BExIS; dataset ID 20907; (Ostrowski et al., 2023).

To further characterize population structure and estimate ancestry proportions among individuals, we used sparse non-negative matrix factorization (SNMF) implemented in the R package LEA v3.18.0 (Frichot & François, 2015). Prior to analysis, we pruned SNPs (dataset 2) for linkage disequilibrium in PLINK (--indep-pairwise 50 kb 5 0.2), to retain approximately independent loci. We ran SNMF for K = 1–20 with 10 repetitions per K and visualized ancestry coefficients for K = 2–4.

We identified private alleles using a custom script (see [Data Availability](https://docs.google.com/document/d/1nd1hvfdYx3ig53NCP_5A6l3bWblxWvhEugsVaDu4M40/edit?tab=t.0#heading=h.atbi85p6axby)) to assess region-specific genetic variation. Because filtering and imputation can remove rare variants, we used the non-imputed dataset filtered only for missingness at the SNP and sample level (dataset 3). Using bcftools (v1.21), we extracted population-specific genotype data and classified SNPs present exclusively in one of the three populations (ALB, HAI, SCH) as private alleles.

To investigate introgression among populations, we performed ABBA–BABA (D-statistics) analyses using the filtered, imputed dataset. We used the *Quercus robur* reference genome assembly V2_2N (Plomion et al., 2018) as an outgroup and calculated D-statistics for all combinations of the three beech populations (ALB, HAI, SCH) with admixr (Petr et al., 2019) and ADMIXTOOLS (Patterson et al., 2012). Analyses included ∼85,000 filtered *Fagus sylvatica* SNPs with orthologous genotypes inferred from the oak genome after alignment to the beech reference genome (Supplementary Methods S4). Significant D-statistics indicate asymmetric derived allele sharing inconsistent with a strictly bifurcating population history. Statistical significance was assessed using a block-jackknife approach (|Z| ≥ 3).

We estimated population differentiation (Weir and Cockerham’s F_st)_, from LD-pruned SNP dataset 2 to reduce bias from linked variants, using VCFtools v0.1.17 (Danecek et al., 2011) in 10 kb non-overlapping windows, separately for trees and seedlings. We calculated nucleotide diversity (π) from both the LD-unpruned imputed and the non-imputed datasets to retain genome-wide variation and assess the effect of imputation.

To assess isolation by distance (IBD) and isolation by environment (IBE), analyses we performed separately for trees and seedlings using mean pairwise F_st_ estimates. We calculated geographic distances between plots as Haversine distances (distHaversine(), R package geosphere) and environmental distances as Euclidean distances based on standardized precipitation, rain days, relative humidity, soil moisture, and maximum air temperature. Using Spearman’s rank correlations (cor.test() R-package stats), we assessed associations between pairwise F_st_ and geographic or environmental distance among plots.

We calculated observed (Ho) and expected heterozygosity (He) for each individual using VCFtools v0.1.17 (Danecek et al., 2011) and summarized values across populations and life stages. Using the (Wang, 2011) estimator (coancestry(), R-package related) (Pew et al., 2015), we estimated pairwise relatedness and considered tree-seedling pairs with a relatedness coefficient ≥ 0.4 as parent-offspring pairs.

#### 2.5.2 Allele frequency differences

To identify SNPs showing allele-frequency shifts between trees and seedlings, we calculated paired allele-frequency differences

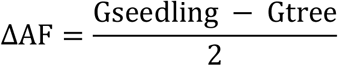

and tested whether the mean ΔAF differed from zero using two-sided one-sample t-tests based on imputed dataset 1. We did not correct population structure because trees and seedlings shared the same population structure within regions (Fig. 3). We performed analyses across all tree– seedling pairs, within regions, by seedling survival status (surviving and dead) across all regions and within each region, for the combined Hainich (HAI) and Schorfheide (SCH) dataset, and for genetically related tree–seedling pairs only. To account for multiple testing, we applied Bonferroni and Benjamini–Hochberg (FDR) corrections and considered SNPs significant at p ≤ 0.05 or q ≤ 0.05. To assess consistency among analyses, we correlated SNP-wise −log10(*p*) values from the analysis including all tree–seedling pairs with those from the analysis restricted to surviving tree-seedling pairs using Pearson and Spearman correlations.

We calculated pairwise linkage disequilibrium (LD) with PLINK v1.9 to examine LD surrounding significant SNPs.

To assess their functional relevance, we estimated LD decay and assigned SNPs to genes if they overlapped annotated gene regions or lay within ±5 kb. We annotated significant SNPs using the *Fagus sylvatica* Bhaga genome annotation (Mishra et al., 2022) and performed Gene Ontology (GO) enrichment analyses with clusterProfiler, testing for over-representation of GO terms among genes linked to outlier SNPs relative to genes linked to all SNPs included in the ΔAF analysis.

#### 2.5.3 Gene-Environment Association analysis

Because genotype–environment association (GEA) analyses can detect loci associated with environmental gradients, including weakly selected variants we performed GEA analyses using dataset 1. We obtained monthly environmental data for the 2023 growing season (March– October) from the Biodiversity Exploratories Information System (BExIS; dataset ID 19007; Wöllauer et al., 2021). Because our aim was to compare current genotype–environment associations between life stages rather than reconstruct historical selection in adult trees, we used the same environmental conditions for seedlings and trees. We aggregated environmental variables to plot-level summaries across the growing season by summing precipitation and rain days and averaging all other variables. The GEA included variables describing water availability, air and soil temperature, and relative humidity (Supplementary Methods S5).

We inferred population structure with sNMF and performed univariate GEA analyses with LFMM2 (R package LEA; Frichot & François, 2015). evaluating K = 1–10 and selecting the best-supported K based on cross-entropy. We ran LFMM2 with K = 2 latent factors and genomic control to account for residual population structure (Caye et al., 2019). and corrected for multiple testing using the Benjamini–Hochberg false discovery rate and the Bonferroni threshold.

To account for differences in seedling establishment year among regions, we repeated the GEA using environmental data from the corresponding growing season (2021 for ALB; 2023 for HAI and SCH).

##### Accounting for false positives

Because GEA methods can produce false positives (Booker et al., 2024; Lotterhos, 2023), we assessed the robustness of LFMM2 results by randomly permuting plot coordinates (Lazic et al., 2024). For each environmental variable, we performed three randomized LFMM2 runs using the same settings (K = 2, genomic control) and applied Benjamini–Hochberg and Bonferroni corrections.

#### 2.5.4 Gene Ontology Enrichment Analysis (GO)

To identify biological processes associated with GEA outlier loci, we performed Gene Ontology (GO) enrichment analyses as described above. We combined over-representation analysis (ORA), testing genes linked to significant SNPs against all genes included in the GEA, with gene set enrichment analysis (GSEA) using mean LFMM *z*-scores of SNPs assigned to each gene. We applied Benjamini–Hochberg correction and restricted gene sets to 5–5000 genes. To assess the influence of temporal environmental resolution, we repeated the GEA, randomization tests, and GO enrichment analyses using environmental data from the year of seedling establishment (2021 for ALB; 2023 for HAI and SCH).

## 3 Results

### 3.1 Seedling survival

Survival differed markedly among regions (Fig. 2 & Table S2). Survival was highest in the two-year old seedlings from Swabian Alb, where 151 of 169 individuals (89%) were still alive one year after the initial sampling. In contrast, of the one-year-old seedlings, only 63 of 183 individuals (34%) survived in Hainich-Dün and 59 of 188 individuals (31%) survived in Schorfheide-Chorin. Across all regions, 273 of 540 sampled individuals (51%) survived, whereas 267 individuals (49%) were classified as dead.

**Figure 1.**
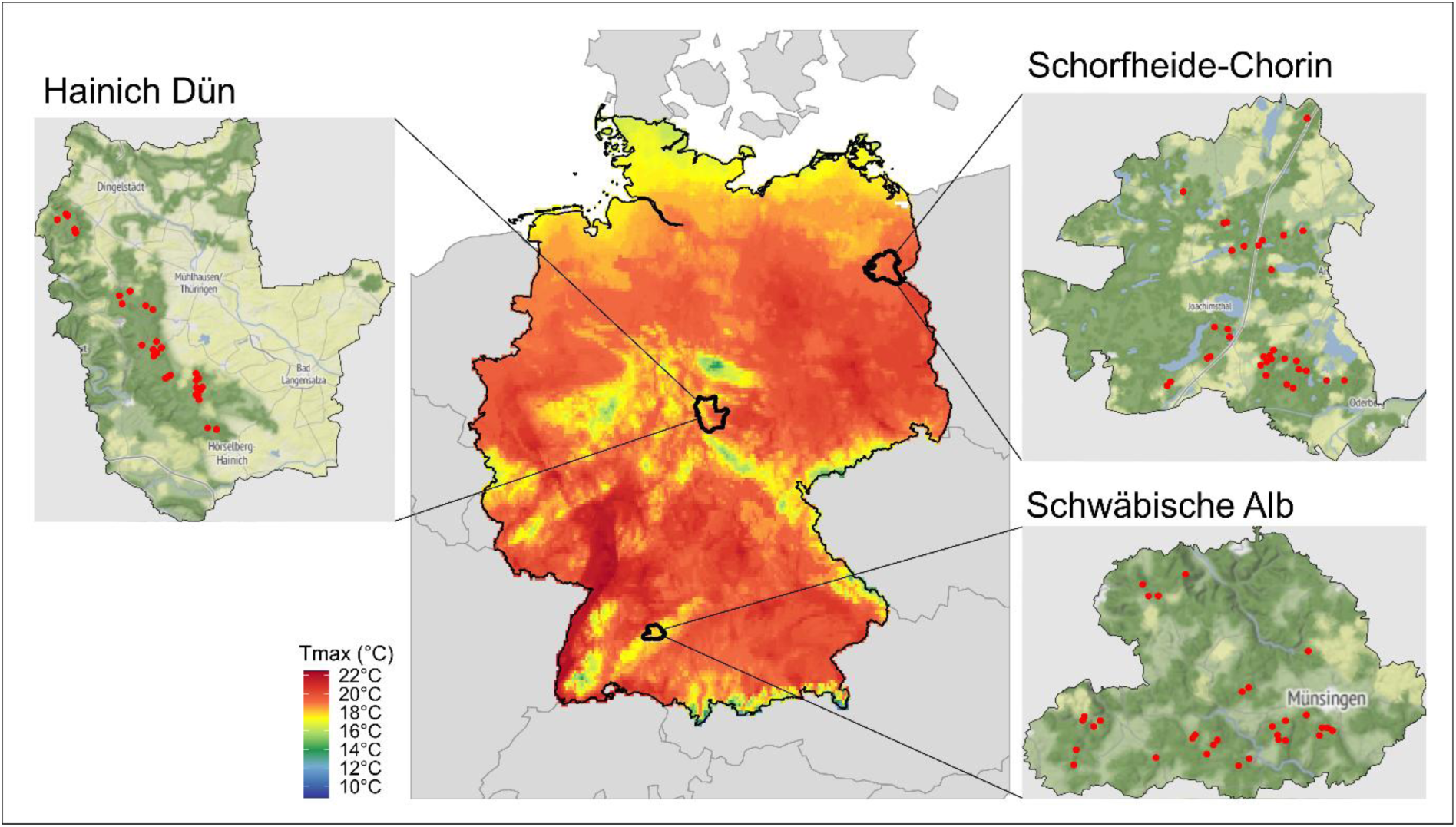
Maximum temperature in the growing season (March – October) across Germany and within the three Exploratories. Maps of the three regions with experimental plots (EPs) indicated as red dots (Swabian Alb: 31 EPs, Hainich-Dün: 31 EPs, Schorfheide-Chorin: 33 EPs). Maximum temperature for the growing season 2023 (March – October) for Germany retrieved from worldclim. Study region boundaries are based on data provided by the Biodiversity Exploratories Information System (BExIS, 2022) and background data obtained from Stadia Maps.

**Figure 2.**
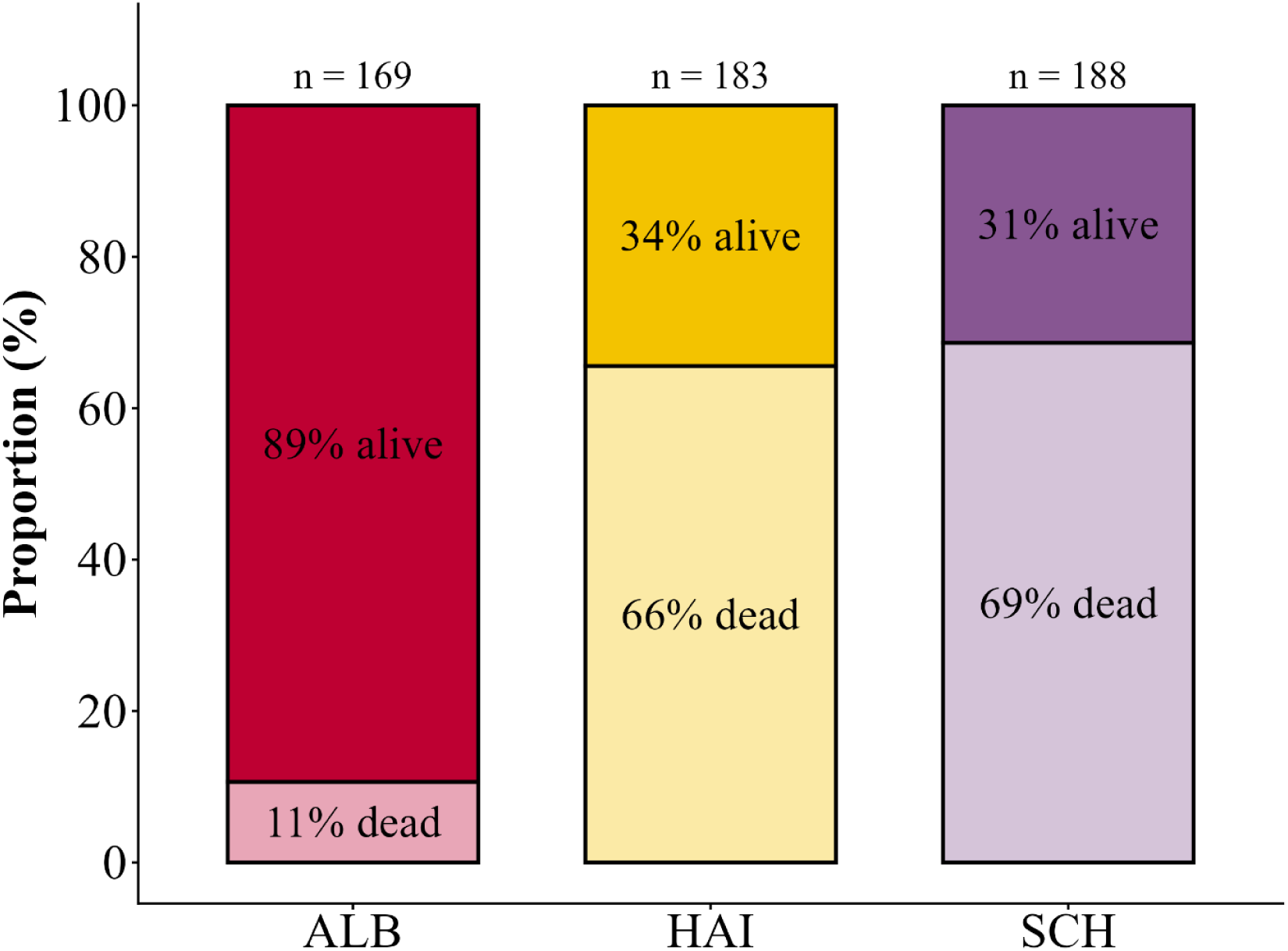
Seedling survival one year after sampling across the three study regions. Survival status of seedlings sampled in May-June 2023 was assessed in April–May 2024. Proportion of dead and alive seedlings one year after sampling in the three study regions (ALB, HAI, SCH). The category “dead” also contains those individuals that were not found again in the second year. Percentages indicate within-region proportions, and numbers above bars indicate total sample sizes (n).

Alive and dead seedlings differed in several morphological traits that we measured in the first year. The strongest differences were observed for stem diameter (F₁,₅₀₃ = 19.65, p < 0.001), leaf area (F₁,₄₃₉ = 12.24, p < 0.001), and chlorophyll content (F₁,₄₈₃ = 11.24, p < 0.001). Canopy openness also differed significantly, although less strongly (DIFN: F₁,₂₁₃.₄₂ = 4.45, p < 0.05) (Table S3 & S4).

### 3.2 Population genetic structure and diversity across tree & seedling and the three study regions

Genetic diversity was similar across regions despite clear geographic structure. Most genetic variation was shared among regions, with private alleles accounting for less than 1% of total variation in each region (Fig. 3d). HAI seedlings harbored more private alleles than adult trees, whereas ALB showed the opposite pattern and SCH showed similar numbers in both life stages. Observed and expected heterozygosity were nearly identical among regions and life stages (Hₒ = 0.292–0.299; Hₑ = 0.297; Table S5).

**Figure 3.**
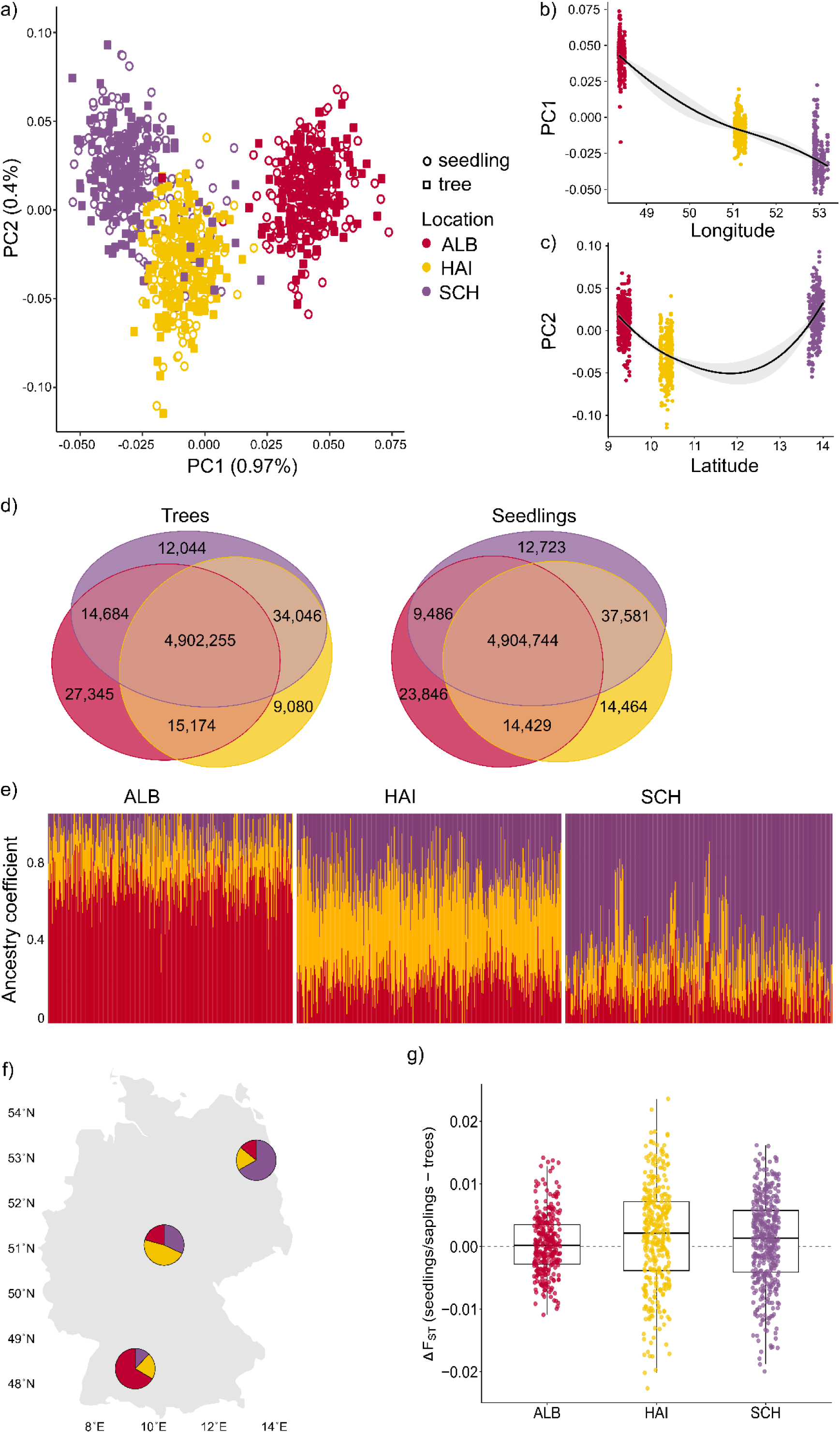
S*p*atial *genetic structure across the three study regions*. *(a)* Principal component analysis (PCA) of all trees and seedlings based on genome-wide SNP data. Individuals cluster by region (ALB, HAI, SCH), with strong overlap between life stages. *(b)* Relationship between PC1 and latitude, showing a strong north south cline in genetic variation (2nd-order polynomial regression, R² = 0.899, p<0.001). (c) PC2 was significantly associated with longitude according to a second-order polynomial regression (R² = 0.47, p < 0.001). Points indicate individual samples colored by region; bold lines with grey shading show the fitted regression and 95% confidence interval. (d) Distribution of shared and private alleles among regions. Venn diagrams of shared and private alleles among ALB, HAI, and SCH for trees(left) and seedlings (right). The central intersection shows alleles shared across all three regions; side intersections indicate alleles shared by two regions; outer sections represent private alleles. For clarity, the large three-way shared category is visually reduced in size. (e) Admixture analysis at K = 3 identifies three major genetic clusters corresponding to ALB, HAI, and SCH. Individuals are shown as vertical bars colored by their ancestry coefficients. ALB and SCH form the most distinct clusters, whereas HAI exhibits mixed ancestry between both. Additional substructure detected at higher (K = 4) and lower (K = 2) is shown in Appendix Fig. S5ab. (f) Geographic distribution of ancestry proportions at K = 3. Pie charts represent the mean ancestry coefficients for each region. ALB and SCH are dominated by distinct clusters (red and purple, respectively), while HAI shows a mixed composition with contributions from all three clusters. (g) Genetic differentiation (ΔF_st_) between trees and seedlings across the three study regions. Genome-wide differentiation between life stages was low in all regions, with slightly higher ΔF_st_ values in HAI than in ALB and SCH.

Principal component analysis revealed clear geographic structure. Individuals clustered by region along the first two principal components, whereas trees and seedlings clustered together within each region (Fig. 3a). PC1 was strongly associated with latitude (R² = 0.887, *p* < 0.001), indicating a north–south genetic gradient, whereas PC2 showed a weaker but significant association with longitude (R² = 0.115, *p* < 0.001; Fig. 3b,c).

Admixture analyses supported the PCA. Cross-entropy identified K = 3 as the best-supported model (Fig. S4), corresponding closely to the three study regions (Fig. 3e,f). ALB and SCH formed distinct genetic clusters, whereas HAI showed mixed ancestry consistent with its intermediate geographic position. At K = 2, ALB separated from HAI and SCH, whereas K = 4 revealed additional substructures within HAI and SCH (Fig. S5).

Nucleotide diversity was identical among populations (π = 2.50 × 10⁻³) and similar between trees and seedlings (Table S6a). Estimates from the non-imputed dataset were slightly lower (π = 2.20 × 10⁻³; Table S6b), consistent with the effect of imputing missing genotypes. Genome-wide ΔF_st_ between trees and seedlings was low within regions, with only slightly elevated values in HAI (Fig. 3g). Between regions F_st_ was lowest between HAI and SCH (F_st_ = 0.0026), intermediate between HAI and ALB (F_st_ = 0.0051), and highest between SCH and ALB (F_st_ = 0.0074) (Table S7), although overall differentiation was low, as expected for populations within Germany.

Isolation by distance was significant across all populations in both trees (Spearman’s *r* = 0.384, *p* < 0.001) and seedlings (*r* = 0.402, *p* < 0.001; Fig. S6a). Isolation by environment was likewise significant for trees (*r* = 0.290, *p* < 0.001) and seedlings (*r* = 0.211, *p* < 0.001; Fig. S6b). In contrast, isolation by distance nor isolation by environment was significant within any of the three regions (Figs. S7, S8).

ABBA–BABA analyses, to investigate introgression, revealed significant asymmetries in derived allele sharing among trees, consistent with strong regional differentiation (Fig. S9). When ALB, HAI, and SCH were assigned as P1, P2, and P3, respectively, a significant negative D-statistic (D = −0.01, Z > 3) indicated greater derived allele sharing between SCH and HAI than between ALB and HAI. Conversely, assigning ALB as P3 (P1 = HAI, P2 = SCH) yielded a significant positive D-statistic (D = 0.01, Z > 3), indicating greater derived allele sharing between ALB and HAI than between SCH and HAI. Together, these results suggest that ALB is the most differentiated region, whereas HAI shares greater genetic affinity with both ALB and SCH. Seedlings showed weaker ABBA–BABA signals than trees, with only one significant test (P1 = HAI, P2 = SCH, P3 = ALB; D = 0.01, Z > 3), consistent with the stronger differentiation of ALB seedlings observed in the other analyses.

### 3.3 Allele frequency differences

Allele-frequency shifts were concentrated in surviving seedlings vs. adult trees and largely absent in dead seedlings vs. adult trees. Across all tree - seedling pairs (n = 478 pairs), we identified 45 outlier SNPs showing significant allele-frequency differences (Fig. 4a). When separating pairs by surviving (Fig. 4b) and dead seedlings (Fig. 4c), we observed clear differences. Pairs with surviving seedlings (n = 242 pairs) showed consistent allele-frequency shifts relative to their paired trees (n = 49 outlier SNPs), whereas dead seedlings (n = 236 pairs) showed only little deviation from tree allele frequencies (n = 4 outlier SNPs). These outlier SNPs were located on chromosomes 2, 4, 5, 7, 10, and 12. Correlation analyses of SNP-wise −log10(*p*) values from the full and survival-restricted allele-frequency shift analyses revealed a positive relationship (Pearson *r* = 0.50, Spearman ρ = 0.38; both *p* < 0.005; Fig. S11), indicating a consistency of SNP-wise allele-frequency shift signals between the two analyses. Linkage disequilibrium around these loci was generally low (Figs. S16), suggesting that the detected allele-frequency shifts were distributed across multiple largely independent genomic regions rather than concentrated within a few highly linked haplotypes.

**Figure 4.**
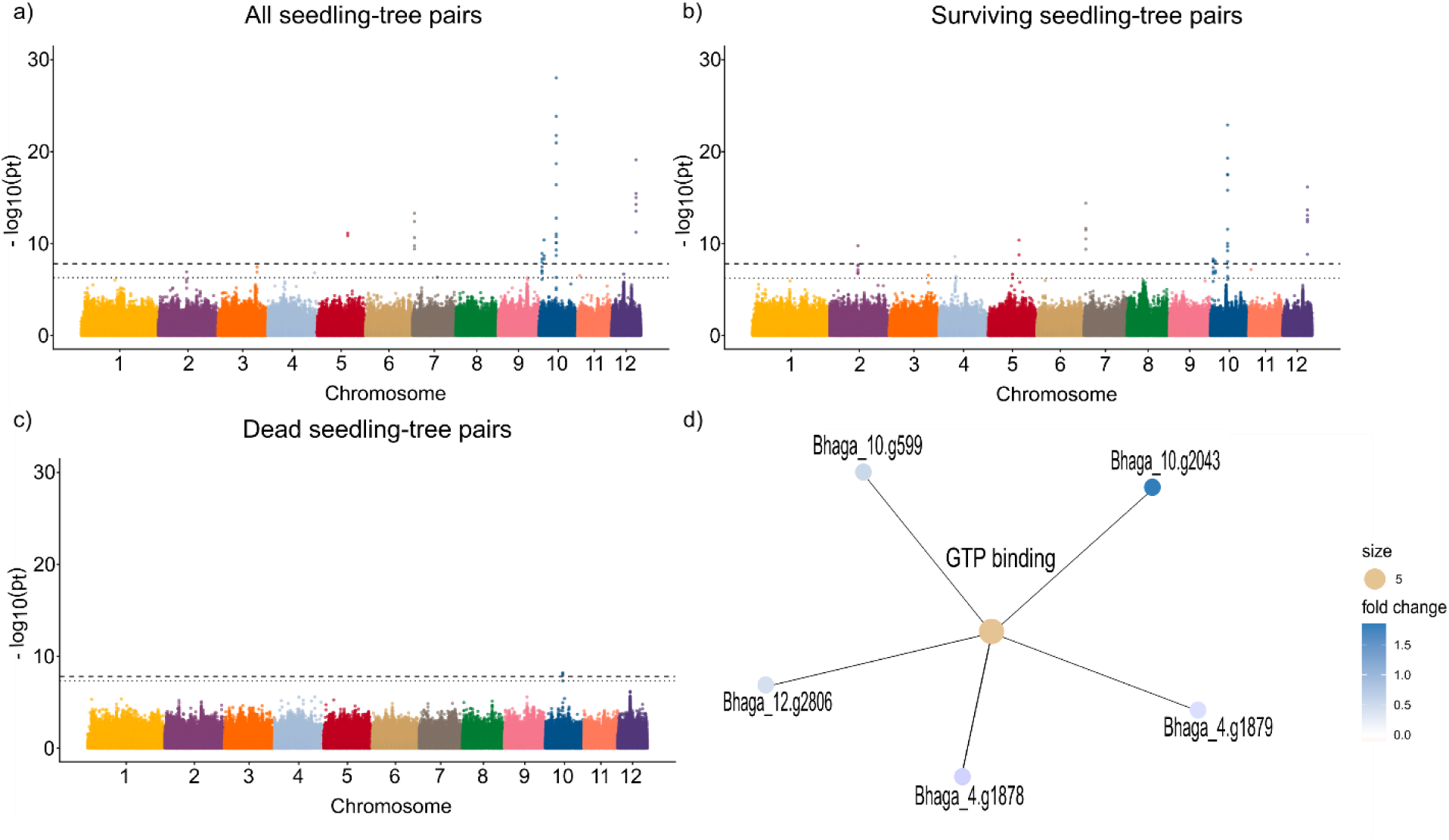
Allele-frequency shifts between trees and seedlings. (a–c) Manhattan plots of allele-frequency differences between seedlings and their paired trees. Points are colored by chromosomes. The dashed line represents the Bonferroni-corrected genome-wide significance threshold, and the dotted line indicates the false discovery rate (FDR) threshold. (a) All pairs (n = 478), identifying 45 outlier SNPs distributed across chromosomes 2, 3, 5, 7, 10, 11, and 12. (b) Pairs with surviving seedlings (n = 242), identifying 49 outlier SNPs across chromosomes 2, 3, 4, 5, 7, 10, 11, and 12. (c) Pairs with dead seedlings (n = 236), identifying four outlier SNPs located on chromosome 10. (d) Gene Ontology (GO) enrichment network (cnetplot) for genes associated with outlier SNPs in surviving seedlings. The GO term GTP binding is enriched and connected to five genes.

Among the 49 SNPs showing significant allele-frequency differences between trees and surviving seedlings, 45 were located within annotated genes, corresponding to 10 unique genes (Table S8). Gene ontology over-representation analysis identified GTP binding (GO:0005525) as the only significantly enriched GO term, represented by five genes (Fig. 4d). These genes are involved in membrane trafficking (Rab7 and ADP-ribosylation factors), cytoskeletal organization (TUB4), and plastid protein synthesis (chloroplastic elongation factor TuB) (Table 2). Four of these genes have previously been reported to be involved in responses to biotic or abiotic stresses (Agarwal et al., 2009; Chun et al., 2021; Fu et al., 2012; Sui et al., 2017).

**Table 1.** Overview of sequencing and variant-calling statistics, including raw reads, mapped reads, mapping rate, and SNP numbers before and after filtering and imputation.

| Metric | Mean value | SNP Dataset |
| --- | --- | --- |
| Mean no. Raw reads | 23,738,718 | - |
| Mean no. Mapped reads | 21,727,303 | - |
| Mean mapping rate % | 91.53 | - |
| No. SNPs unfiltered | 9,683,432 | - |
| No. SNPs after missingness filtering | 4,976,121 | Dataset 3 |
| No. SNPs after filtering not imputed | 1,605,676 | - |
| No. SNPs after filtering, imputed | 3,161,583 | Dataset 1 |
| No SNPs after LD pruning | 147,845 | Dataset 2 |

**Table 2.**
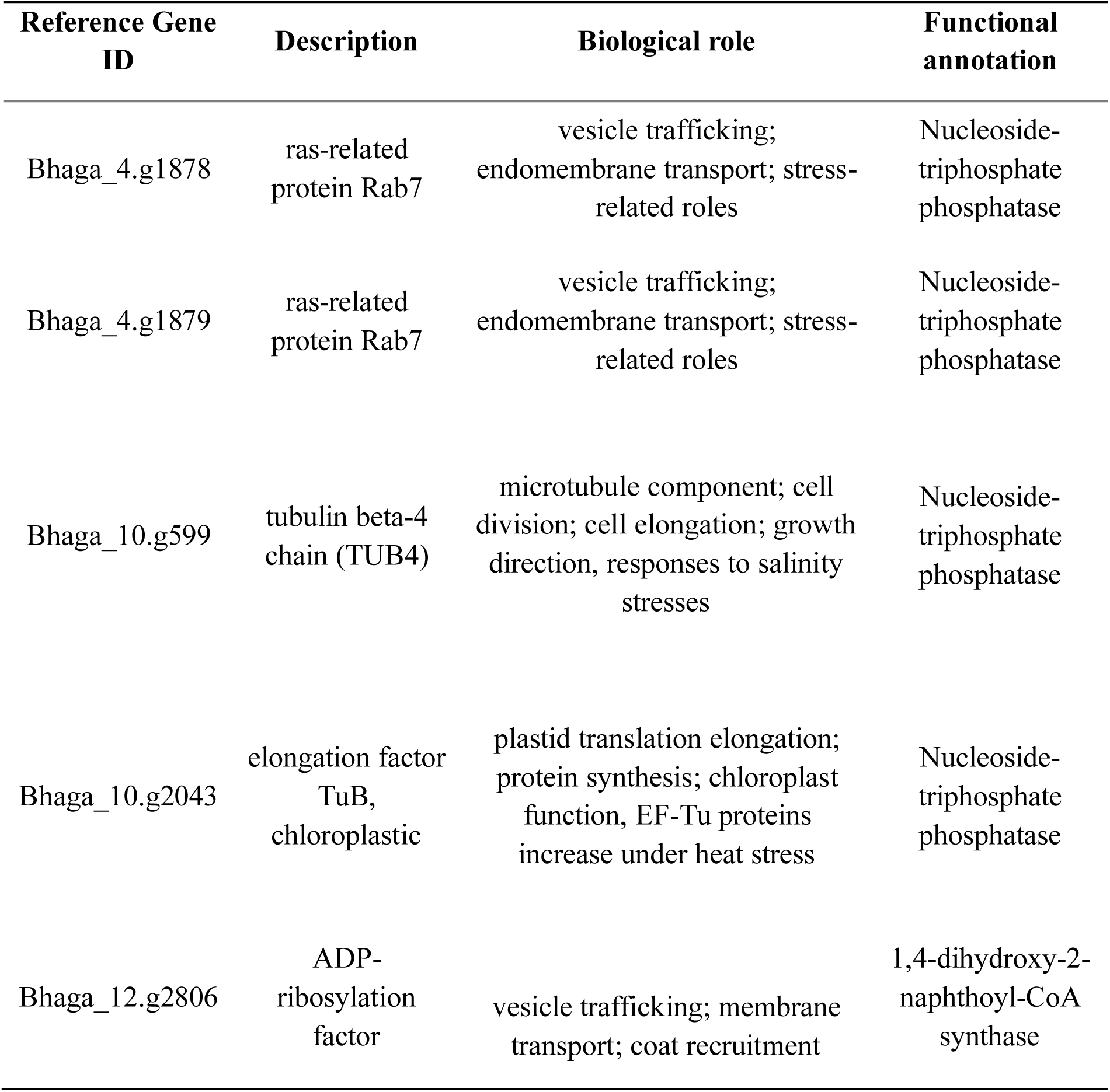
Genes associated with the significantly enriched GO term "GTP binding" identified by overrepresentation analysis (ORA), including gene IDs, described biological roles, and functional annotation.

Restricting the analysis to closely related tree–seedling pairs (Wang relatedness ≥ 0.4) (Fig. S10) yielded similar results for all pairs (n = 252 pairs) and surviving pairs (n = 128 pairs), including the same outlier SNPs on chromosomes 10 and 12 and the absence of outlier SNPs among dead seedling-tree pairs (n = 124 pairs) (Fig. S14a, b, c). However, fewer significant outlier SNPs were detected when using only closely related pairs (13 in all pairs; 9 in surviving pairs), consistent with the smaller sample size.

Separating the analysis by region showed that allele-frequency shifts varied among regions, with the strongest differentiation observed in two-year-old seedlings from ALB (n = 150 pairs), whereas patterns in HAI (n = 163 pairs) and SCH (n = 165 pairs) were weaker (Fig. S12). In the survival-restricted analysis, the number of surviving tree–seedling pairs differed substantially among regions (ALB: n = 134 pairs; HAI: n = 57 pairs; SCH: n = 51 pairs), potentially reducing statistical power in HAI and SCH. Nevertheless, in non-surviving seedlings no distinct outlier signals were detected in either HAI or SCH (ALB: n = 16 pairs; HAI: n = 106 pairs; SCH: n = 114 pairs), despite the substantially larger sample sizes in HAI and SCH (Fig. S13). This suggests that the allele frequency differences detected in surviving seedlings might indeed indicate signals of selection.

To further corroborate this finding, we combined seedling-tree pairs from HAI and SCH (n = 328) and repeated the analysis (Fig. S15). The outlier signal on chromosome 10 persisted and remained weakly detectable among pairs containing surviving seedlings only (n = 108). When restricting the analysis to pairs containing dead seedlings (n = 220) only from these two regions, the signal was absent.

### 3.4 Gene environmental associations (GEA)

Environmental variables differed among regions and varied slightly among plots within regions (Fig. 5c; Fig. S18 & S19). GEA analyses identified significant associations between SNPs and multiple environmental variables in both life stages (Fig. 5 & 6). No SNPs overlapped with those identified in the allele-frequency shift analysis between seedlings and trees, consistent with the different objectives of the two approaches: allele-frequency shift analyses identify loci showing consistent differences between life stages, whereas GEA identifies loci associated with environmental variation within a life stage. The number of outlier SNPs varied among environmental variables and between life stages (Table S9). Mean growing-season temperature, for instance, was associated with 3004 outlier SNPs in seedlings and 1431 in trees. Across all variables, seedlings showed more outlier SNPs than trees for maximum air temperature and maximum and minimum relative humidity, whereas trees showed more outlier SNPs for soil moisture, precipitation, and rain days. Despite this, ORA detected enrichment across a broader range of environmental variables and functional categories in seedlings, whereas enrichment in trees was restricted to fewer environmental variables and GO terms (Fig. 7). Similar patterns were observed in GSEA (Fig. S22), indicating greater functional diversity of genotype– environment associations in seedlings.

**Figure 5.**
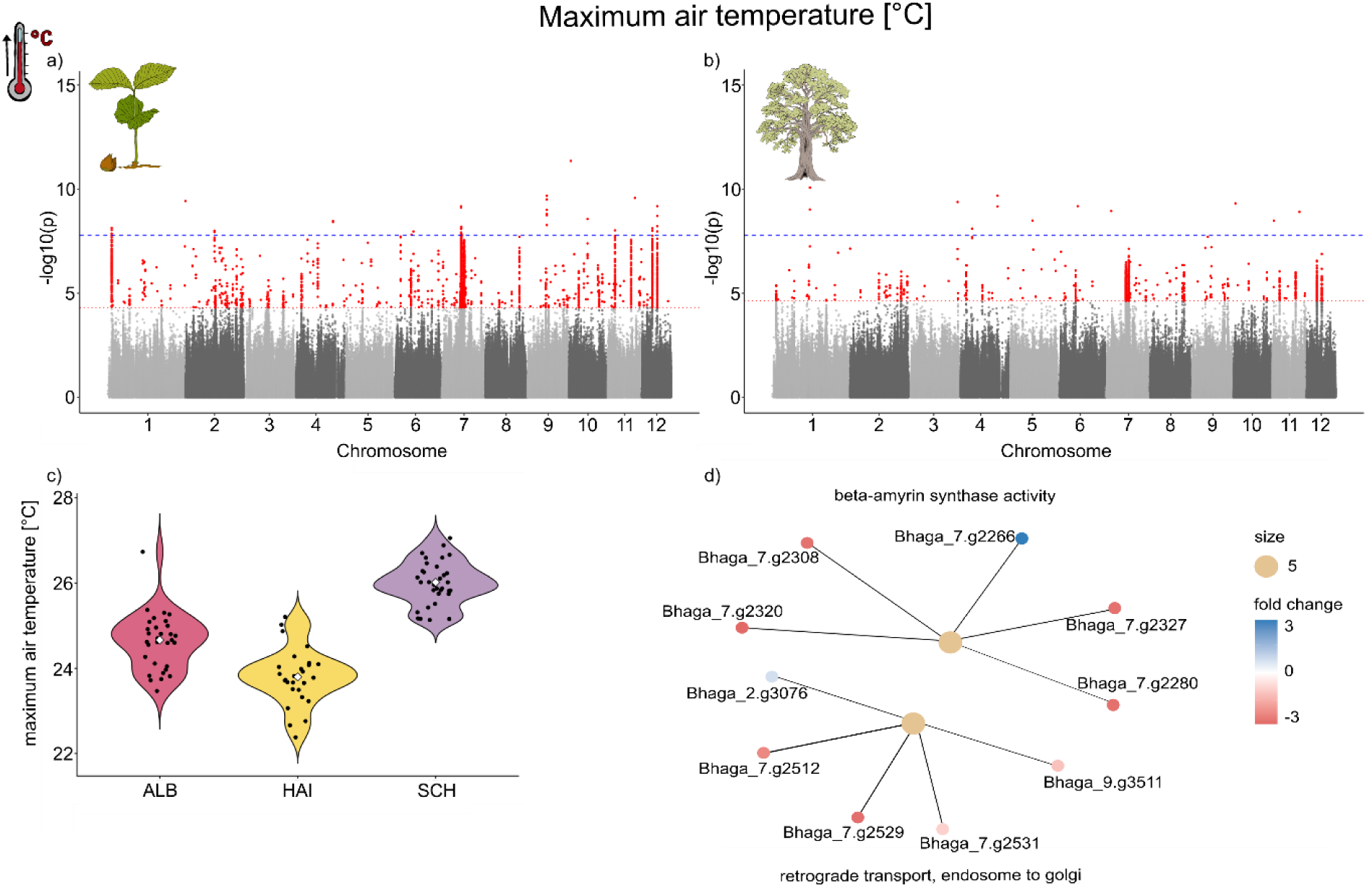
Genotype–environment associations with maximum air temperature during the growing season (March–October 2023). (a–b) Manhattan plots of genotype–environment associations for maximum air temperature at 2 m height in (a) seedlings (n = 499) and (b) trees (n = 477). The blue line represents the Bonferroni-corrected genome-wide significance threshold, and the red line indicates the false discovery rate (FDR) threshold. Seedlings show multiple outlier SNPs exceeding the Bonferroni threshold, with the strongest signal on chromosome 7, whereas trees show fewer outlier SNPs, with most signals below the Bonferroni threshold and fewer exceeding the FDR threshold. (c) Distribution of maximum air temperature across the three study regions, ranging from approximately 22 °C to 28 °C. Temperatures were highest in Schorfheide-Chorin (SCH) and lowest in Hainich-Dün (HAI), with additional variation among plots within regions. (d) Gene Ontology (GO) overrepresentation analysis (ORA) for seedlings, identifying enriched terms including beta-amyrin synthase activity and retrograde transport from endosome to Golgi.

Air temperature and water availability variables differed in the strength and functional composition of their associations. Maximum air temperature showed stronger outlier signals in seedlings and was associated with enrichment only in seedlings. Significant GO terms were β-amyrin synthase activity and retrograde transport from endosome to Golgi (Fig. 5). Across water-related variables, seedlings generally exhibited broader functional enrichment than trees (Fig. 6). Relative humidity was associated with chitin-related, carbohydrate metabolic, and protein turnover processes, whereas soil moisture was linked primarily to lipid- and sterol-related functions in both life stages. Rain days and precipitation showed similar genomic signals between life stages but differed in functional enrichment, with broader enrichment detected in seedlings. Grouping GO terms into broader functional categories revealed consistent enrichment of cell wall/defense, carbohydrate metabolism, protein turnover, and transport/trafficking functions in seedlings, whereas trees showed more restricted enrichment patterns dominated by lipid and sterol metabolism (Fig. 7).

**Figure 6.**
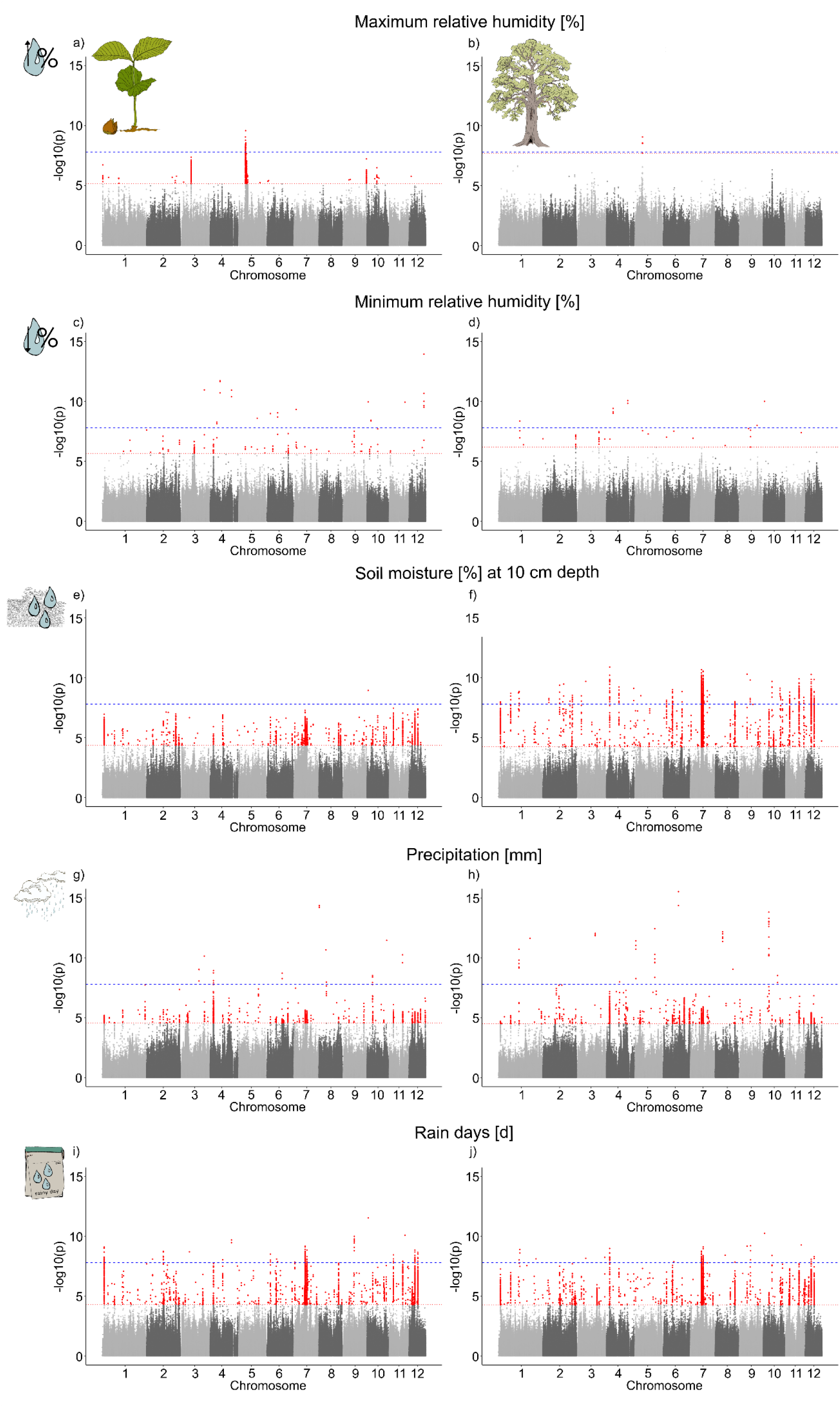
Genotype–environment associations for environmental variables related to water availability, showing variable-specific patterns of outlier SNPs. (a–j) Manhattan plots of genotype–environment associations for multiple climatic variables in seedlings and trees. The blue line represents the Bonferroni-corrected significance threshold, and the red line indicates the false discovery rate (FDR) threshold. Maximum relative humidity is shown for (a) seedlings (n = 447) and (b) trees (n = 442), with more outlier SNPs observed in seedlings. Minimum relative humidity at 2 m height is shown for (c) seedlings (n = 508) and (d) trees (n = 492), also showing a higher number of outlier SNPs in seedlings. Soil moisture at 10 cm depth is shown for (e) seedlings (n = 493) and (f) trees (n = 472), with more outlier SNPs detected in trees. Precipitation is shown for (g) seedlings (n = 514) and (h) trees (n = 492), with a higher number of outlier SNPs in trees. Rain days are shown for (i) seedlings (n = 508) and (j) trees (n = 486), showing broadly similar patterns between life stages.

**Figure 7.**
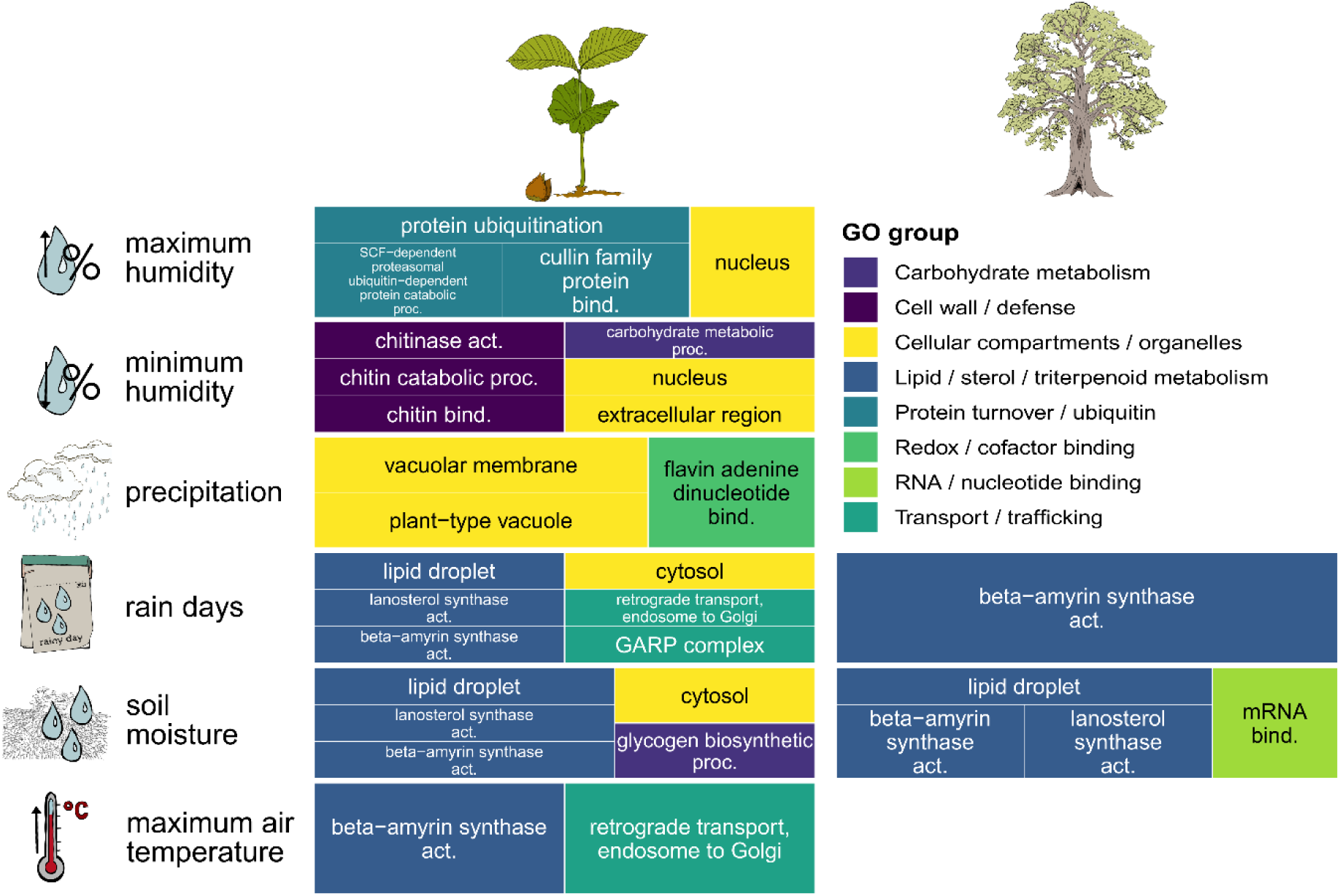
Gene Ontology enrichment of LFMM outlier SNPs (ORA) in seedlings and trees (2023). Treemap plot summarizing all significantly enriched GO terms identified by over-representation analysis (ORA) for each environmental variable. Rows correspond to environmental variables, and columns to life stages (seedlings, trees). Each rectangle represents a GO term, grouped into broader functional categories. In seedlings, significant enrichment was observed across multiple environmental variables, including terms related to cell wall and defense (e.g. chitinase activity, chitin binding, chitin catabolic process), carbohydrate metabolism, and redox/cofactor processes. In contrast, trees showed fewer enriched GO terms, limited to rain days and soil moisture, with enrichment primarily associated with lipid and sterol metabolism (e.g. beta-amyrin synthase activity, lanosterol synthase activity) and membrane-associated cellular components. Overall, seedlings exhibit a more functionally diverse set of enriched biological processes for more diverse environmental variables.

To account for potential false positives in the GEA, we repeated LFMM analyses using randomized geographic coordinates. This randomized approach did not identify any significant outlier SNPs (Fig. S21), suggesting that the observed associations were unlikely to represent false-positive signals. Repeating GEA, ORA, and GSEA using environmental conditions from the year of seedling establishment (2021 for ALB & 2023 for HAI & SCH) produced qualitatively similar results. Although GEA outlier signals were generally stronger in trees under establishment-year conditions (Figs. S25–S26), seedlings consistently exhibited a broader range of enriched GO terms in ORA and GSEA (Figs. S27–S28).

## 4 Discussion

Our results provide genomic evidence that current climate conditions already influence selection during early establishment in European beech. Surviving seedlings, but not dead seedlings, exhibited consistent localized allele-frequency shifts relative to the adult population in stress response related genes indicating that contemporary climatic filtering already alters the genomic composition of the next generation. Independent of these survival-associated shifts, genotype–environment associations revealed marked differences between life stages, with seedlings showing stronger current-climate-associated signals and a broader diversity of enriched biological functions. Together, these findings demonstrate that selection during regeneration shapes adaptive responses of long-lived forest trees to ongoing climate change and are not adequately represented by analyses restricted to mature trees.

### 4.1 Population structure and (low) genetic differentiation

Genetic variation between regions followed geographic gradients, as reflected by strong correlations of the principal components with spatial coordinates and by isolation-by-distance across regions, consistent with patterns observed at the European scale (Lazic et al., 2024). Together with evidence for ongoing, bidirectional gene flow among regions from admixture and D-statistics, with only slightly elevated differentiation in ALB relative to HAI and SCH, these patterns indicate that genetic structure reflects the combined effects of dispersal limitation and historical demographic processes (Postolache et al., 2021). Genome-wide diversity (π, H_e_, H_o_) was highly similar across regions and life stages, consistent with range-wide patterns in *Fagus sylvatica* (Höhn et al., 2021). Low genetic differentiation (F_st_) and few private alleles indicate extensive connectivity and shared standing genetic variation among regions, providing the potential for adaptive responses despite low genome-wide differentiation (Comps et al., 2001; Pluess et al., 2016). Although low-coverage sequencing and stringent filtering may reduce the detection of rare variants and subtle population structure (Lou et al., 2021), genome-wide diversity is only an indirect proxy for adaptive potential, as adaptive variation in beech appears to be concentrated in specific genomic regions (Lazic et al., 2024). Overall, beech populations appear genetically well connected and diverse, but it remains unclear which genetic variants are relevant for survival under ongoing climate change.

### 4.2 Surviving seedlings show shifts in allele frequencies in narrow genomic windows

The observed allele-frequency shifts between paired trees and seedlings were confined to narrow genomic windows and primarily driven by surviving seedlings in the Swabian Alb, where sampled seedlings were two years old and showed higher survival rates. The stronger signals may reflect the cumulative effects of selection during early establishment, since the strongest mortality appears to occur during the first year after establishment, resulting in stronger filtering of the surviving seedlings in the second year, as expected in long-lived tree species (Petit & Hampe, 2006). At the same time, the stronger signal in the Swabian Alb may reflect regional and environmental differences in selection potential, although the relative contributions of environmental conditions and differences in sample composition cannot be disentangled in the present study. Similar geographically localized genomic responses have been reported in beech populations exceeding their historical climatic niche (Eberhardt et al., 2026), indicating that selection intensity may vary strongly among regions. The absence of younger cohorts in ALB may additionally indicate reduced recent recruitment, which is closely linked to masting in beech that can be disrupted under increasing temperatures (Foest et al., 2024). However, the persistence of the chromosome 10 outlier in pairs containing surviving seedlings after excluding ALB from the analysis demonstrates that the observed allele-frequency shift was not restricted to ALB or the two-year-old seedlings sampled there. Overall, analyses restricted to pairs with non-surviving seedlings revealed almost no outlier loci, suggesting that the observed patterns were associated with seedling survival during early establishment in 2023. Consistent with this, surviving seedlings generally showed higher vitality, including higher chlorophyll content, larger leaves, and thicker stems compared to seedlings that were about to die until the next year. Together, these observations are more consistent with selection acting on specific genomic regions than with broad genome-wide divergence between seedlings and trees.

Restricting analysis to closely related tree–seedling pairs reduced the number of detected outliers, but the remaining signals persisted in surviving seedlings and were absent in non-surviving seedlings, consistent with selection on standing genetic variation (Hoban et al., 2016; Lotterhos & Whitlock, 2015). The persistence of these outlier loci despite the higher relatedness further supports the interpretation of selection. Moreover, the observed genomic patterns are comparable to localized selective sweeps, in which positive selection produces narrow genomic signatures around beneficial alleles (Hermisson & Pennings, 2005; Stephan, 2019). Still, distinguishing signals of adaptation from neutral demographic processes remains challenging, as both can generate overlapping genomic signatures, including spurious outliers (Hohenlohe et al., 2010). Further sampling across life stages and environments will be required to disentangle the effects of selection, demography, and cohort structure.

Notably, linkage disequilibrium (LD) across outlier SNPs was low. However, LD generated by selective sweeps is often transient and decays rapidly with recombination, particularly in large, outcrossing populations with high effective recombination rates (McVean, 2007; Stephan, Wolfgang & Hörger, Anja, 2019). In addition, when selection acts on standing genetic variation or multiple haplotypes—as expected under soft selective sweeps—distinct genetic backgrounds can rise in frequency simultaneously, limiting the buildup of extended LD (Hermisson & Pennings, 2005; Pennings & Hermisson, 2006). Together, these processes can produce localized allele-frequency shifts without strong LD signatures.

Moreover, the distribution of outlier loci across multiple chromosomes suggests polygenic adaptation, with selection acting on multiple traits or pathways. In such cases, allele-frequency changes are spread across many loci, but only a subset shows shifts large enough to be detected as outliers (Barghi et al., 2020). The identified loci may therefore represent only the strongest signals, while additional smaller-effect allele-frequency shifts remain undetected (Metheringham et al., 2025).

Taken together, these results support selection during early life stages, with allele-frequency shifts linked to seedling survival and confined to specific genomic regions. If the selective filtering observed here reoccurs across successive seedling cohorts, changing climatic conditions may gradually reshape allele frequencies and contribute to adaptive responses in beech populations. Comparable cohort-based studies in long-lived tree species reveal ongoing genomic responses to climate change, yet also indicate that the rate of allele-frequency change may lag behind the pace of environmental change (Dauphin et al., 2021; Eberhardt et al., 2026). Thus, although our results provide evidence for ongoing adaptive responses, whether these changes are occurring fast enough to keep pace with climate warming remains uncertain.

### 4.3 Seedlings and trees differ in their correlation with current climate

Long-lived organisms with extended generation times are often subject to an adaptational lag relative to current (Browne et al., 2019) or future climatic conditions (Wilczek et al., 2014), potentially leading to mismatches with rapidly changing environments driven by anthropogenic climate change (Hegerl et al., 2019). Understanding the genetic basis of adaptive responses to such change therefore remains a major task (Franks & Hoffmann, 2012).

Seedlings showed stronger and more functionally diverse genotype–environment associations (GEA) with specific environmental variables. Because seedlings represent a single recent cohort, their genomic composition may more closely reflect selection under current climatic conditions, whereas adult trees established across many years and selective environments. In long-lived species with overlapping generations such as beech, this temporal contrast between life stages may contribute to gradual generational turnover under changing climates. Environmental selection may additionally be strongest during early establishment (Petit & Hampe, 2006), while survival in older trees may increasingly depend on other biotic or stochastic factors. Together, these findings support the idea that climatic selection acts during early establishment and may progressively shape genomic structure across life stages.

In contrast to the life-stage differences observed in GEA analyses, genome-wide isolation-by-environment (IBE) was significant across regions but weaker in seedlings than in trees. However, because the three regions were also strongly differentiated by geographic distance, this difference in IBE likely reflects the combined effects of isolation by distance, demographic structure, and gene flow rather than direct evidence of adaptation. The weaker signal in seedlings may result from their closer temporal proximity to recent dispersal and cross-pollination events, whereas mortality over time may contribute to stronger structuring in adult trees.

### 4.4 Associated genes suggest selection on cellular maintenance and stress-response mechanisms

Functional enrichment analyses suggest that both genes that are either affected by allele-frequency shifts in surviving seedlings and genes associated with environmental factors, are primarily linked to fundamental cellular maintenance and stress-response processes rather than single specialized pathways. Across analyses, significant genes were enriched for functions related to intracellular transport, cellular integrity, metabolism, and physiological homeostasis, indicating that adaptation during early life stages may largely depend on maintaining cellular function under stressful environmental conditions. However, GO term enrichment analyses should generally be interpreted with caution, as results depend on annotation completeness and the choice of background gene sets.

In the allele frequency comparison between trees and surviving seedlings, the GO term GTP binding is enriched among outlier loci. This signal is absent in the comparison of trees and non-surviving seedlings, suggesting that these functions are linked to successful establishment. GTP binding proteins are found in all eukaryotes and function as molecular switches that cycle between “active” and “inactive” states. In plants they regulate the cross-talk with plant hormones and other signals, the regulation of organogenesis (leaf, root, and embryo), polarized cell growth, cell division, and the involvement in various stress and defense responses (Ma, 2007). The associated genes encode proteins involved in intracellular trafficking, cytoskeletal organization, and protein synthesis, including stress-responsive proteins such as Rab7, elongation factor Tu (EF-Tu), and β-tubulin (tubulin beta-4). Rab7 is implicated in vesicle trafficking and stress responses, EF-Tu accumulates under heat stress, and β-tubulin isoforms have been linked to cellular responses to salinity and other abiotic stresses (Agarwal et al., 2009; Chun et al., 2021; Fu et al., 2012; Yan et al., 2024; Yu et al., 2015).

Functional enrichment analyses (ORA) of genotype–environment associations revealed pronounced life stage–specific differences. Seedlings showed a broader range of biological processes related to cellular homeostasis, intracellular transport, stress responses, signaling, and primary metabolism, whereas trees exhibited fewer and functionally less diverse enrichments, mainly associated with lipid and sterol metabolism, which also play important roles in plant stress responses (Du et al., 2022; Zhao et al., 2025). This contrast was particularly pronounced for maximum air temperature, where enrichment was restricted to seedlings, suggesting that cellular maintenance and physiological flexibility may be especially important during early establishment under heat stress (Zhu, 2016). Water-related variables showed more heterogeneous patterns, but functional responses remained consistently broader in seedlings.

Similar patterns were observed in the supplementary GSEA analyses, which likewise indicated broader and more diverse functional responses in seedlings than in trees. Together, these findings are consistent with stronger abiotic filtering during seedling establishment, while survival and performance in older trees may increasingly reflect additional ecological processes such as competition, biotic interactions, and mortality.

To assess the influence of seedling establishment-year conditions, we repeated the GEA using 2021 environmental data for the Swabian Alb. Although outlier patterns between life stages were less distinct than in the main analyses, ORA again revealed broader and more functionally diverse enrichments in seedlings than in trees across environmental variables. Seedlings were primarily enriched for functions related to cellular organization, protein turnover, and sterol and lipid metabolism, whereas trees showed enrichments only for sterol and lipid metabolism and only in association with two water-related variables (rain days and soil moisture). Supplementary GSEA analyses additionally revealed stronger enrichment of chitin-related processes in both life stages, suggesting an increased importance of defense-related pathways (Vaghela et al., 2022) under the relatively cold and wet conditions of 2021. Together, these patterns suggest that reduced drought stress in 2021 may have shifted associations away from drought-related responses toward broader physiological and defense-related processes.

The patterns we found are consistent with recent findings in European beech showing that younger cohorts are enriched for pathways related to cellular integrity and homeostasis under increasing abiotic stress, whereas older cohorts are more strongly associated with processes linked to biotic interactions (Eberhardt et al., 2026). The broad functional diversity of associated genes further suggests that climate-related responses in beech may involve multiple biological pathways rather than a small number of major loci, consistent with expectations for polygenic adaptation in forest trees (Csilléry et al., 2018; Yeaman, 2015). Together, our results suggest that responses during early establishment may rely less on major shifts in specialized pathways and more on subtle changes affecting broad cellular maintenance and stress-response mechanisms. Early-life selection may therefore favor genotypes capable of maintaining intracellular function and structural integrity under stressful environmental conditions.

### 4.5 Limitations and future directions

While lcWGS enables genome-wide analyses across many individuals, the low sequencing depth required genotype imputation to obtain datasets suitable for stringent downstream analyses. Nevertheless, imputation allowed the recovery of biologically meaningful population structure. At the same time, interpretation of these signals remains constrained by the current single-reference framework, as the Bhaga genome (Mishra et al., 2022) does not allow to capture the entire genomic diversity now emerging from beech pangenome studies. Thus, aside from more sequencing depth, future studies should establish a pangenome perspective. However, pangenome resources for European beech and compatible analytical workflows are still under development.

The inference of adaptive processes is limited not only by the current availability of genomic resources but also by methodological challenges. Genotype–environment association approaches remain subject to several limitations, including the confounding effects of population structure and uncertainty in environmental predictors. We partly accounted for neutral structure by incorporating latent factors, thereby reducing spurious associations, and further controlled for false positives by repeating analyses with randomized geographic coordinates (Lazic et al., 2024), which yielded no significant outliers, supporting the presence of non-random associations between environment and genotypes. In addition, our use of high-resolution, plot-level environmental data improves spatial accuracy; however, such in situ measurements typically represent short-term conditions and may not fully capture the long-term selective environments experienced over evolutionary timescales (Dauphin et al., 2023). As our aim was to assess adaptation to current environmental conditions rather than reconstruct historical selection, current environmental measurements were appropriate for addressing our research question.

Interpretation of the observed allele-frequency shifts is constrained by differences in seedling age among regions. Because the strongest signals were detected in two-year-old seedlings from the Swabian Alb, it remains difficult to disentangle the effects of cumulative selection over time from region-specific environmental conditions. Future studies, sampling different age classes across regions and tracking successive cohorts through time could help resolve these effects.

## 5 Conclusion

In this study, we provide evidence that contemporary climate is already shaping genomic variation in European beech, with seedlings exhibiting stronger and more functionally diverse genotype–environment associations and localized allele-frequency shifts linked to survival during early establishment. At the same time, high gene flow may help maintain standing genetic variation and further facilitate adaptive responses. These findings highlight early life stages as a critical period of selection, during which environmental filtering can rapidly alter genetic composition. Together, our results suggest that adaptive responses to climate change are already ongoing, although their extent and long-term consequences remain uncertain.

Notably, the strongest allele-frequency shifts were detected in two-year-old seedlings, suggesting that selection events may accumulate during early development. Resolving how these signals change across seedling age classes represents an important direction for future research. It remains unclear when the strongest selection pulses occur during early development and how extreme environmental conditions influence their intensity and timing. Addressing whether the genetic variants retained under these dynamics confer resilience to future extremes (Coleman & Wernberg, 2020), or instead reflect responses to the most immediate stressors, will be critical for understanding the adaptive potential of long-lived species under rapid climate change.

## Acknowledgments

We thank the managers of the three Exploratories, Julia Bass, Max Müller, Anna K. Franke, Robert Künast, Franca Marian, Melissa Jüds and all former managers for their work in maintaining the plot and project infrastructure; Victoria Grießmeier for giving support through the central office, Andreas Ostrowski for managing the central database, and Markus Fischer, Eduard Linsenmair, Dominik Hessenmöller, Daniel Prati, Ingo Schöning, François Buscot, Ernst-Detlef Schulze, Wolfgang W. Weisser and Elisabeth Kalko for their role in setting up the Biodiversity Exploratories project. We thank the administration of the Hainich national park, the UNESCO Biosphere Reserve Swabian Alb and the UNESCO Biosphere Reserve Schorfheide-Chorin as well as all landowners for the excellent collaboration. The work has been (partly) funded by the DFG Priority Program 1374 "Biodiversity-Exploratories" (DFG-Refno.). Field work permits were issued by the responsible state environmental offices of Baden-Württemberg, Thüringen, and Brandenburg.

We acknowledge the support by the High Performance and Cloud Computing Group at the Zentrum für Datenverarbeitung of the University of Tübingen, the state of Baden-Württemberg through bwHPC and the German Research Foundation (DFG) for funding under "Project number 455787709" (bwForCluster BinAC 2).

We further thank Marco Göttig and Silke Fröhlich for technical assistance in the lab and Linni Dahlhausen, Jason Drescher and Maia Schillmann for assistance in sampling in the field, and the AForGen consortium for discussions on the topic.

## Data Availability

This work is based on data elaborated by the project TREEvolution of the Biodiversity Exploratories program (DFG Priority Program 1374). The datasets (ID 32454) are publicly available in the Biodiversity Exploratories Information System (http://doi.org/10.17616/R32P9Q).

Raw sequencing data is deposited in the NCBI BioProject database under accession number PRJEB121531.

Code used for the analyses is available as follows: https://gitlab.uni-marburg.de/fb17/ag-opgenoorth/treevolution-fagus-sylvatica.git

## Author contributions

ML conducted fieldwork, curated data, performed statistical analyses, prepared visualizations, and wrote the original manuscript. MS and CL contributed to fieldwork, provided methodological and statistical support, and co-supervised the project. MS also provided bioinformatical support and created all illustrations. MS, CL, LO, and KH acquired funding, conceptualized the study, supervised its implementation, and reviewed and edited the manuscript. All authors read and approved of the final manuscript.

## References

Agarwal, P., Reddy, M. K., Sopory, S. K., & Agarwal, P. K. (2009). Plant rabs: Characterization, functional diversity, and role in stress tolerance. Plant Molecular Biology Reporter, 27(4), 417–430. 10.1007/s11105-009-0100-9

Barghi, N., Hermisson, J., & Schlötterer, C. (2020). Polygenic adaptation: A unifying framework to understand positive selection. Nature Reviews Genetics, 21(12), 769–781. 10.1038/s41576-020-0250-z

Benito Garzón, M., Alía, R., Robson, T. M., & Zavala, M. A. (2011). Intra-specific variability and plasticity influence potential tree species distributions under climate change. Global Ecology and Biogeography, 20(5), 766–778. 10.1111/j.1466-8238.2010.00646.x

Benito Garzón, M., Robson, T. M., & Hampe, A. (2019). ΔTraitSDMs: Species distribution models that account for local adaptation and phenotypic plasticity. New Phytologist, 222(4), 1757–1765. 10.1111/nph.15716

BMEL (2023): Ergebnisse der Waldzustandserhebung 2023. Herausgegeben vom Bundesministerium für Ernährung und Landwirtschaft. Verfügbar unter: BMLEHWaldzustandserhebung 2023 [abgerufen am: 09.07.2026].

Booker, T. R., Yeaman, S., Whiting, J. R., & Whitlock, M. C. (2024). The WZA: A window-based method for characterizing genotype–environment associations. Molecular Ecology Resources, 24(2), e13768. 10.1111/1755-0998.13768

Browne, L., Wright, J. W., Fitz-Gibbon, S., Gugger, P. F., & Sork, V. L. (2019). Adaptational lag to temperature in valley oak (Quercus lobata) can be mitigated by genome-informed assisted gene flow. Proceedings of the National Academy of Sciences, 116(50), 25179– 25185. 10.1073/pnas.1908771116

Browning, B. L., Tian, X., Zhou, Y., & Browning, S. R. (2021). Fast two-stage phasing of large-scale sequence data. The American Journal of Human Genetics, 108(10), 1880–1890. 10.1016/j.ajhg.2021.08.005

Browning, B. L., Zhou, Y., & Browning, S. R. (2018). A one-penny imputed genome from next-generation reference panels. The American Journal of Human Genetics, 103(3), 338–348. 10.1016/j.ajhg.2018.07.015

Bruegmann, T., Fladung, M., & Schroeder, H. (2022). Flexible DNA isolation procedure for different tree species as a convenient lab routine. Silvae Genetica, 71(1), 20–30. 10.2478/sg-2022-0003

Caccianiga, M., & Compostella, C. (2012). Growth forms and age estimation of treeline species. Trees, 26(2), 331–342. 10.1007/s00468-011-0595-1

Caye, K., Jumentier, B., Lepeule, J., & François, O. (2019). LFMM 2: Fast and accurate inference of gene-environment associations in genome-wide studies. Molecular Biology and Evolution, 36(4), 852–860. 10.1093/molbev/msz008

Chen, Y., Chen, Y., Shi, C., Huang, Z., Zhang, Y., Li, S., Li, Y., Ye, J., Yu, C., Li, Z., Zhang, X., Wang, J., Yang, H., Fang, L., & Chen, Q. (2018). SOAPnuke: A MapReduce acceleration-supported software for integrated quality control and preprocessing of high-throughput sequencing data. GigaScience, 7(1), gix120. 10.1093/gigascience/gix120

Chun, H. J., Baek, D., Jin, B. J., Cho, H. M., Park, M. S., Lee, S. H., Lim, L. H., Cha, Y. J., Bae, D.-W., Kim, S. T., Yun, D.-J., & Kim, M. C. (2021). Microtubule dynamics plays a vital role in plant adaptation and tolerance to salt stress. International Journal of Molecular Sciences, 22(11), 5957. 10.3390/ijms22115957

Coleman, M. A., & Wernberg, T. (2020). The silver lining of extreme events. Trends in Ecology & Evolution, 35(12), 1065–1067. 10.1016/j.tree.2020.08.013

Collet, C., & Le Moguedec, G. (2007). Individual seedling mortality as a function of size, growth and competition in naturally regenerated beech seedlings. Forestry: An International Journal of Forest Research, 80(4), 359–370. 10.1093/forestry/cpm016

Comps, B., Gömöry, D., Letouzey, J., Thiébaut, B., & Petit, R. J. (2001). Diverging trends between heterozygosity and allelic richness during postglacial colonization in the European beech. Genetics, 157(1), 389–397. 10.1093/genetics/157.1.389

Csilléry, K., Rodríguez-Verdugo, A., Rellstab, C., & Guillaume, F. (2018). Detecting the genomic signal of polygenic adaptation and the role of epistasis in evolution. Molecular Ecology, 27(3), 606–612. 10.1111/mec.14499

Danecek, P., Auton, A., Abecasis, G., Albers, C. A., Banks, E., DePristo, M. A., Handsaker, R. E., Lunter, G., Marth, G. T., Sherry, S. T., McVean, G., Durbin, R., & 1000 Genomes Project Analysis Group. (2011). The variant call format and VCFtools. Bioinformatics, 27(15), 2156–2158. 10.1093/bioinformatics/btr330

Danecek, P., Bonfield, J. K., Liddle, J., Marshall, J., Ohan, V., Pollard, M. O., Whitwham, A., Keane, T., McCarthy, S. A., Davies, R. M., & Li, H. (2021). Twelve years of SAMtools and BCFtools. GigaScience, 10(2), giab008. 10.1093/gigascience/giab008

Dauphin, B., Rellstab, C., Schmid, M., Zoller, S., Karger, D. N., Brodbeck, S., Guillaume, F., & Gugerli, F. (2021). Genomic vulnerability to rapid climate warming in a tree species with a long generation time. Global Change Biology, 27(6), Article 6. 10.1111/gcb.15469

Dauphin, B., Rellstab, C., Wüest, R. O., Karger, D. N., Holderegger, R., Gugerli, F., & Manel, S. (2023). Re-thinking the environment in landscape genomics. Trends in Ecology & Evolution, 38(3), 261–274. 10.1016/j.tree.2022.10.010

De La Torre, A. R., Wilhite, B., & Neale, D. B. (2019). Environmental genome-wide association reveals climate adaptation is shaped by subtle to moderate allele frequency shifts in loblolly pine. Genome Biology and Evolution, 11(10), 2976–2989. 10.1093/gbe/evz220

Du, Y., Fu, X., Chu, Y., Wu, P., Liu, Y., Ma, L., Tian, H., & Zhu, B. (2022). Biosynthesis and the roles of plant sterols in development and stress responses. International Journal of Molecular Sciences, 23(4), 2332. 10.3390/ijms23042332

Dyderski, M. K., Paź, S., Frelich, L. E., & Jagodziński, A. M. (2018). How much does climate change threaten European forest tree species distributions? Global Change Biology, 24(3), 1150–1163. 10.1111/gcb.13925

Eberhardt, L., Reuss, F., Blázquez, M. E. N., Hetzer, J., Feldmeyer, B., & Pfenninger, M. (2026). Climate change intensifies rapid genomic selection beyond the ancestral niche of Fagus sylvatica (p. 2026.03.09.710448). bioRxiv. 10.64898/2026.03.09.710448

Fischer, M., Bossdorf, O., Gockel, S., Hänsel, F., Hemp, A., Hessenmöller, D., Korte, G., Nieschulze, J., Pfeiffer, S., Prati, D., Renner, S., Schöning, I., Schumacher, U., Wells, K., Buscot, F., Kalko, E. K. V., Linsenmair, K. E., Schulze, E.-D., & Weisser, W. W. (2010). Implementing large-scale and long-term functional biodiversity research: The Biodiversity Exploratories. Basic and Applied Ecology, 11(6), 473–485. 10.1016/j.baae.2010.07.009

Foest, J. J., Bogdziewicz, M., Pesendorfer, M. B., Ascoli, D., Cutini, A., Nussbaumer, A., Verstraeten, A., Beudert, B., Chianucci, F., Mezzavilla, F., Gratzer, G., Kunstler, G., Meesenburg, H., Wagner, M., Mund, M., Cools, N., Vacek, S., Schmidt, W., Vacek, Z., & Hacket-Pain, A. (2024). Widespread breakdown in masting in European beech due to rising summer temperatures. Global Change Biology, 30(5), e17307. 10.1111/gcb.17307

Franks, S. J., & Hoffmann, A. A. (2012). Genetics of climate change adaptation. Annual Review of Genetics, 46(Volume 46, 2012), 185–208. 10.1146/annurev-genet-110711-155511

Frichot, E., & François, O. (2015). LEA: An R package for landscape and ecological association studies. Methods in Ecology and Evolution, 6(8), 925–929. 10.1111/2041-210X.12382

Fu, J., Momčilović, I., & Prasad, P. V. V. (2012). Roles of protein synthesis elongation factor EF-Tu in heat tolerance in plants. Journal of Botany, 2012(1), 835836. 10.1155/2012/835836

Gárate-Escamilla, H., Hampe, A., Vizcaíno-Palomar, N., Robson, T. M., & Benito Garzón, M. (2019). Range-wide variation in local adaptation and phenotypic plasticity of fitness-related traits in Fagus sylvatica and their implications under climate change. Global Ecology and Biogeography, 28(9), 1336–1350. 10.1111/geb.12936

Grossnickle, S. C. (2012). Why seedlings survive: Influence of plant attributes. New Forests, 43(5), 711–738. 10.1007/s11056-012-9336-6

Hanewinkel, M., Cullmann, D. A., Schelhaas, M.-J., Nabuurs, G.-J., & Zimmermann, N. E. (2013). Climate change may cause severe loss in the economic value of European forest land. Nature Climate Change, 3(3), 203–207. 10.1038/nclimate1687

Hegerl, G. C., Brönnimann, S., Cowan, T., Friedman, A. R., Hawkins, E., Iles, C., Müller, W., Schurer, A., & Undorf, S. (2019). Causes of climate change over the historical record. Environmental Research Letters, 14(12), 123006. 10.1088/1748-9326/ab4557

Hermisson, J., & Pennings, P. S. (2005). Soft sweeps: Molecular population genetics of adaptation from standing genetic variation. Genetics, 169(4), 2335–2352. 10.1534/genetics.104.036947

Hoban, S., Kelley, J. L., Lotterhos, K. E., Antolin, M. F., Bradburd, G., Lowry, D. B., Poss, M. L., Reed, L. K., Storfer, A., & Whitlock, M. C. (2016). Finding the genomic basis of local adaptation: Pitfalls, practical solutions, and future directions. The American Naturalist, 188(4), 379–397. 10.1086/688018

Hohenlohe, P. A., Phillips, P. C., & Cresko, W. A. (2010). Using population genomics to detect selection in natural populations: Key concepts and methodological considerations. International Journal of Plant Sciences, 171(9), 1059–1071. 10.1086/656306

Höhn, M., Major, E., Avdagić, A., Bielak, K., Bosela, M., Coll, L., Dinca, L., Giammarchi, F., Ibrahimspahić, A., Mataruga, M., Pach, M., Uhl, E., Zlatanov, T., Cseke, K., Kovács, Zs., Palla, B., Ladányi, M., & Heinze, B. (2021). Local characteristics of the standing genetic diversity of European beech with high within-region differentiation at the eastern part of the range. Canadian Journal of Forest Research, 51(12), 1791–1798. 10.1139/cjfr-2020-0413

Kremer, A., Chen, J., & Lascoux, M. (2025). ‘Chimes of resilience’: What makes forest trees genetically resilient? New Phytologist, 246(5), 1934–1951. 10.1111/nph.70108

Lazic, D., Geßner, C., Liepe, K. J., Lesur-Kupin, I., Mader, M., Blanc-Jolivet, C., Gömöry, D., Liesebach, M., González-Martínez, S. C., Fladung, M., Degen, B., & Müller, N. A. (2024). Genomic variation of European beech reveals signals of local adaptation despite high levels of phenotypic plasticity. Nature Communications, 15(1), 8553. 10.1038/s41467-024-52933-y

Leuschner, C. (2020). Drought response of European beech (Fagus sylvatica L.)—A review. Perspectives in Plant Ecology, Evolution and Systematics, 47, 125576. 10.1016/j.ppees.2020.125576

Lotterhos, K. E. (2023). The paradox of adaptive trait clines with nonclinal patterns in the underlying genes. Proceedings of the National Academy of Sciences, 120(12), e2220313120. 10.1073/pnas.2220313120

Lotterhos, K. E., & Whitlock, M. C. (2015). The relative power of genome scans to detect local adaptation depends on sampling design and statistical method. Molecular Ecology, 24(5), 1031–1046. 10.1111/mec.13100

Lou, R. N., Jacobs, A., Wilder, A. P., & Therkildsen, N. O. (2021). A beginner’s guide to low-coverage whole genome sequencing for population genomics. Molecular Ecology, 30(23), 5966–5993. 10.1111/mec.16077

Ma, Q.-H. (2007). Small GTP-binding proteins and their functions in plants. Journal of Plant Growth Regulation, 26(4), 369–388. 10.1007/s00344-007-9022-7

Martinez del Castillo, E., Zang, C. S., Buras, A., Hacket-Pain, A., Esper, J., Serrano-Notivoli, R., Hartl, C., Weigel, R., Klesse, S., Resco de Dios, V., Scharnweber, T., Dorado-Liñán, I., van der Maaten-Theunissen, M., van der Maaten, E., Jump, A., Mikac, S., Banzragch, B.-E., Beck, W., Cavin, L., … de Luis, M. (2022). Climate-change-driven growth decline of European beech forests. Communications Biology, 5(1), 163. 10.1038/s42003-022-03107-3

McVean, G. (2007). The structure of linkage disequilibrium around a selective sweep. Genetics, 175(3), 1395–1406. 10.1534/genetics.106.062828

Metheringham, C. L., Plumb, W. J., Flynn, W. R. M., Stocks, J. J., Kelly, L. J., Nemesio Gorriz, M., Grieve, S. W. D., Moat, J., Lines, E. R., Buggs, R. J. A., & Nichols, R. A. (2025). Rapid polygenic adaptation in a wild population of ash trees under a novel fungal epidemic. Science, 388(6754), 1422–1425. 10.1126/science.adp2990

Miranda, J. C., Calderaro, C., Cocozza, C., Lasserre, B., Tognetti, R., & von Arx, G. (2022). Wood anatomical responses of European beech to elevation, land use change, and climate variability in the central apennines, Italy. Frontiers in Plant Science, 13. 10.3389/fpls.2022.855741

Mishra, B., Ulaszewski, B., Meger, J., Aury, J.-M., Bodénès, C., Lesur-Kupin, I., Pfenninger, M., Da Silva, C., Gupta, D. K., Guichoux, E., Heer, K., Lalanne, C., Labadie, K., Opgenoorth, L., Ploch, S., Le Provost, G., Salse, J., Scotti, I., Wötzel, S., … Thines, M. (2022). A chromosome-level genome assembly of the European beech (Fagus sylvatica) reveals anomalies for organelle DNA integration, repeat content and distribution of SNPs. Frontiers in Genetics, 12. https://www.frontiersin.org/articles/10.3389/fgene.2021.691058

Ostrowski, A., Nieschulze, J., Schulze, E.-D., & König-Ries, B. (2023). *Coordinates and Inventory Overview of all Grid Plots (GPs)* (Version 5) [Dataset]. Biodiversity Exploratories Information System. www.bexis.uni-jena.de

Pasaniuc, B., Rohland, N., McLaren, P. J., Garimella, K., Zaitlen, N., Li, H., Gupta, N., Neale, B. M., Daly, M. J., Sklar, P., Sullivan, P. F., Bergen, S., Moran, J. L., Hultman, C. M., Lichtenstein, P., Magnusson, P., Purcell, S. M., Haas, D. W., Liang, L., … Price, A. L. (2012). Extremely low-coverage sequencing and imputation increases power for genome-wide association studies. Nature Genetics, 44(6), 631–635. 10.1038/ng.2283

Patterson, N., Moorjani, P., Luo, Y., Mallick, S., Rohland, N., Zhan, Y., Genschoreck, T., Webster, T., & Reich, D. (2012). Ancient admixture in human history. Genetics, 192(3), 1065–1093. 10.1534/genetics.112.145037

Patterson, N., Price, A. L., & Reich, D. (2006). Population structure and eigenanalysis. PLoS Genetics, 2(12), e190. 10.1371/journal.pgen.0020190

Pennings, P. S., & Hermisson, J. (2006). Soft sweeps II - Molecular population genetics of adaptation from recurrent mutation or migration. Molecular Biology and Evolution, 23(5), 1076–1084. 10.1093/molbev/msj117

Petit, R. J., & Hampe, A. (2006). Some evolutionary consequences of being a tree. Annual Review of Ecology, Evolution, and Systematics, 37(1), 187–214. 10.1146/annurev.ecolsys.37.091305.110215

Petr, M., Vernot, B., & Kelso, J. (2019). admixr—R package for reproducible analyses using ADMIXTOOLS. Bioinformatics, 35(17), 3194–3195. 10.1093/bioinformatics/btz030

Pew, J., Muir, P. H., Wang, J., & Frasier, T. R. (2015). related: An R package for analysing pairwise relatedness from codominant molecular markers. Molecular Ecology Resources, 15(3), 557–561. 10.1111/1755-0998.12323

Pfenninger, M., Reuss, F., Kiebler, A., Schönnenbeck, P., Caliendo, C., Gerber, S., Cocchiararo, B., Reuter, S., Blüthgen, N., Mody, K., Mishra, B., Bálint, M., Thines, M., & Feldmeyer, B. (2021). Genomic basis for drought resistance in European beech forests threatened by climate change. eLife, 10, e65532. 10.7554/eLife.65532

Plomion, C., Aury, J.-M., Amselem, J., Leroy, T., Murat, F., Duplessis, S., Faye, S., Francillonne, N., Labadie, K., Le Provost, G., Lesur, I., Bartholomé, J., Faivre-Rampant, P., Kohler, A., Leplé, J.-C., Chantret, N., Chen, J., Diévart, A., Alaeitabar, T., … Salse, J. (2018). Oak genome reveals facets of long lifespan. Nature Plants, 4(7), 440–452. 10.1038/s41477-018-0172-3

Pluess, A. R., Frank, A., Heiri, C., Lalagüe, H., Vendramin, G. G., & Oddou-Muratorio, S. (2016). Genome–environment association study suggests local adaptation to climate at the regional scale in Fagus sylvatica. New Phytologist, 210(2), 589–601. 10.1111/nph.13809

Postolache, D., Oddou-Muratorio, S., Vajana, E., Bagnoli, F., Guichoux, E., Hampe, A., Le Provost, G., Lesur, I., Popescu, F., Scotti, I., Piotti, A., & Vendramin, G. G. (2021). Genetic signatures of divergent selection in European beech (Fagus sylvatica L.) are associated with the variation in temperature and precipitation across its distribution range. Molecular Ecology, 30(20), 5029–5047. 10.1111/mec.16115

Rohner, B., Kumar, S., Liechti, K., Gessler, A., & Ferretti, M. (2021). Tree vitality indicators revealed a rapid response of beech forests to the 2018 drought. Ecological Indicators, 106903. 10.1016/j.ecolind.2020.106903

Rubinacci, S., Ribeiro, D. M., Hofmeister, R. J., & Delaneau, O. (2021). Efficient phasing and imputation of low-coverage sequencing data using large reference panels. Nature Genetics, 53(1), 120–126. 10.1038/s41588-020-00756-0

Schuldt, B., Buras, A., Arend, M., Vitasse, Y., Beierkuhnlein, C., Damm, A., Gharun, M., Grams, T. E. E., Hauck, M., Hajek, P., Hartmann, H., Hiltbrunner, E., Hoch, G., Holloway-Phillips, M., Körner, C., Larysch, E., Lübbe, T., Nelson, D. B., Rammig, A., … Kahmen, A. (2020). A first assessment of the impact of the extreme 2018 summer drought on Central European forests. Basic and Applied Ecology, 45, 86–103. 10.1016/j.baae.2020.04.003

Senf, C., & Seidl, R. (2021). Persistent impacts of the 2018 drought on forest disturbance regimes in Europe. Biogeosciences, 18(18), 5223–5230. 10.5194/bg-18-5223-2021

Slavov, G. T., Macaya-Sanz, D., DiFazio, S. P., & Howe, G. T. (2025). Population structure limits the use of genomic data for predicting phenotypes and managing genetic resources in forest trees. Proceedings of the National Academy of Sciences, 122(26), e2425691122. 10.1073/pnas.2425691122

Stephan, W. (2019). Selective Sweeps. Genetics, 211(1), 5–13. 10.1534/genetics.118.301319

Stephan, Wolfgang & Hörger, Anja. (2019). Molekulare Populationsgenetik. Springer. 10.1007/978-3-662-59428-5

Sui, J., Li, G., Chen, G., Zhao, C., Kong, X., Hou, X., Qiao, L., & Wang, J. (2017). RNA-seq analysis reveals the role of a small GTP-binding protein, Rab7, in regulating clathrin-mediated endocytosis and salinity-stress resistance in peanut. Plant Biotechnology Reports, 11(1), 43–52. 10.1007/s11816-017-0428-9

Vaghela, B., Vashi, R., Rajput, K., & Joshi, R. (2022). Plant chitinases and their role in plant defense: A comprehensive review. Enzyme and Microbial Technology, 159, 110055. 10.1016/j.enzmictec.2022.110055

Valladares, F., Matesanz, S., Guilhaumon, F., Araújo, M. B., Balaguer, L., Benito-Garzón, M., Cornwell, W., Gianoli, E., van Kleunen, M., Naya, D. E., Nicotra, A. B., Poorter, H., & Zavala, M. A. (2014). The effects of phenotypic plasticity and local adaptation on forecasts of species range shifts under climate change. Ecology Letters, 17(11), 1351– 1364. 10.1111/ele.12348

Wang, J. (2011). coancestry: A program for simulating, estimating and analysing relatedness and inbreeding coefficients. Molecular Ecology Resources, 11(1), 141–145. 10.1111/j.1755-0998.2010.02885.x

Wilczek, A. M., Cooper, M. D., Korves, T. M., & Schmitt, J. (2014). Lagging adaptation to warming climate in Arabidopsis thaliana. Proceedings of the National Academy of Sciences, 111(22), 7906–7913. 10.1073/pnas.1406314111

Wöllauer, S., Zeuss, D., Hänsel, F., & Nauss, T. (2021). TubeDB: An on-demand processing database system for climate station data. Computers & Geosciences, 146, 104641. 10.1016/j.cageo.2020.104641

Yan, M., Dong, Z., Pan, T., Li, L., Zhou, Z., Li, W., Ke, Z., Feng, Z., & Yu, S. (2024). Systematical characterization of Rab7 gene family in Gossypium and potential functions of GhRab7B3-A gene in drought tolerance. BMC Genomics, 25(1), 1023. 10.1186/s12864-024-10930-x

Yeaman, S. (2015). Local Adaptation by Alleles of Small Effect. The American Naturalist, 186(S1), S74–S89. 10.1086/682405

Yu, J., Qiu, H., Liu, X., Wang, M., Gao, Y., Chory, J., & Tao, Y. (2015). Characterization of tub4P287L, a β-tubulin mutant, revealed new aspects of microtubule regulation in shade. Journal of Integrative Plant Biology, 57(9), 757–769. 10.1111/jipb.12363

Zhao, Y., Cao, R., Li, J., Xu, Y., Zhou, L., & Ye, Y. (2025). Lipid droplets in plants: Turnover and stress responses. Frontiers in Plant Science, 16. 10.3389/fpls.2025.1625830

Zhu, J.-K. (2016). Abiotic Stress Signaling and Responses in Plants. Cell, 167(2), 313–324. 10.1016/j.cell.2016.08.029

