## Supplementary Information for "Early establishment acts as a selective filter shaping climate-associated genomic variation in European beech"

### Supplements

#### Supplementary Methods

##### ***Supplementary Methods S1 | DNA extraction***

We extracted genomic DNA from 20–50 mg silica-dried leaf tissue homogenized with a Retsch mill (MM 301) following the ATMAB protocol of Bruegmann et al. (2022) with minor modifications. Samples were centrifuged at 13,000 rpm, and DNA was precipitated by overnight incubation with cold isopropanol at room temperature instead of 30 min at −20 °C. DNA pellets were resuspended in RNase solution (10 µg $ml^{-1}$), incubated overnight at room temperature, and RNase was inactivated at 37 °C for 30 min. DNA quantity and quality were assessed using a Qubit Fluorometer and NanoDrop spectrophotometer (Thermo Fisher Scientific), and DNA was stored at −20 °C until further processing.

##### ***SupplementaryMethods S2 | Read alignment and variant calling***

We merged FASTQ files from the two sequencing lanes of each sample and aligned reads to the chromosome-level *Fagus sylvatica* reference genome Bhaga (Mishra et al., 2022) using BWA-MEM v0.7.17 (Li, 2013). BAM files were sorted and indexed with Novosort v1.03.01 (Novocraft Technologies), and mapping statistics were calculated with SAMtools (Danecek et al., 2021). We merged all BAM files into a multi-sample BAM using Picard Tools v2.13.3 ("Picard Toolkit," 2019) and performed chromosome-wise SNP calling with SAMtools mpileup (minimum mapping quality 30, minimum base quality 20) and bcftools call v1.17 (Danecek et al., 2021). Chromosome-specific VCFs were concatenated, sorted, and reheadered to generate a final dataset comprising 1,032 individuals.

##### ***Supplementary Methods S3 | Variant filtering***

We first masked genotypes with sequencing depth <3 or genotype quality <5 as missing (bcftools +setGT; FMT/DP < 3 | FMT/GQ < 5). We removed sites with excessive heterozygosity

(>90% heterozygous genotypes; INFO/AC_Het / INFO/NS > 0.9) and retained variants with QUAL ≥40, alternate allele frequency ≥0.01 (AC/AN ≥0.01), and ≤90% missing genotypes (N_MISSING/AN ≤0.9).

For downstream analyses, we applied stricter genotype filters (--minDP 3, --minGQ 20), removed sites with mean depth >100 (--max-meanDP 100), retained only biallelic SNPs (-m2 -M2 -v snps, MAC >0), excluded variants with >20% missing genotypes (--max-missing 0.8), removed singletons (--mac 2), and excluded individuals with >60% missing data (--missing-indv, threshold 0.6).

***Supplementary Methods S4 | D-statistics***

Preparation of the outgroup dataset for D-statistics. To obtain an outgroup for ABBA–BABA (D-statistics) analyses, the Quercus robur reference genome assembly version V2_2N (Plomion et al., 2018) was aligned to the chromosome-level Fagus sylvatica Bhaga reference genome using minimap2. Genotypes for Q. robur were then inferred at the genomic positions of the filtered F. sylvatica SNP dataset using bcftools. The resulting outgroup genotypes were merged with the filtered beech SNP dataset, retaining only genomic positions for which both the ingroup and outgroup genotypes were available. The merged dataset was converted to EIGENSTRAT format using PLINK and convertf for downstream analyses with ADMIXTOOLS.

***Supplementary Methods S5 | Environmental variables for GEA***

Environmental variables used in GEA analysis included precipitation, number of rain days, relative humidity at 2 m (minimum, mean, and maximum), soil moisture at 10 cm depth, air temperature at 2 m (minimum, mean, and maximum), soil temperature at 10 cm and 20 cm depth, and the number of cool and summer days. Monthly values were aggregated to plot-level means.

#### Supplementary Tables

***Table S1 | Sampling overview***

*Forest plots measured 100 × 100 m and represented different management regimes. Sampling was conducted between May and June 2023. One leaf was collected from each seedling and 1–3 leaves from each adult tree. Adult leaves were collected using a slingshot (Bigshot, Sherrill Inc.). Seedlings were fenced and marked to enable relocation during the survival assessment in 2024. Leaf samples were dried in silica gel after removal of the central midrib. Individual positions relative to plot centers were recorded using a Haglöf Vertex Laser Geo.*

| Region | No. of Plots | No. of tree–seedling pairs | No. of individuals |
|---|---|---|---|
| ALB | 31 | 169 | 338 |
| HAI | 31 | 183 | 366 |
| SCH | 33 | 188 | 376 |
| Total | 95 | 540 | 1080 |

***Table S2 | Summary of seedling survival one year after sampling across the three study regions.***

*Seedlings were sampled in May–June 2023 and revisited in April–May 2024. Individuals recorded as dead and those not found during the resurvey were combined into a single "dead" category for all subsequent analyses. The table reports count of surviving, dead, and not-found seedlings for each region (ALB, HAI, SCH), together with total sample sizes and aggregated values across all 540 sampled seedlings.*

| Region | Surviving seedlings | Dead seedlings | Not found seedlings | Dead (combined) | Total |
|---|---|---|---|---|---|
| ALB | 151 | 16 | 2 | 18 | 169 |
| HAI | 63 | 79 | 41 | 120 | 183 |
| SCH | 59 | 77 | 52 | 129 | 188 |
| Total | 273 | 172 | 95 | 267 | 540 |

***Table S3 | Differences in morphological traits across the three locations between surviving and non-surviving seedlings with LMM***

| Trait | F | df | p |
|---|---|---|---|
| Chlorophyll content | 11.24 | 483.12 | **< 0.01** |
| Leaf area | 12.24 | 439.10 | **< 0.01** |
| Cotyledone height | 2.03 | 500.69 | 0.155 |
| Height of first.scar | 3.63 | 494.34 | 0.06 |
| Apical height | 2.58 | 508.94 | 0.11 |
| Stem diameter | 19.65 | 502.70 | **< 0.01** |
| DIFN | 4.45 | 213.42 | **< 0.05** |

***Table S4 | Differences in morphological traits across the three locations between surviving and non-surviving seedlings with GLMM***

| Trait | Model | z-value | df | p |
|---|---|---|---|---|
| Leaf thickness | Gamma | -0.21 | NA | 0.84 |
| Number of leaves | Poisson | 0.88 | NA | 0.38 |

***Table S5 | Observed and expected heterozygosity for the three study regions (ALB, HAI, SCH) separated by trees and seedlings.***

Values are based on genome-wide SNP data. Observed heterozygosity (H_O) and expected heterozygosity (H_E) show very similar levels across regions, indicating comparable genetic diversity.

| Population | Life stage | Ho_mean | He_mean | N_samples |
|---|---|---|---|---|
| ALB | Seedling | 0.292 | 0.297 | 160 |
| ALB | Tree | 0.299 | 0.297 | 157 |
| HAI | Seedling | 0.294 | 0.297 | 178 |
| HAI | Tree | 0.295 | 0.297 | 165 |
| SCH | Seedling | 0.292 | 0.297 | 176 |
| SCH | Tree | 0.296 | 0.297 | 170 |

***Table S6a | Mean genetic diversity (pi) per region and life stage is similar among the three populations ($\pi = 2.50 \times 10^{-3}$) based on the imputed dataset***

| Population | Life stage | π (genetic diversity) |
|---|---|---|
| ALB | Seedling | 0.00249 |
| ALB | Tree | 0.00251 |
| HAI | Seedling | 0.00250 |
| HAI | Tree | 0.00250 |
| SCH | Seedling | 0.00250 |
| SCH | Tree | 0.00250 |

***Table S6b*** | ***Mean genetic diversity (pi) per region and life stage is similar among the three populations ($\pi = 2.20 \times 10^{-3}$) based on the non-imputed dataset***

| Population | Life stage | π (genetic diversity) |
|---|---|---|
| ALB | Seedling | 0.00214 |
| ALB | Tree | 0.00224 |
| HAI | Seedling | 0.00230 |
| HAI | Tree | 0.00222 |
| SCH | Seedling | 0.00220 |
| SCH | Tree | 0.00230 |

***Table S7 | Mean pairwise Fst among the three locations, separated by trees and seedlings and survival status of seedlings (dead / survived) assessed one year after sampling.***

| | ALB | HAI | SCH |
|---|---|---|---|
| Dead | none | 0.00657 | 0.0111 |
| Survived | 0.00543 | 0.0113 | 0.00386 |
| Seedlings | 0.00573 | 0.00987 | 0.0122 |
| Trees | 0.00552 | 0.00777 | 0.0110 |

***Table S8 | Summary of candidate genes containing significant SNPs identified by the allele-frequency shift analysis with surviving seedlings.*** *The table lists the significant SNPs associated with each gene, the number of significant SNPs per gene, the functional annotation extracted from the Fagus sylvatica Bhaga reference genome GFF annotation (Mishra et al. 2021), and the associated Gene Ontology (GO) identifiers.*

| Gene ID | SNPs | N SNPs | GFF Description | GO IDs |
|---|---|---|---|---|
| Bhaga_10.g352 | 10_2861430; 10_2861439; 10_2861442; 10_2861451; 10_2861457; 10_2861463; 10_2861469 | 7 | XP_023898678.1 ADP/ATP carrier protein, mitochondrial-like | GO:0005471; GO:0005743; GO:0015866; GO:0015867; GO:0016021; GO:0055085 |
| Bhaga_10.g599 | 10_4948007; 10_4948013 | 2 | XP_023877142.1 tubulin beta-4 chain | GO:0000226; GO:0000278; GO:0003924; GO:0005200; GO:0005525; GO:0005737; GO:0005874 |
| Bhaga_10.g2043 | 10_16577607; 10_16577618; 10_16577621; 10_16577630; | 12 | XP_030938629.1 elongation factor TuB, chloroplastic | GO:0003746; GO:0003924; GO:0005525; |

| | | | | |
|---|---|---|---|---|
| | 10_16577642;<br>10_16577648;<br>10_16577657;<br>10_16577661;<br>10_16577663;<br>10_16577681;<br>10_16577690;<br>10_16577699 | | | GO:0005739;<br>GO:0070125 |
| Bhaga_11.g253 | 11_2317781 | 1 | KAB1205632.1ATP synthase subunit beta, mitochondrial | GO:0000275;<br>GO:0005524;<br>GO:0016887;<br>GO:0042776;<br>GO:0046933 |
| Bhaga_12.g2806 | 12_23644005;<br>12_23644007;<br>12_23644013;<br>12_23644016;<br>12_23644019;<br>12_23644034 | 6 | KAB1207895.1ADP-ribosylation factor | GO:0005524;<br>GO:0005525;<br>GO:0005737;<br>GO:0006886;<br>GO:0008935;<br>GO:0009234;<br>GO:0016192;<br>GO:0016301;<br>GO:0046854;<br>GO:0048015 |

| | | | | |
|---|---|---|---|---|
| Bhaga_2.g2933 | 2_26894512; 2_26894514; 2_26894515; 2_26894518; 2_26894521; 2_26894530; 2_26894557 | 7 | XP_023919209.1 guanine nucleotide-binding protein subunit beta-like protein | GO:0001934; GO:0005080; GO:0005634; GO:0016021; GO:0022627; GO:0043022; GO:0072344 |
| Bhaga_3.g4267 | 3_37739291 | 1 | XP_023903852.1 heat shock protein 83 | GO:0005524; GO:0005829; GO:0005886; GO:0006457; GO:0009986; GO:0032991; GO:0034605; GO:0048471; GO:0050821; GO:0051082 |
| Bhaga_4.g1879 | 4_16128642; 4_16128648 | 2 | XP_030952138.1 ras-related protein Rab7 | GO:0003924; GO:0005525; GO:0005774 |
| Bhaga_5.g3334 | 5_29565256; 5_29565262 | 2 | XP_027340522.1 adenosylhomocysteinase isoform X1 | GO:0004013; GO:0005829; GO:0006730; GO:0033353 |

| | | | | |
|---|---|---|---|---|
| Bhaga_7.g206 | 7_1750320;<br>7_1750323;<br>7_1750326;<br>7_1750329;<br>7_1750341 | 5 | XP_007041601.1 actin-100 | GO:0005524;<br>GO:0005737;<br>GO:0005856 |

***Table S9 | Number of outlier SNPs per lifestage & environmental variable for the GEA analysis with environmental variables from the growing season (March – October) 2023***

| **Environmental variable** | **No. of outlier for GEA with seedlings** | **No. of outlier for GEA with trees** |
|---|---|---|
| Maximum air temperature at 2m height [°C] | 3004 | 1431 |
| Maximum relative humidity | 434 | 4 |
| Minimum relative humidity | 133 | 41 |
| Soil moisture at 10cm depth | 2617 | 3586 |
| Precipitation | 1738 | 1964 |
| Rain days | 3171 | 3338 |

#### Supplementary Figures

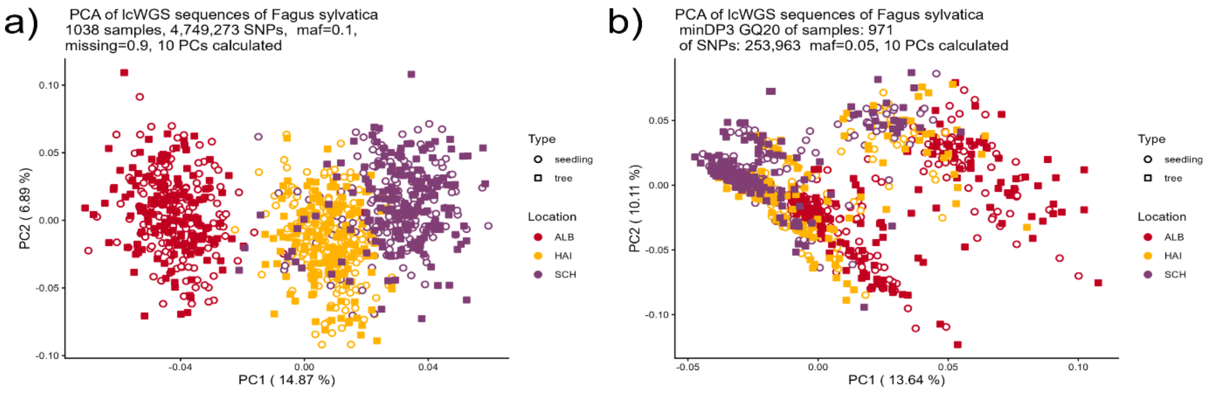


***Figure S1 | Principal Component Analysis*** *a) PCA of the raw unfiltered dataset showing clear population structure between three regions and b) PCA of the filtered VCF shows distorted population structure after stringent filtering without imputation*

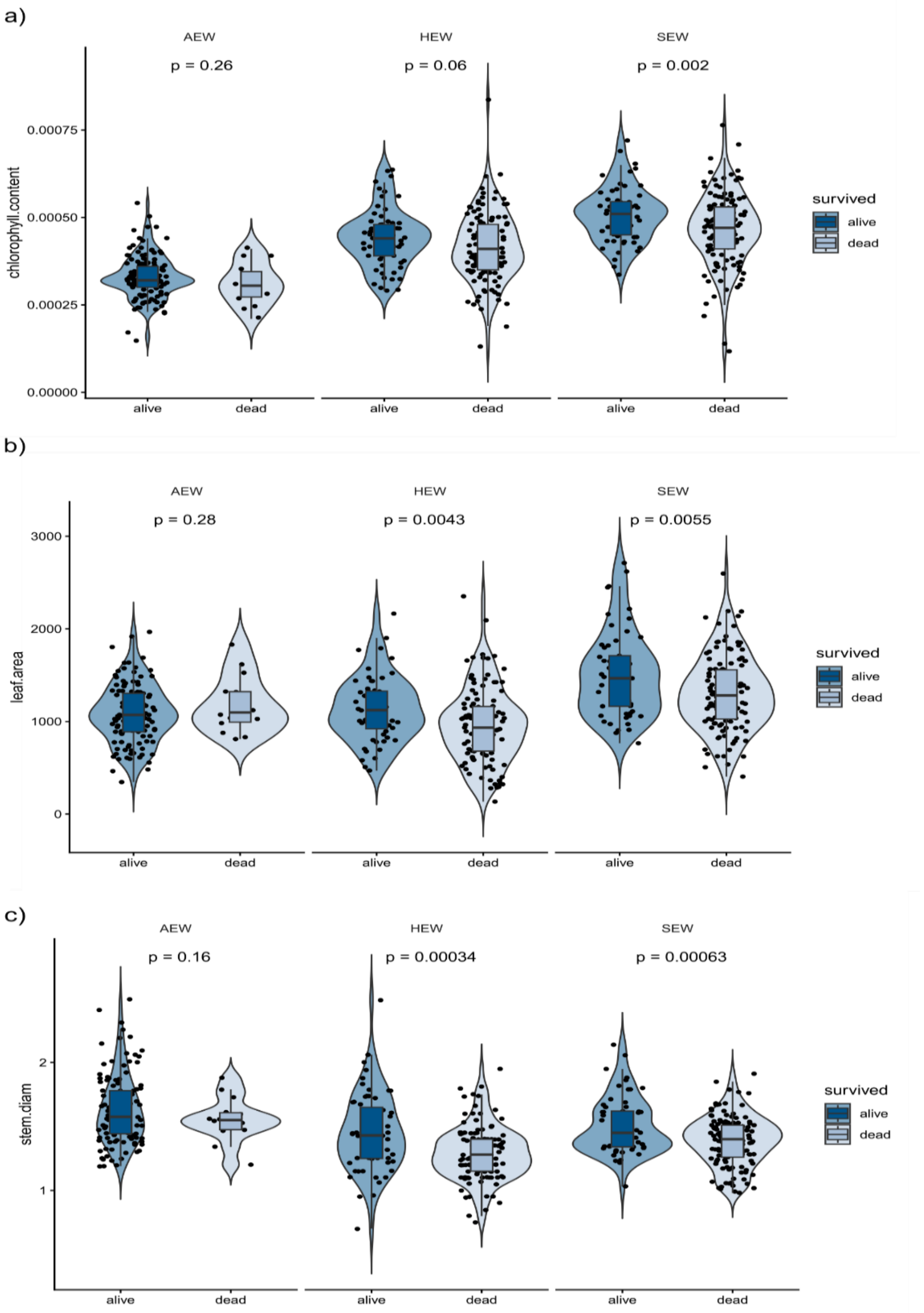


***Figure S2 | Differences in morphological traits between seedlings that survived or died after one year.*** *Morphological traits were measured at the time of sampling in 2023, before seedling survival was assessed. Survival status was determined during resampling in 2024. Panels show a) chlorophyll content, b) leaf area, and c) stem diameter for surviving and dead seedlings within each region.*

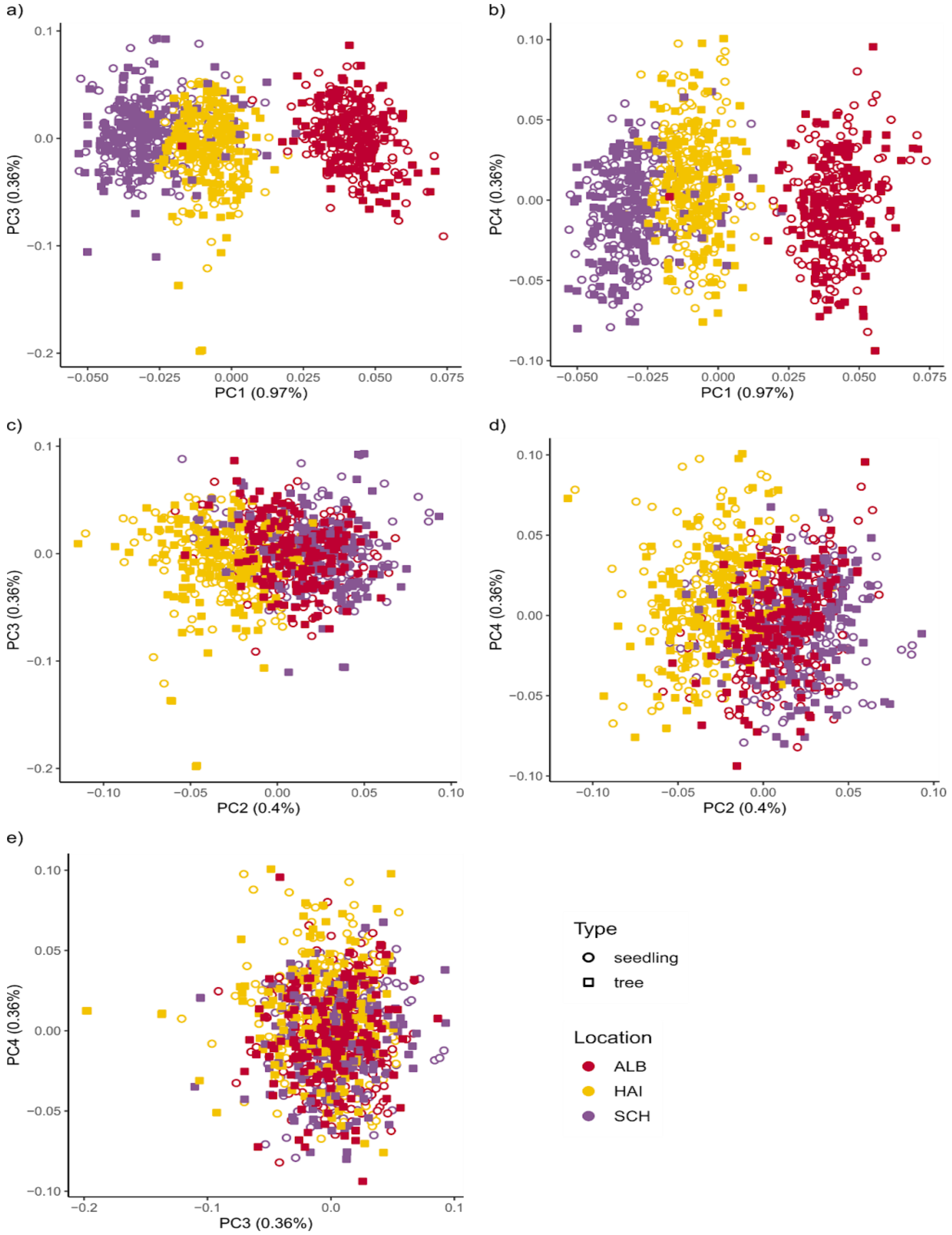


***Figure S3 | Principle component analysis (PCA) shows geographic differentiation for PC1 and PC2.***

*A – e) PCA for PC1 – PC4 of the three locations, visually separated by tree and seedlings. PC1 and PC2 show marked geographic differentiations, whereas PC3 and PC4 show no clear differentiations.*

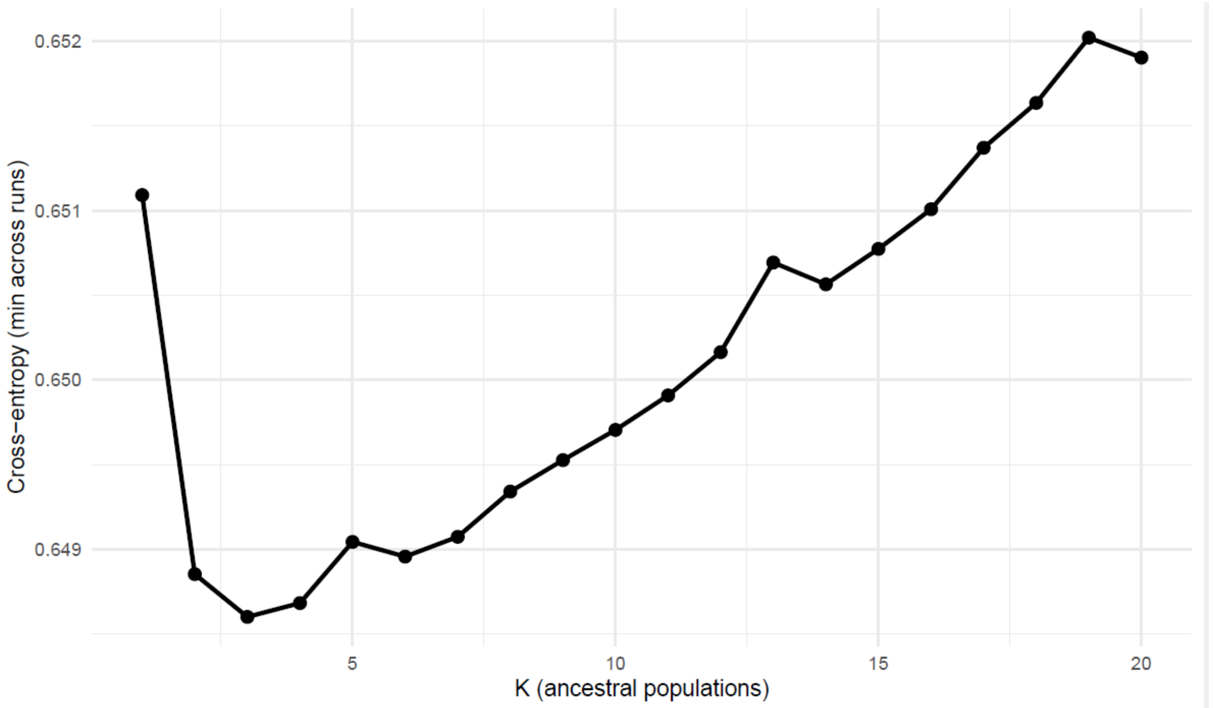


***Figure S4 | Cross-entropy of ancestry coefficients analysis***

*Cross-entropy values obtained from sparse non-negative matrix factorization (SNMF) analysis for K = 1 – 20 genetic clusters. Three major ancestral genetic clusters are indicated by the lowest K at 3 calculated from independent genome-wide variants.*

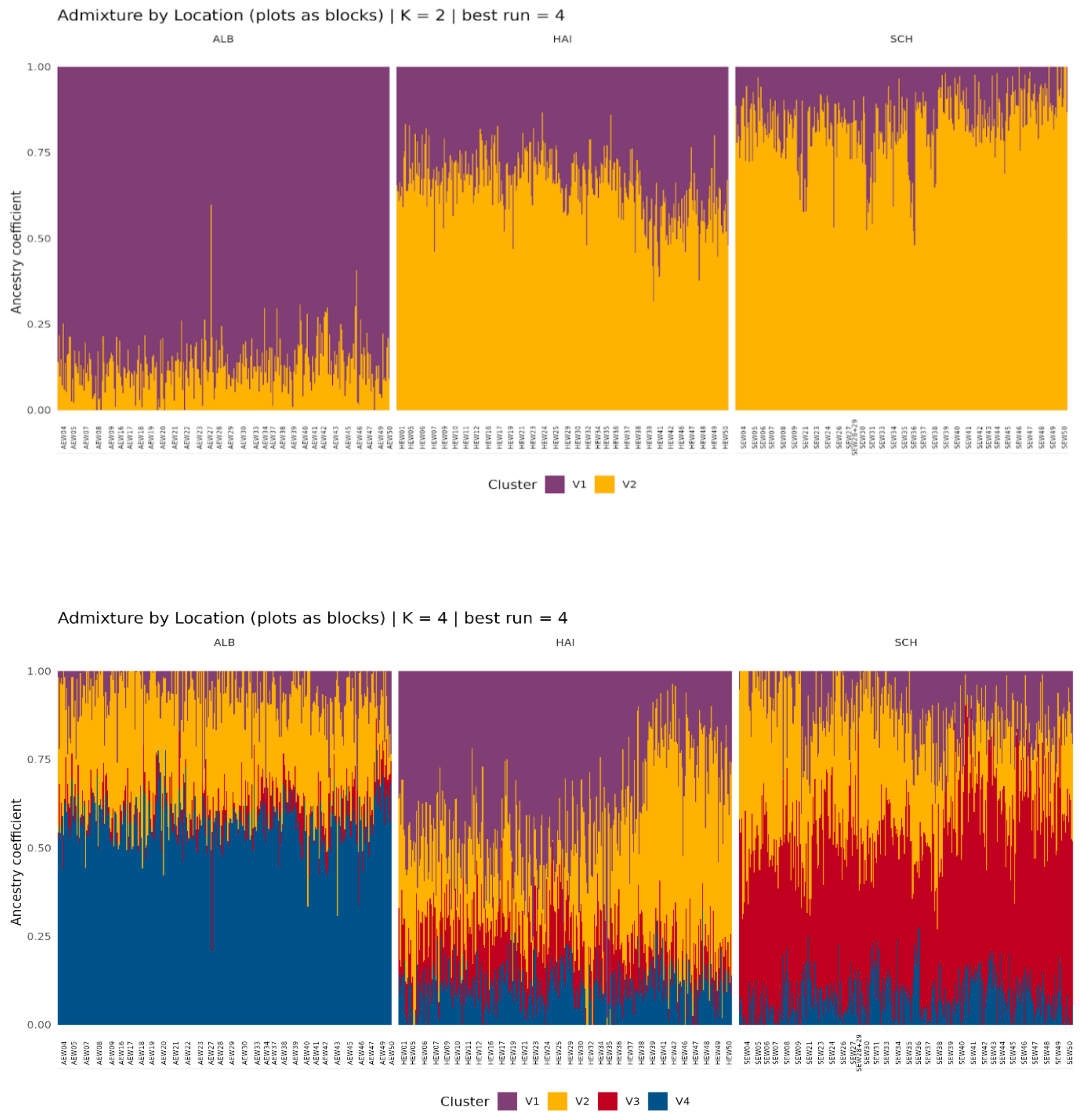


***Figure S5 | Admixture analysis of the three beech populations.***

*(a) Admixture proportions inferred for K = 2 genetic clusters. Individuals from Hainich-Dün (HAI) and Schorfheide-Chorin (SCH) predominantly shared the same ancestry component, whereas individuals from the Swabian Alb (ALB) were largely assigned to a distinct ancestry component. (b) Admixture proportions inferred for K = 4 genetic clusters, revealing additional substructures within regions while maintaining the overall differentiation of ALB from HAI and SCH.*

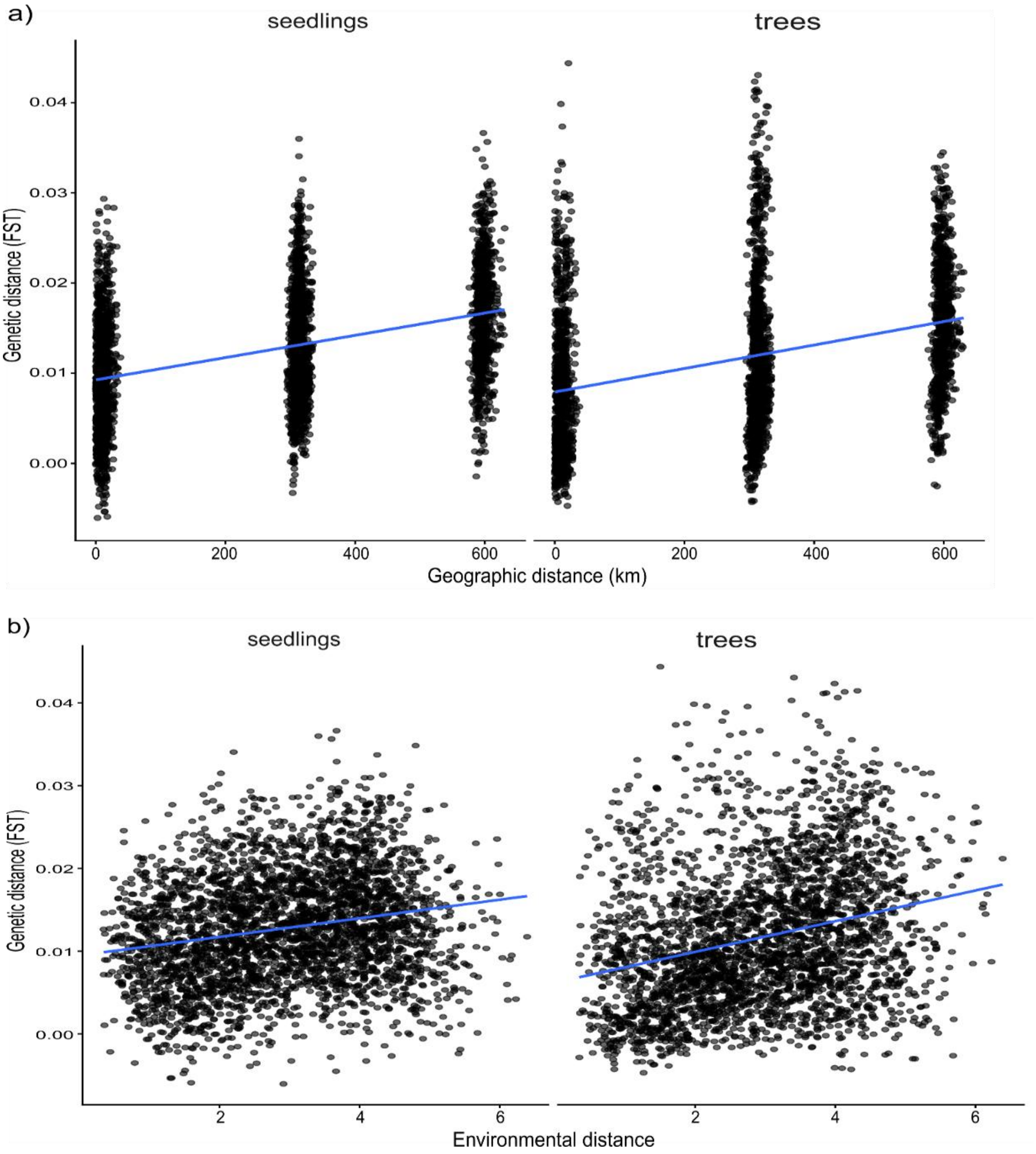


***Figure S6 | Isolation by distance and isolation by environment***

***(a)****Pairwise genetic differentiation ($F_{st}$) between populations related to geographic distances reveals significant isolation-by-distance with a slightly stronger IBD in seedlings (Spearmans r = 0.402,* p<0.001*) compared to trees (Spearmans r = 0.384, p<0.001)* ***(b)*** *Isolation by environment (IBE) is significant for trees (Spearmans r = 0.290, p<0.001) and seedlings (Spearmans r = 0.211, p<0.001). The blue lines show the corresponding linear model.*

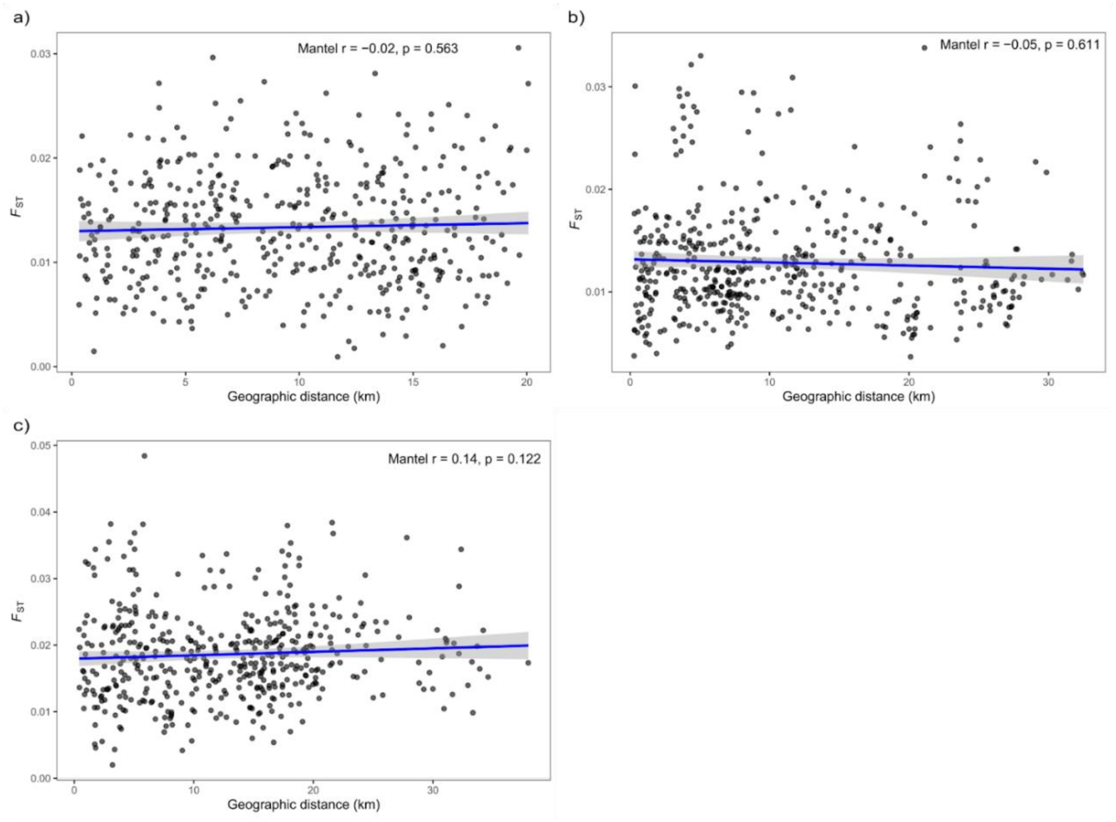


***Figure S7 | Isolation-by-distance within regions.*** *Relationship between geographic and genetic distances among plots within (a) the Swabian Alb (ALB), (b) Hainich-Dün (HAI), and (c) Schorfheide-Chorin (SCH). Mantel tests revealed no significant correlations between geographic and genetic distances within any of the three regions.*

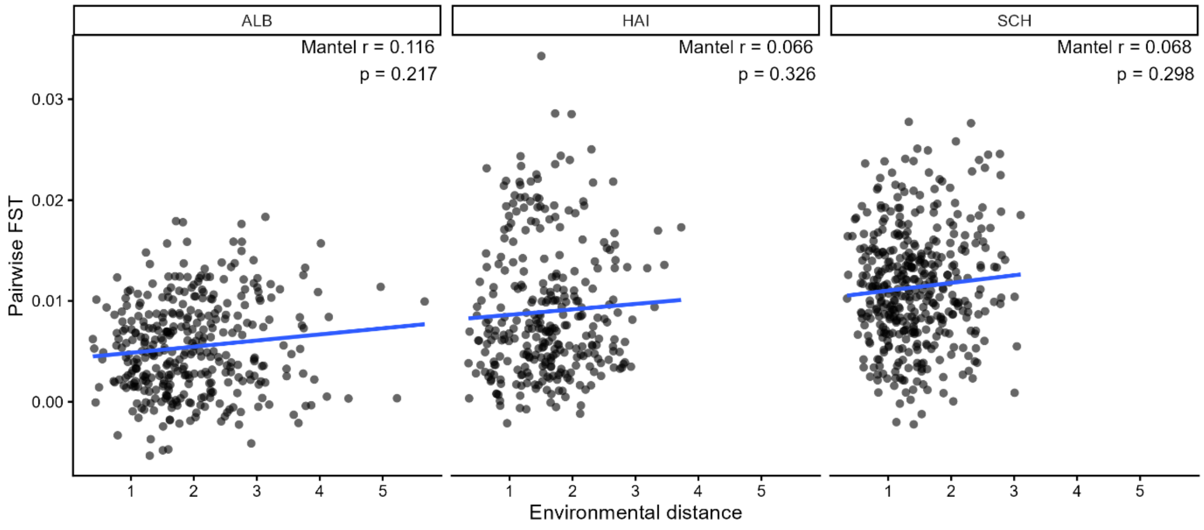


***Figure S8 | Isolation by environment within regions.*** *Relationship between pairwise genetic differentiation (FST) and environmental distance among plots within the three study regions (ALB, HAI, and SCH). Mantel tests revealed no significant isolation by environment within any region. Blue lines show linear regression fits for visualization.*

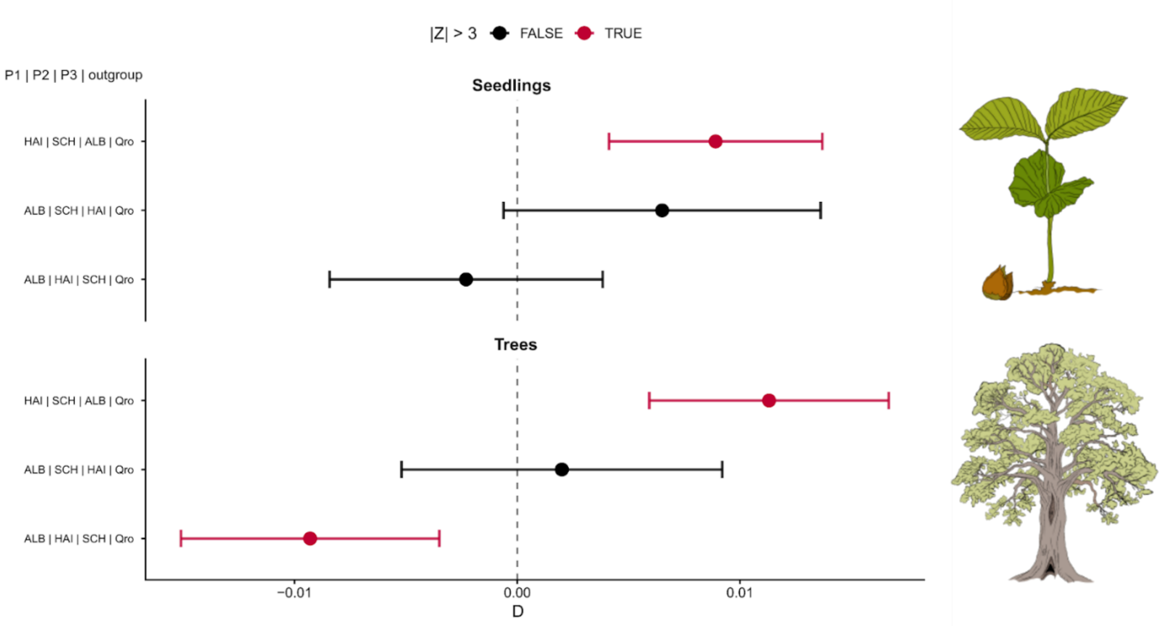


***Figure S9 | D-Statistics for seedlings (top) and trees (bottom)***

*D-statistics (ABBA–BABA tests) for seedlings (top) and trees (bottom) calculated for all combinations of the three study regions using Quercus robur as an outgroup. Error bars represent standard errors. We assessed statistical significance with a block-jackknife approach ($|Z| \geq 3$).*

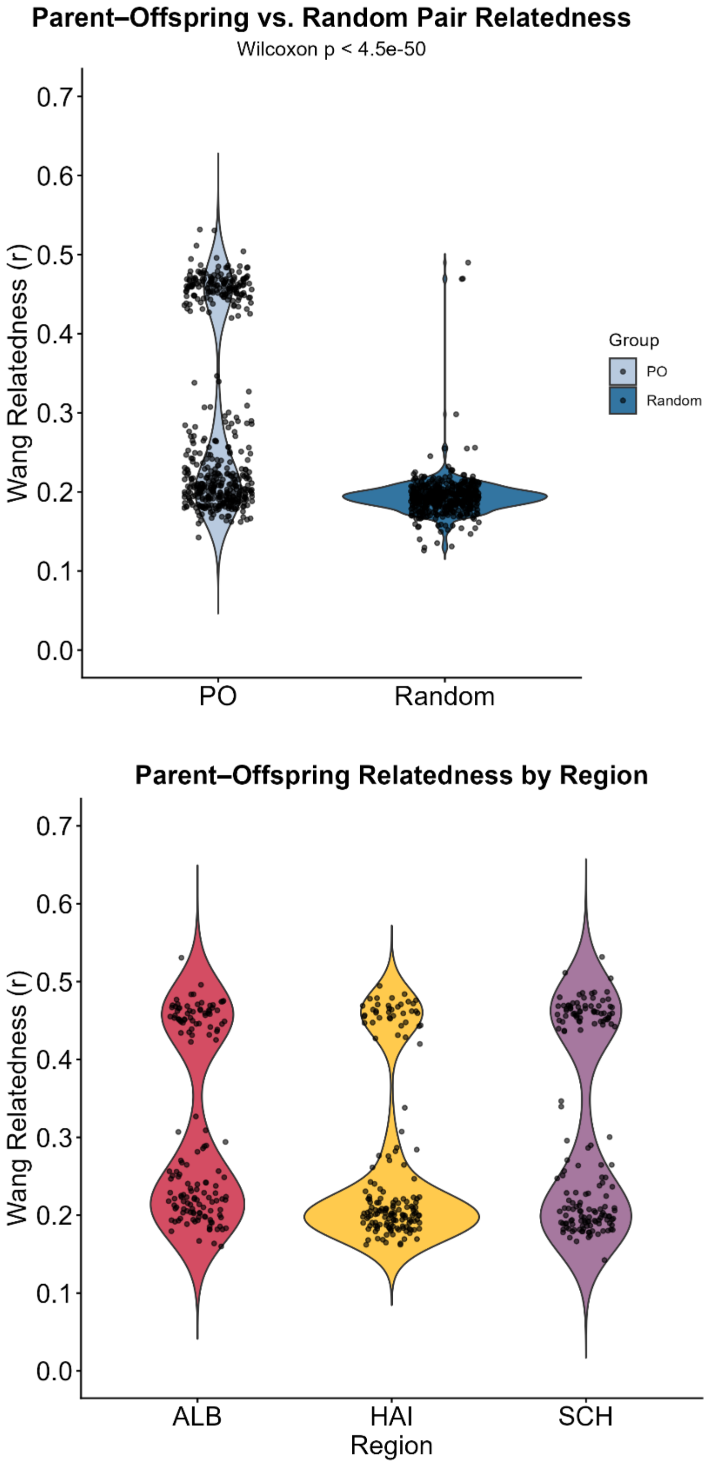


***Figure S10 | Wang relatedness estimates for parent–offspring pairs.*** *(a) Distribution of Wang relatedness values for observed parent–offspring pairs across all samples compared with an artificially generated random sample distribution. (b) Wang relatedness estimates for parent–offspring pairs separated by study region (Swabian Alb, Hainich-Dün, and Schorfheide-Chorin).*

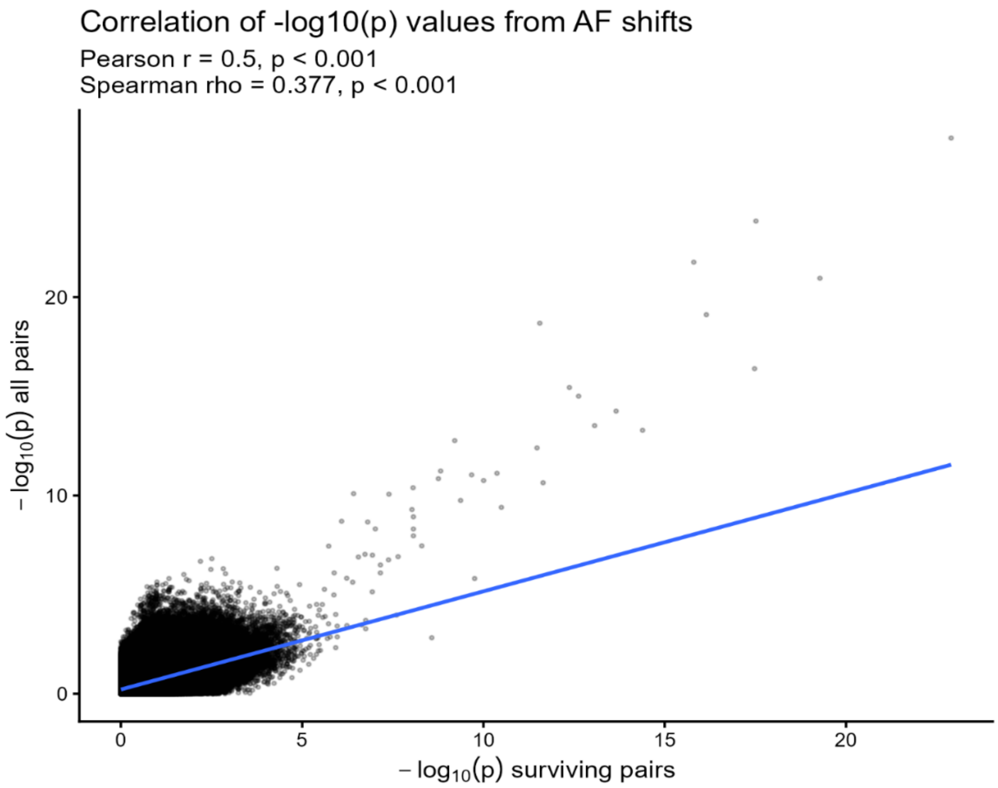


***Figure S11 | Correlation of SNP-wise −log10(p) values between allele-frequency shift analyses based on all tree–seedling pairs and analyses restricted to surviving seedlings.*** *The comparison was used to assess the consistency of statistical signals and to evaluate whether significant allele-frequency shifts were primarily driven by surviving seedlings.*

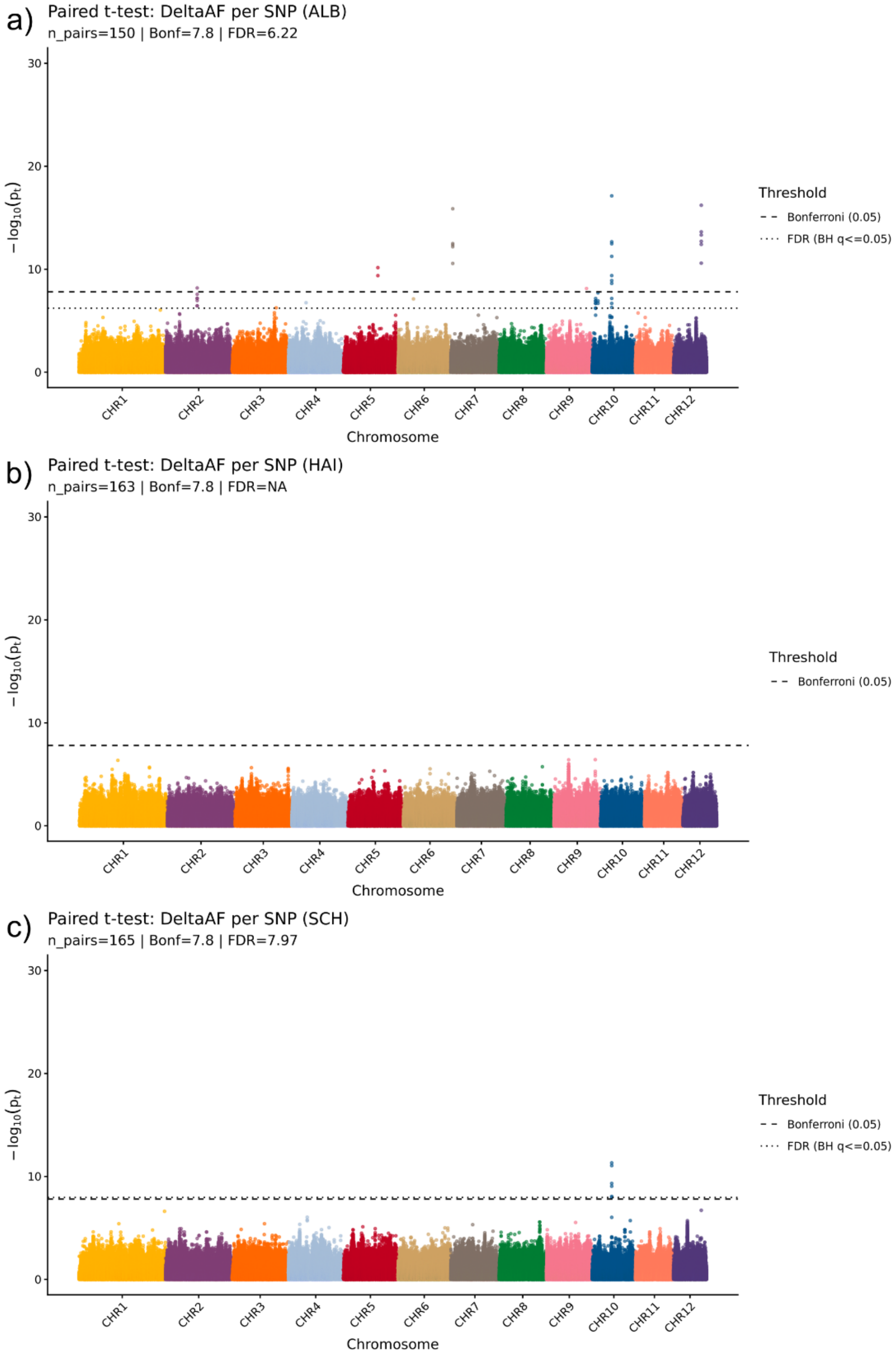


***Figure S12 | Allele-frequency shift analyses based on all tree–seedling pairs*** *(surviving and non-surviving seedlings combined) within (a) Swabian Alb, (b) Hainich-Dün, and (c) Schorfheide-Chorin. Manhattan plots show region-specific distributions of SNP-wise significance values across the genome.*

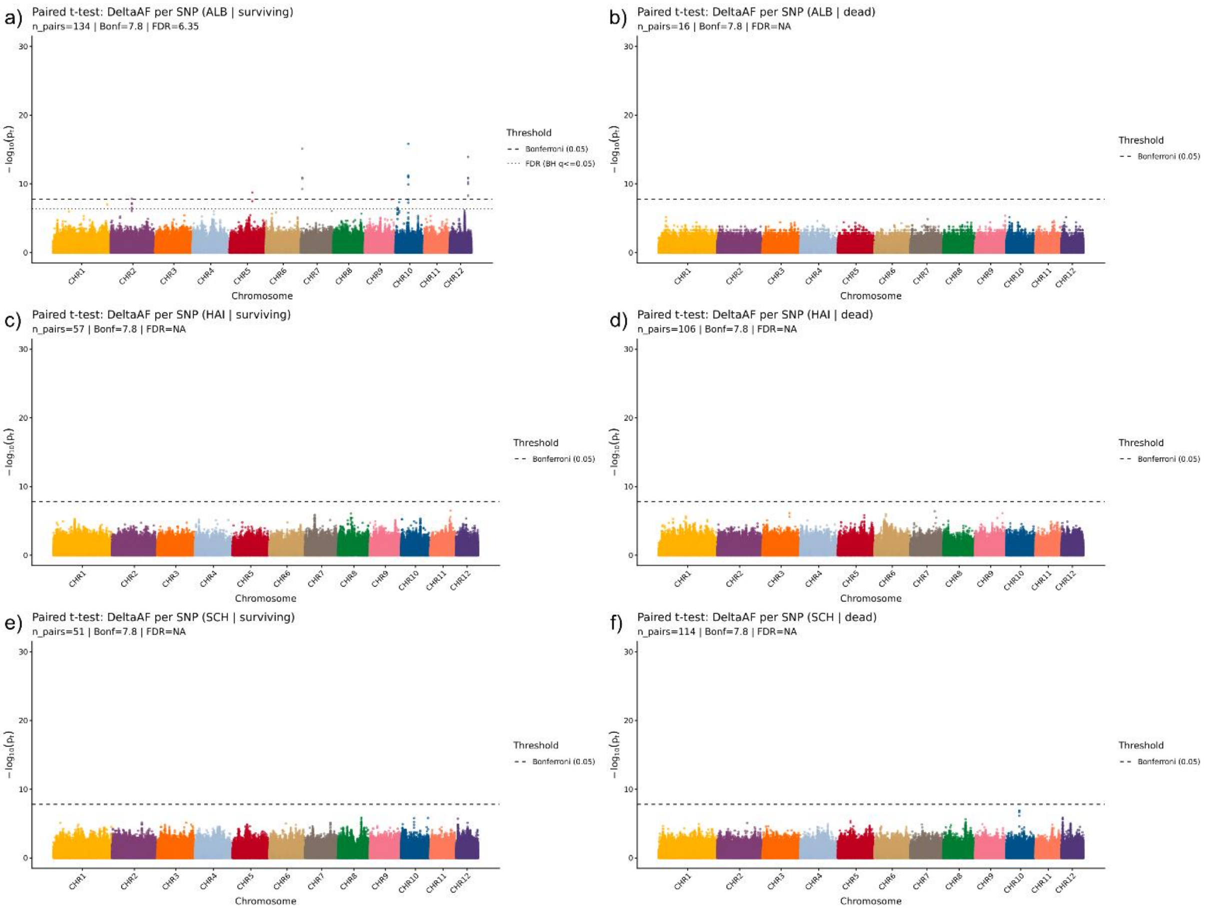


***Figure S13 | Allele-frequency differences between seedlings and trees within each study region, separated by surviving and non-surviving seedlings:*** *(a) Swabian Alb surviving, (b) Swabian Alb dead, (c) Hainich-Dün surviving, (d) Hainich-Dün dead, (e) Schorfheide-Chorin surviving, and (f) Schorfheide-Chorin dead. The outlier signal was detected only in surviving seedlings from the Swabian Alb, whereas no comparable outliers were observed in the remaining comparisons.*

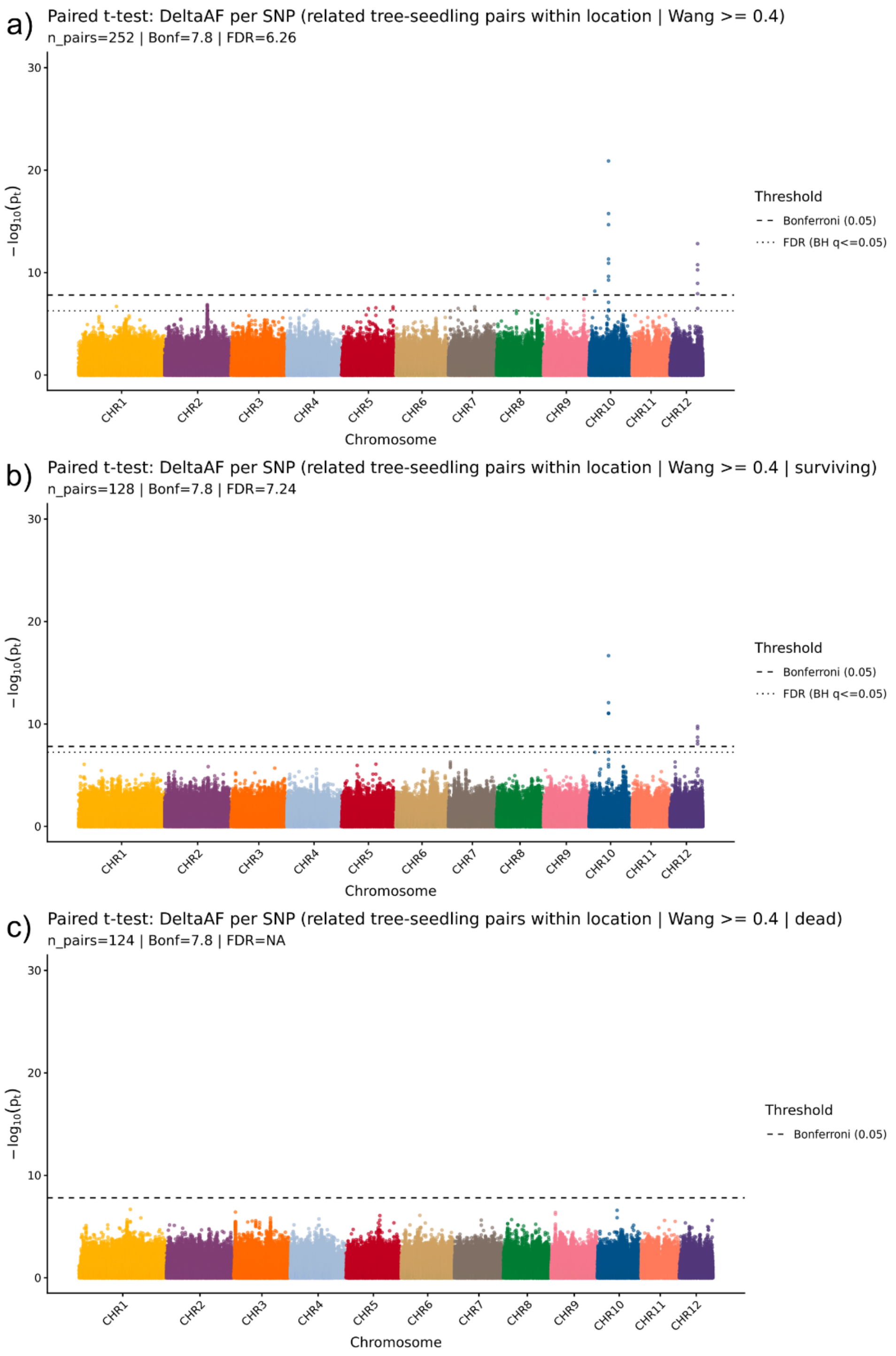


***Figure S14 | a) Closely related tree-seedling pairs (Wang > 0.4)*** *a) All pairs together, b) only surviving seedling-tree pairs and c) only dead seedling-tree pairs that are closely related (Wang >0.4)*

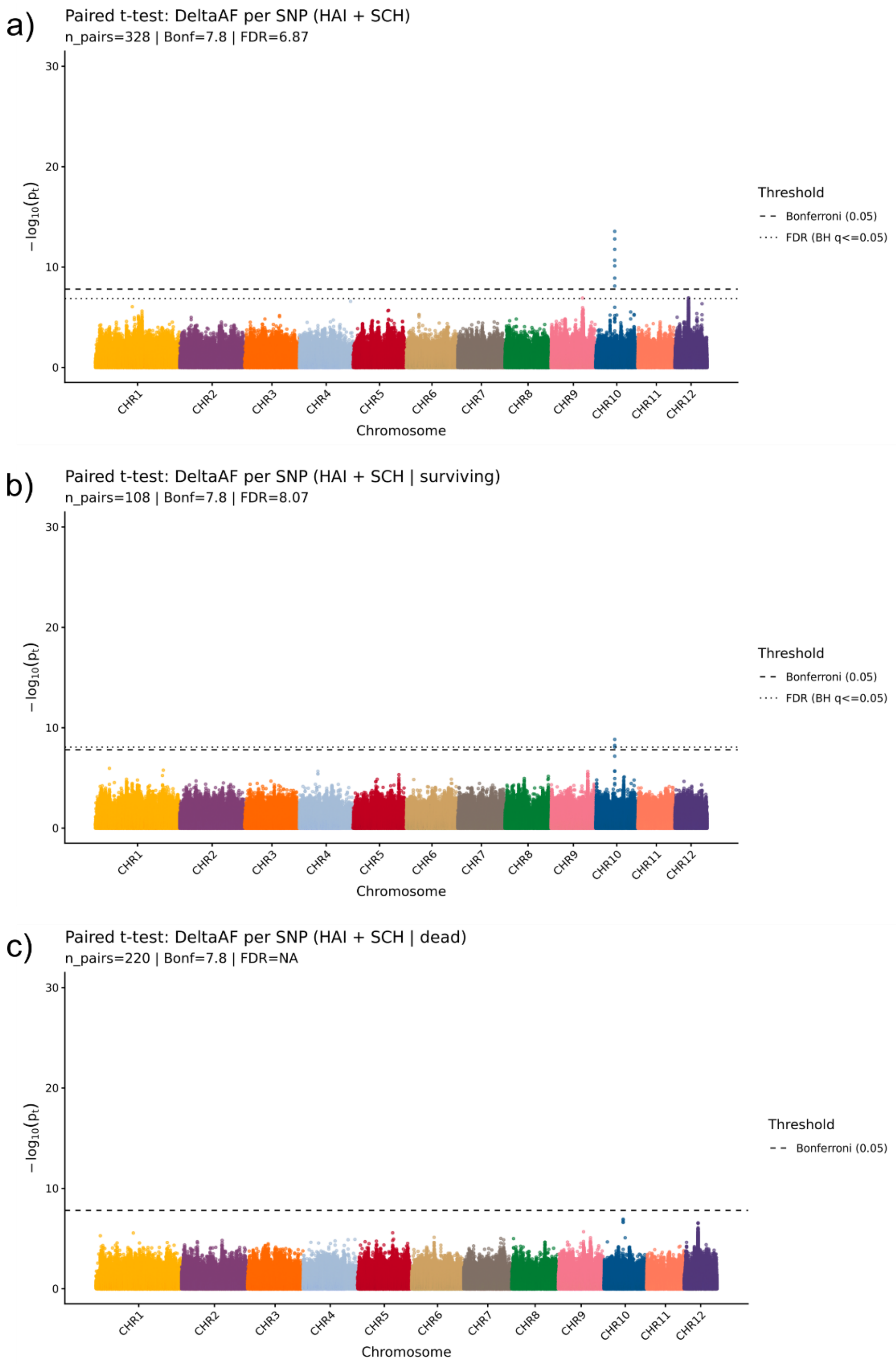


***Figure S15 | Allele frequency differences between seedlings and trees in seedling–tree pairs from Hainich (HAI) and Schorfheide (SCH), where seedlings were established in 2023.*** *a) All seedling–tree pairs from both regions, b) only pairs containing surviving seedlings from both regions, and (c) only pairs containing dead seedlings from both regions.*

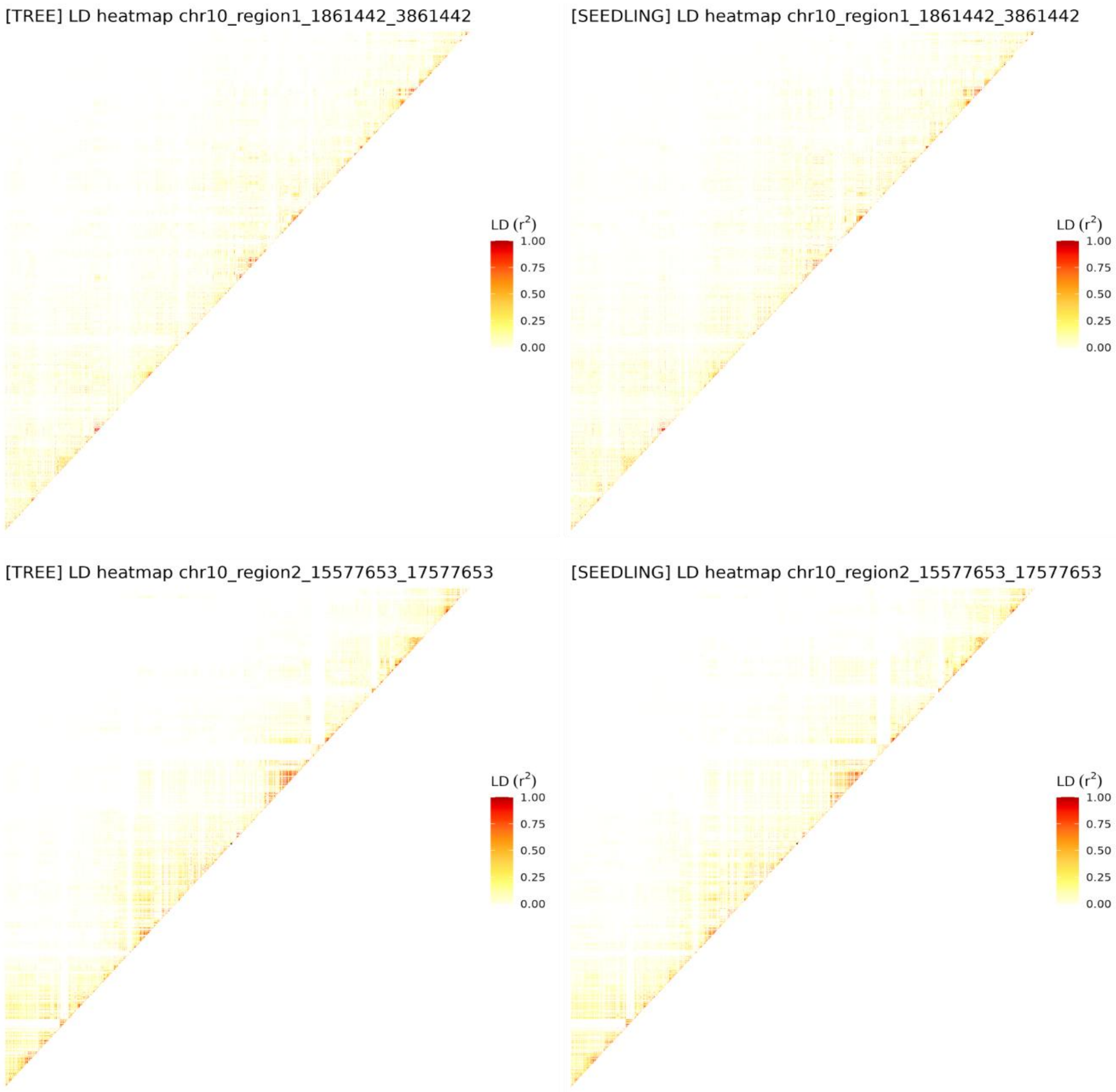
[TREE] LD heatmap chr10_region1_1861442_3861442
LD ($r^2$)
1.00
0.75
0.50
0.25
0.00
[SEEDLING] LD heatmap chr10_region1_1861442_3861442
LD ($r^2$)
1.00
0.75
0.50
0.25
0.00
[TREE] LD heatmap chr10_region2_15577653_17577653
LD ($r^2$)
1.00
0.75
0.50
0.25
0.00
[SEEDLING] LD heatmap chr10_region2_15577653_17577653
LD ($r^2$)
1.00
0.75
0.50
0.25
0.00

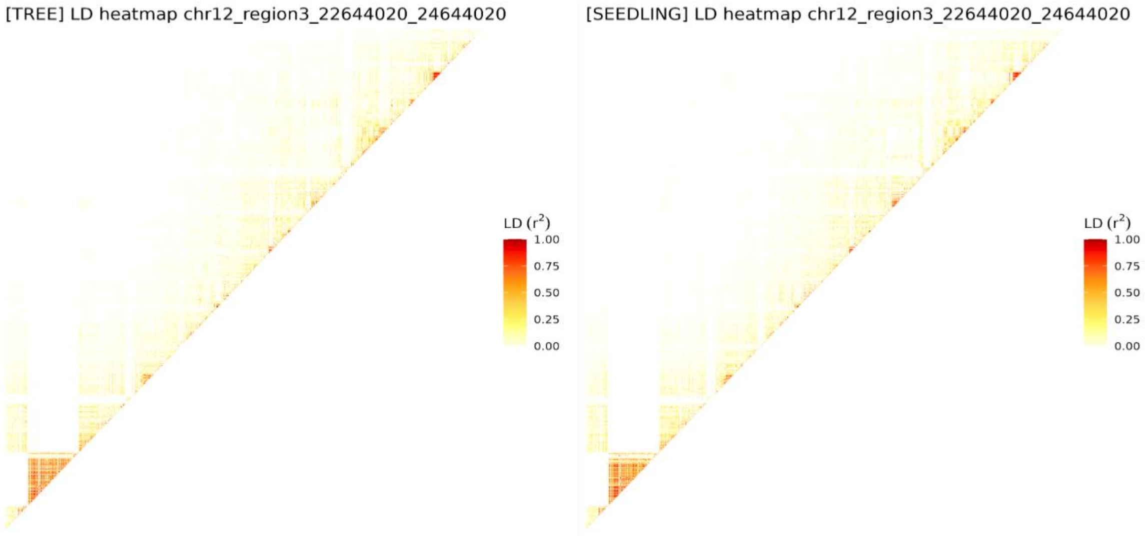
[TREE] LD heatmap chr12_region3_22644020_24644020
LD ($r^2$)
1.00
0.75
0.50
0.25
0.00
[SEEDLING] LD heatmap chr12_region3_22644020_24644020
LD ($r^2$)
1.00
0.75
0.50
0.25
0.00

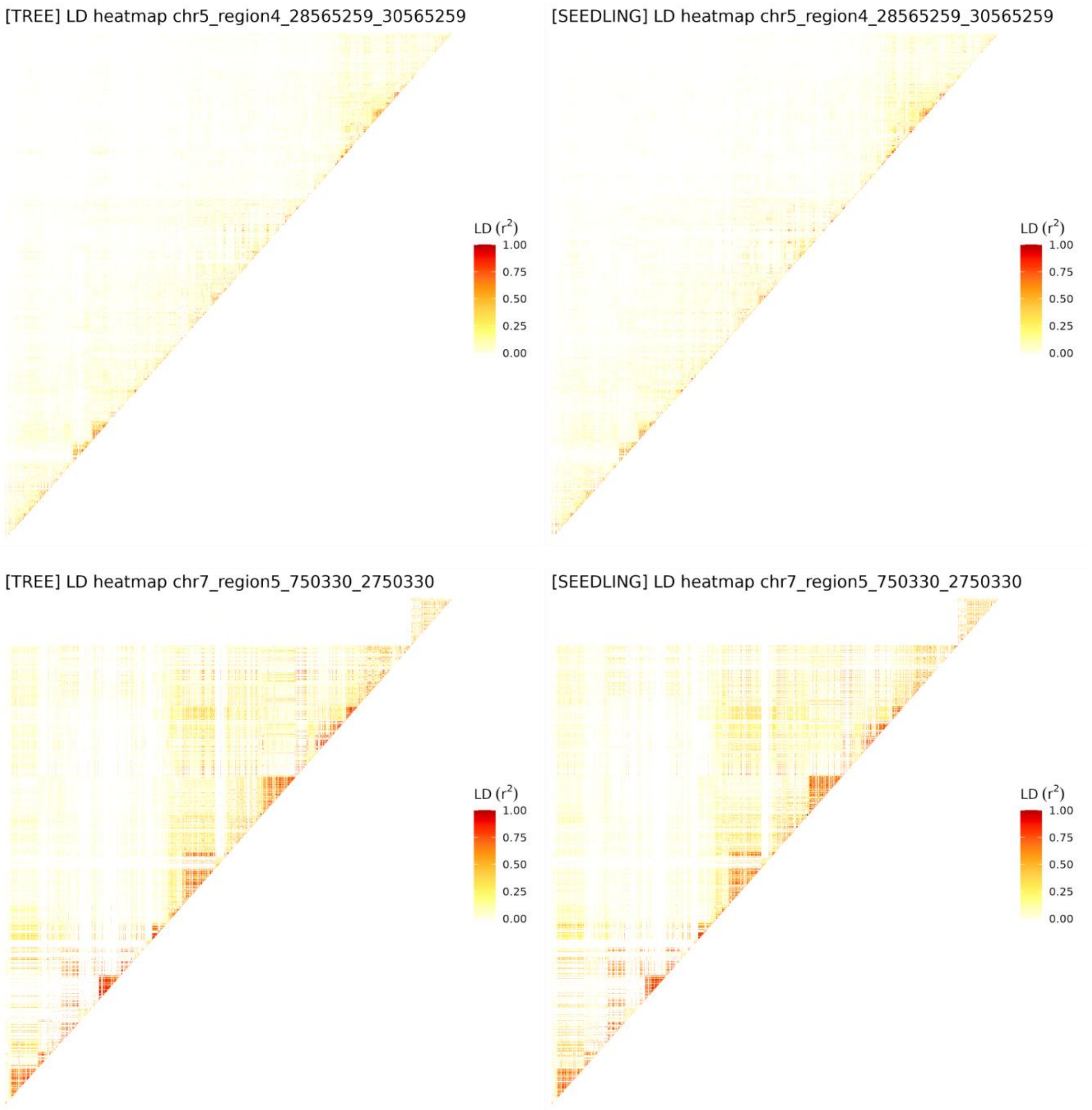


***Figure S16 | Linkage disequilibrium (LD) heatmaps showing pairwise $r^2$ values across 2 Mb genomic windows containing outlier SNPs identified in the allele frequency shift analyses of all pairs and surviving pairs.*** *Allele frequency outlier SNPs are in regions of low local LD.*

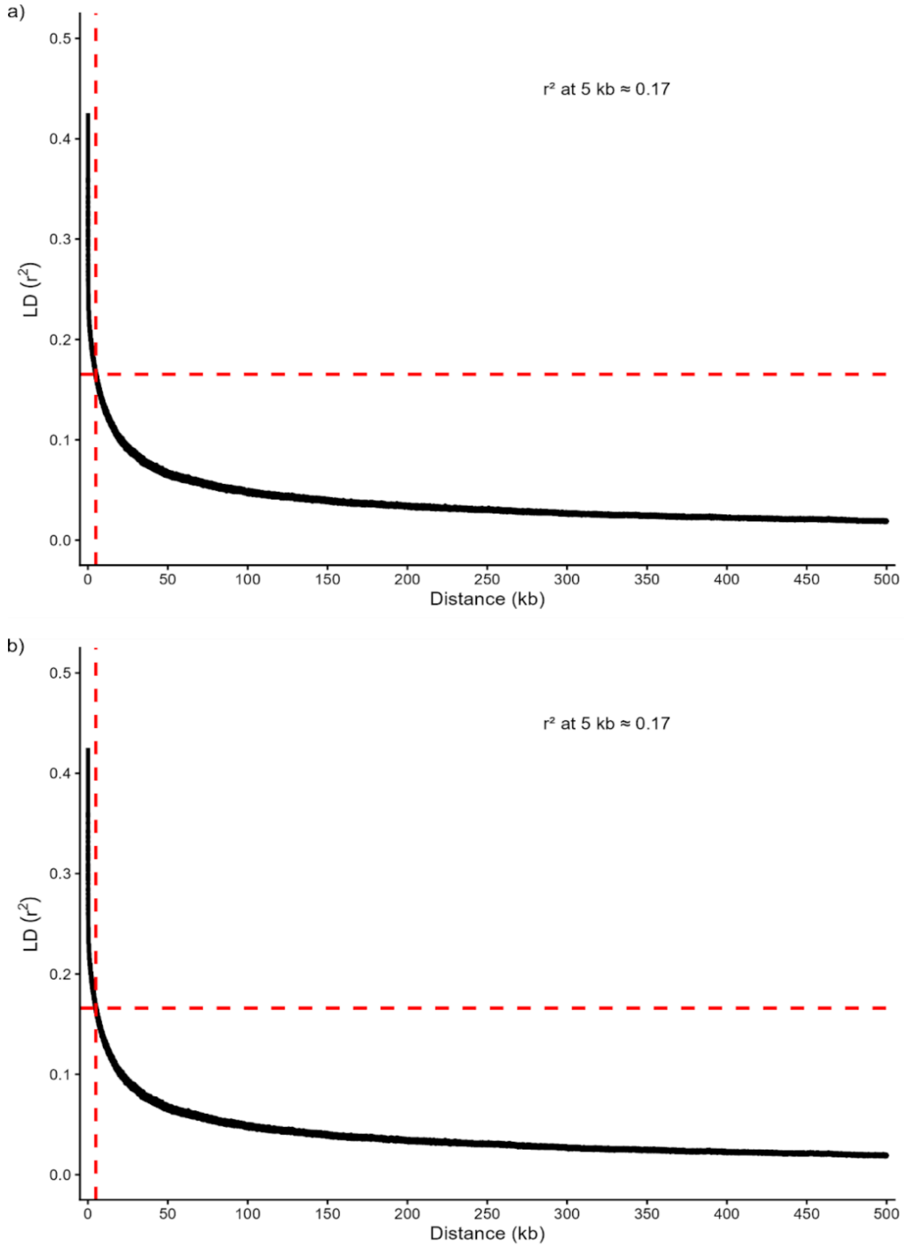


***Figure S17 | Linkage disequilibrium (LD) decay per chromosome for a) trees and b) seedlings.*** *Pairwise LD ($r^2$) is plotted as a function of physical distance between SNPs. Both groups exhibited similar LD decay, with mean $r^2$ of approximately 0.17 at 5 kb.*

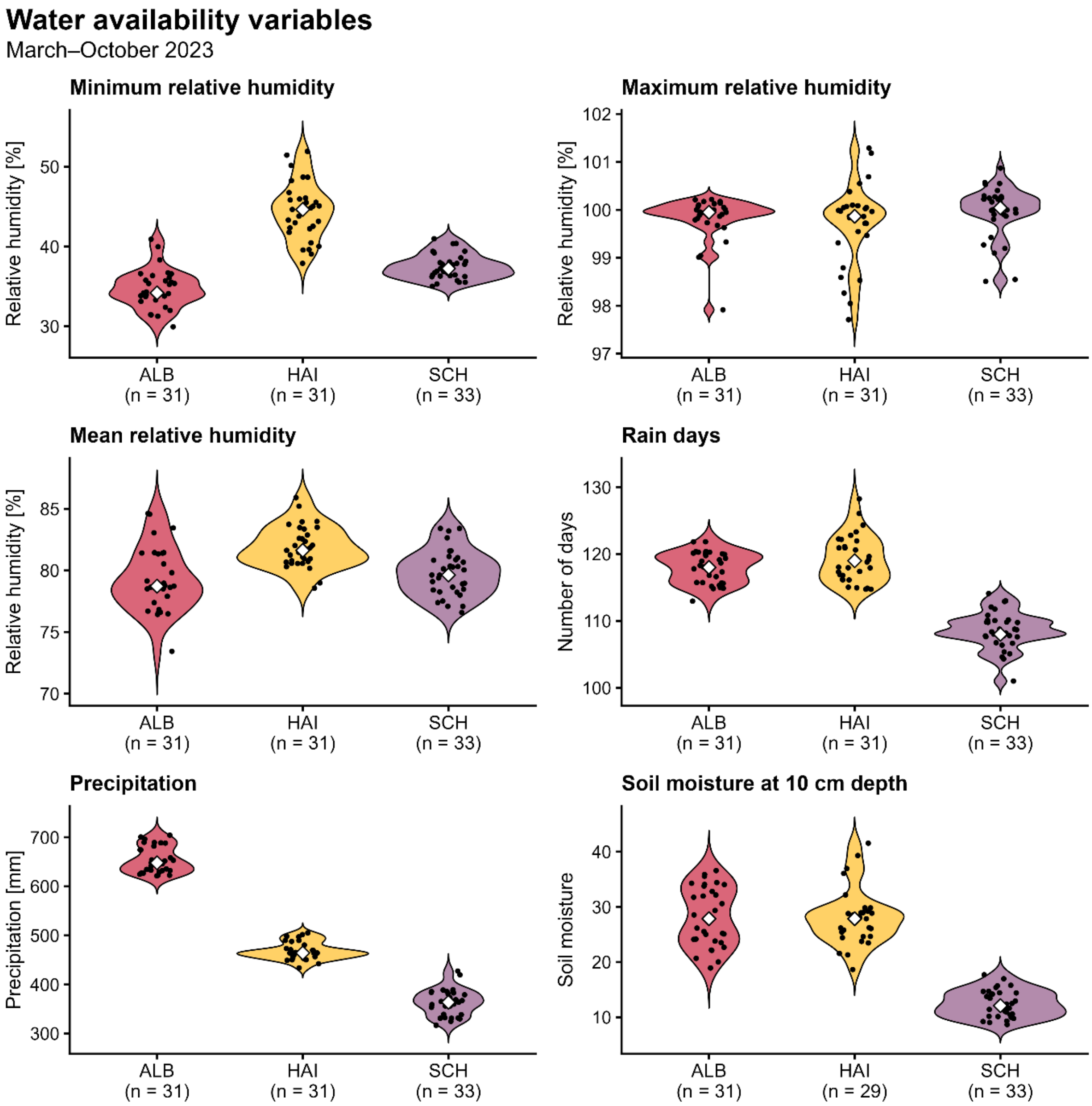


***Figure S18 | Violin plots show the distribution of water availability-related environmental variables across the 95 sampled plots in 2023 for each study location.*** *The distributions indicate regional differences in water availability. Data from Bexis Dataset ID 19007.*

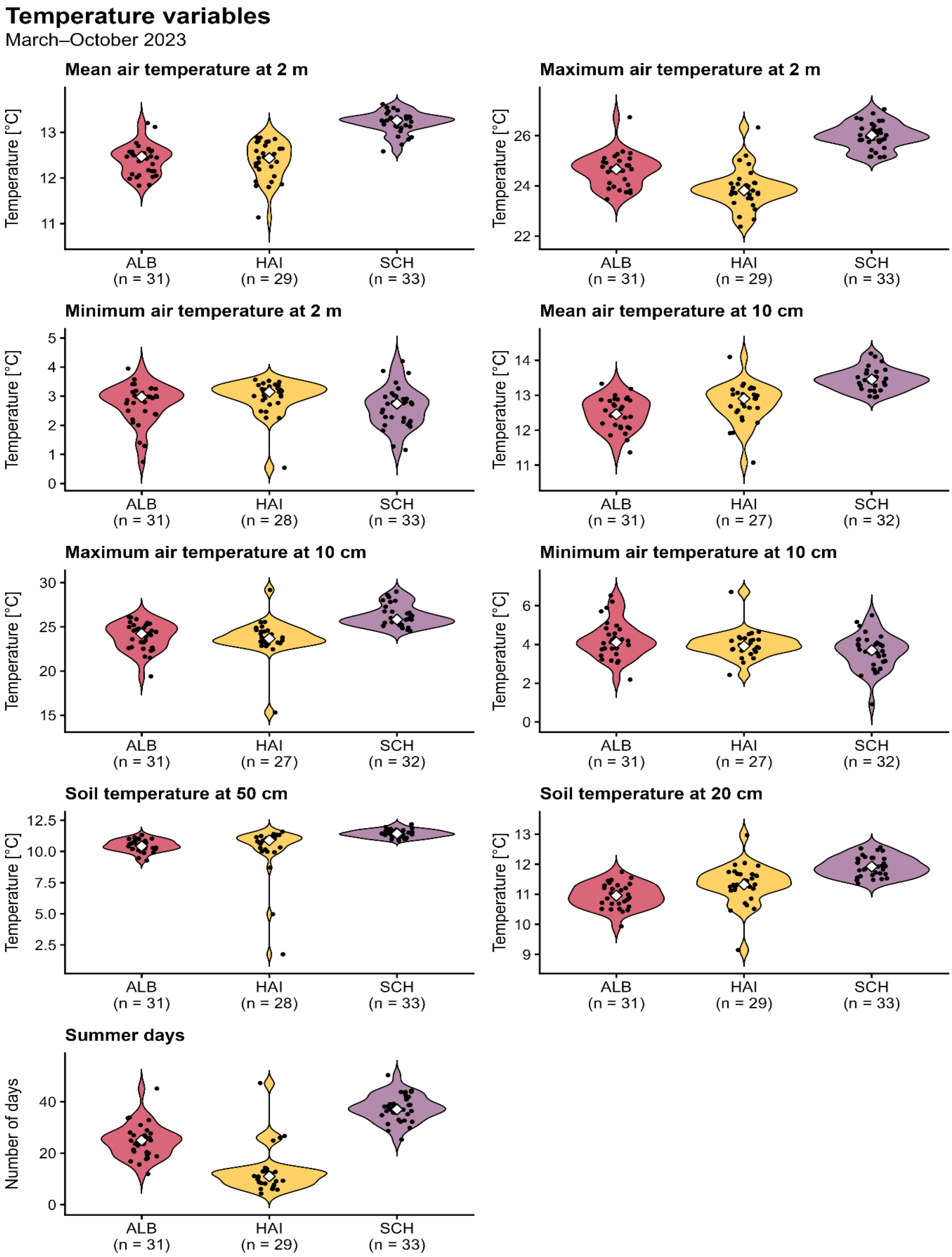


***Figure S19 | Violin plots show the distribution of temperature-related environmental variables across the 95 sampled plots in 2023 for each study location.*** *The distributions indicate regional differences in water availability. Data from Bexis Dataset ID 19007.*

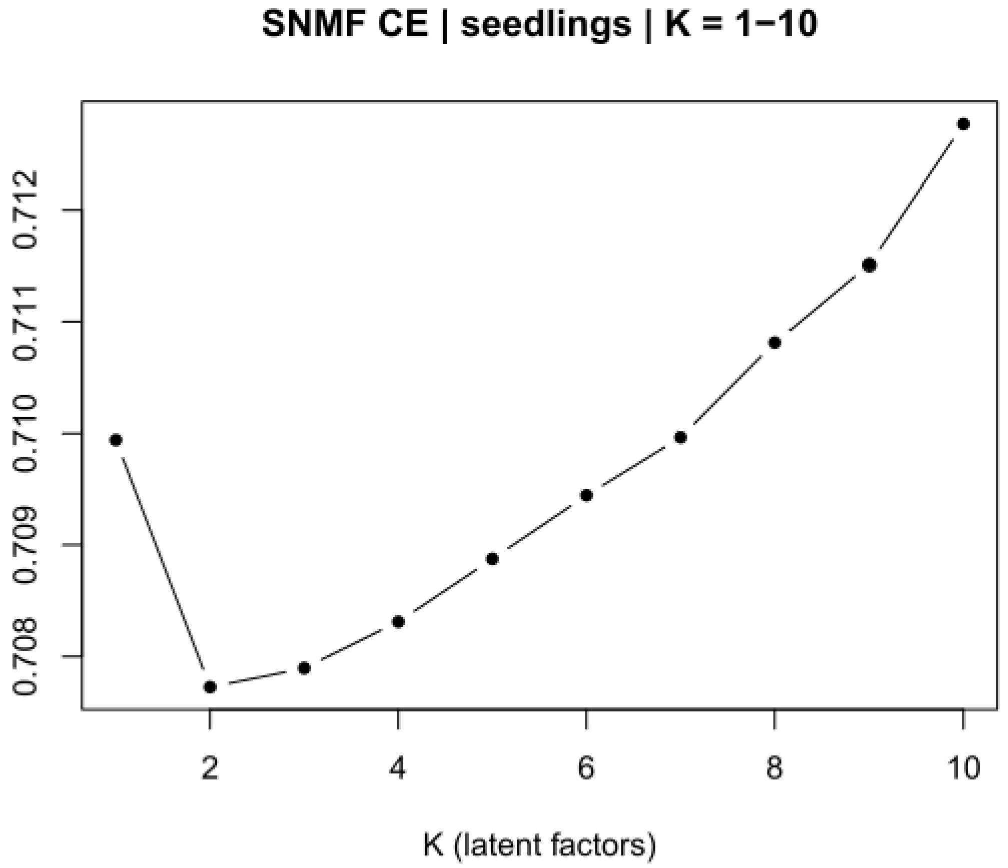


***Figure S20 | Cross-entropy values used to select the number of latent factors (K) for LFMM gene–environment association analyses.*** *The minimum cross-entropy was obtained for K = 2.*

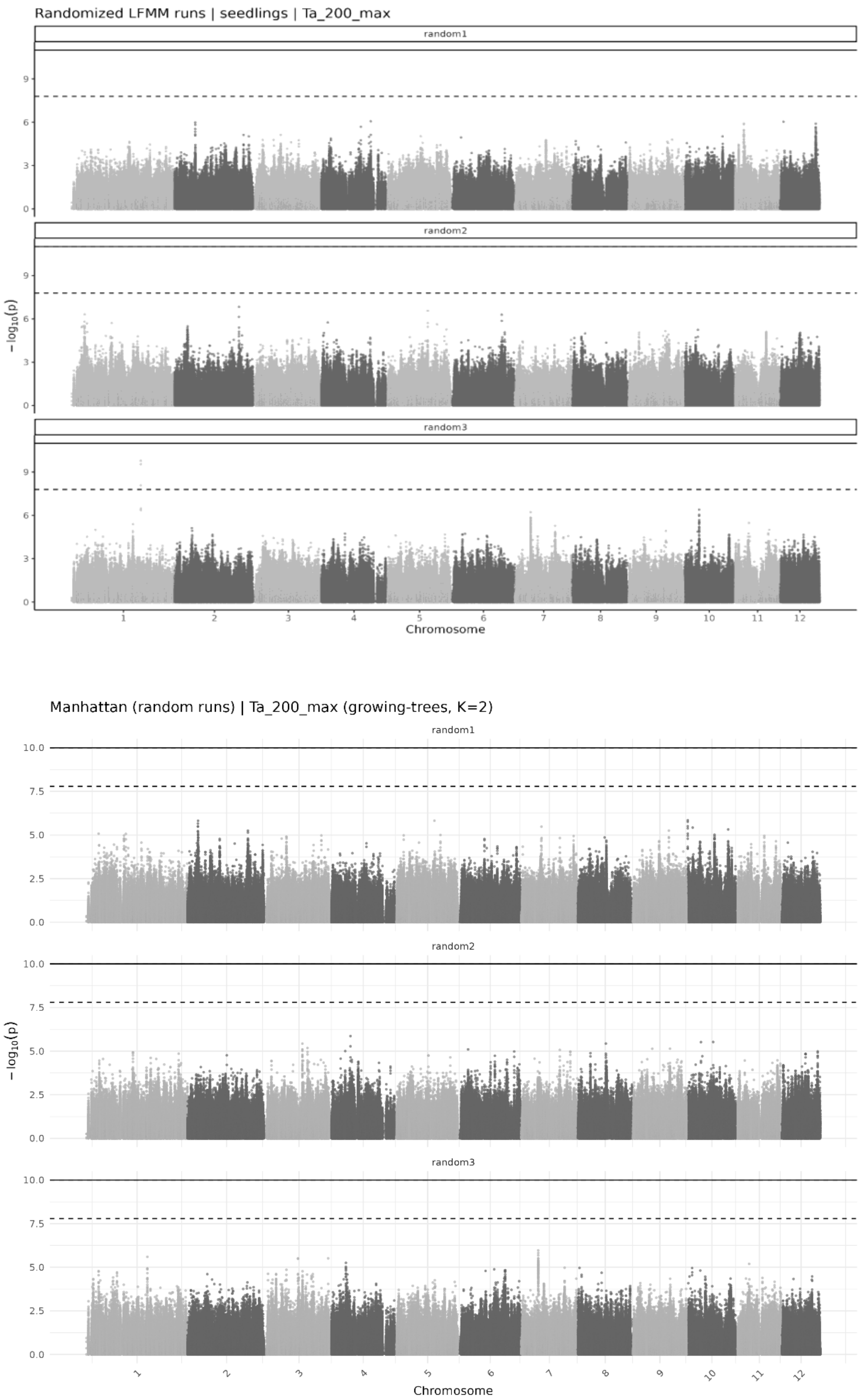


***Figure S21 | Representative randomized gene–environment association (GEA) analyses for maximum air temperature at 2 m in (a) seedlings and (b) trees during the 2023 growing season.*** *The dashed line indicates the false discovery rate (FDR) threshold, and the solid line the Bonferroni threshold. All other randomized analyses showed comparable patterns without significant outlier loci.*

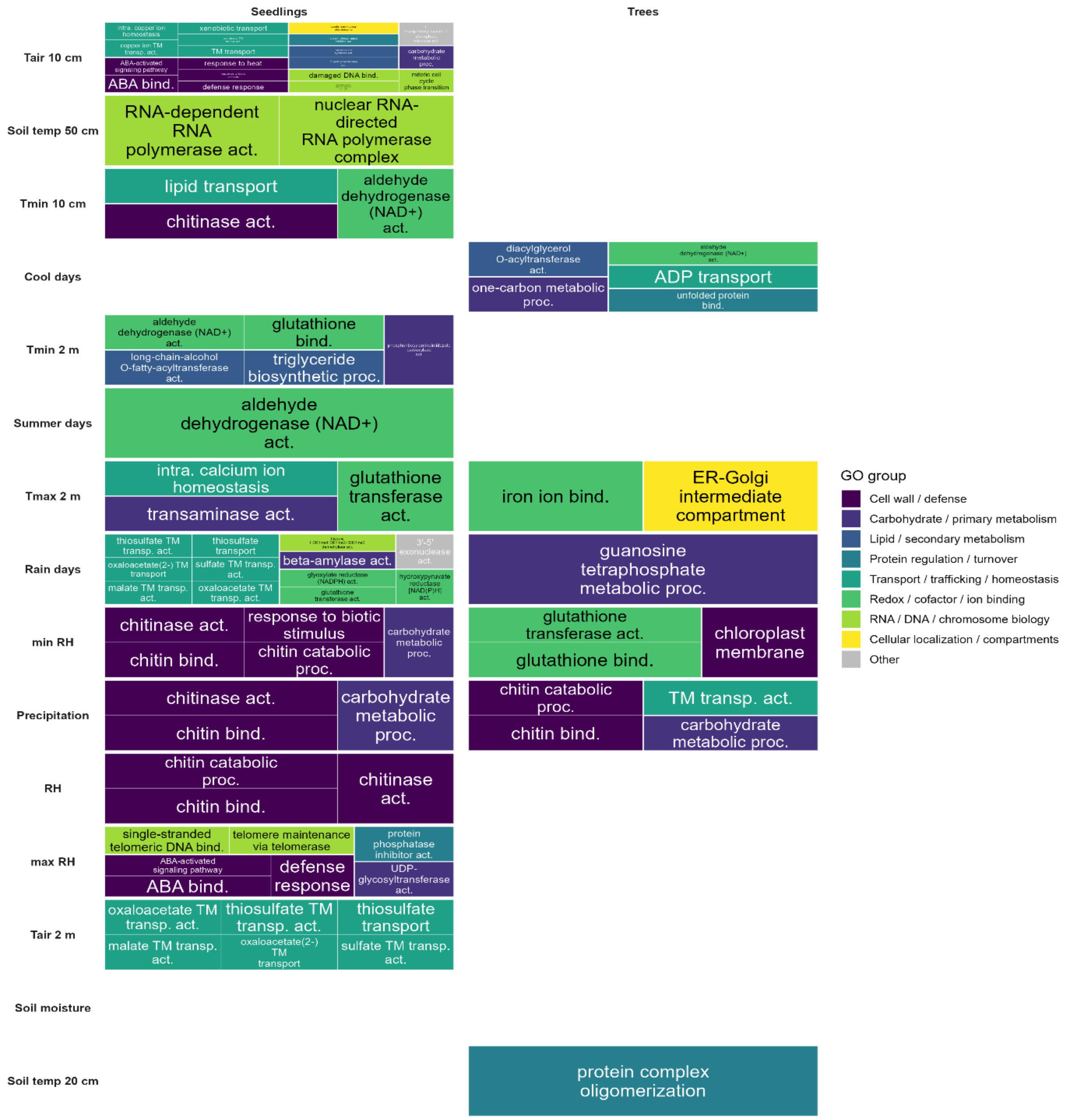


***Figure S22 | Gene Ontology (GO) terms identified by gene set enrichment analysis (GSEA) for seedlings (left) and trees (right) based on environmental variables from the 2023 growing season.*** *Tree map plot summarizes all significantly enriched GO terms identified by gene set enrichment analysis (GSEA) for each environmental variable. Rows correspond to environmental variables, and columns to life stages (seedlings, trees). Each rectangle represents a GO term, grouped into broader functional categories. Environmental variables correspond to the growing season (march – October) 2023. T = temperature; RH = relative humidity.*

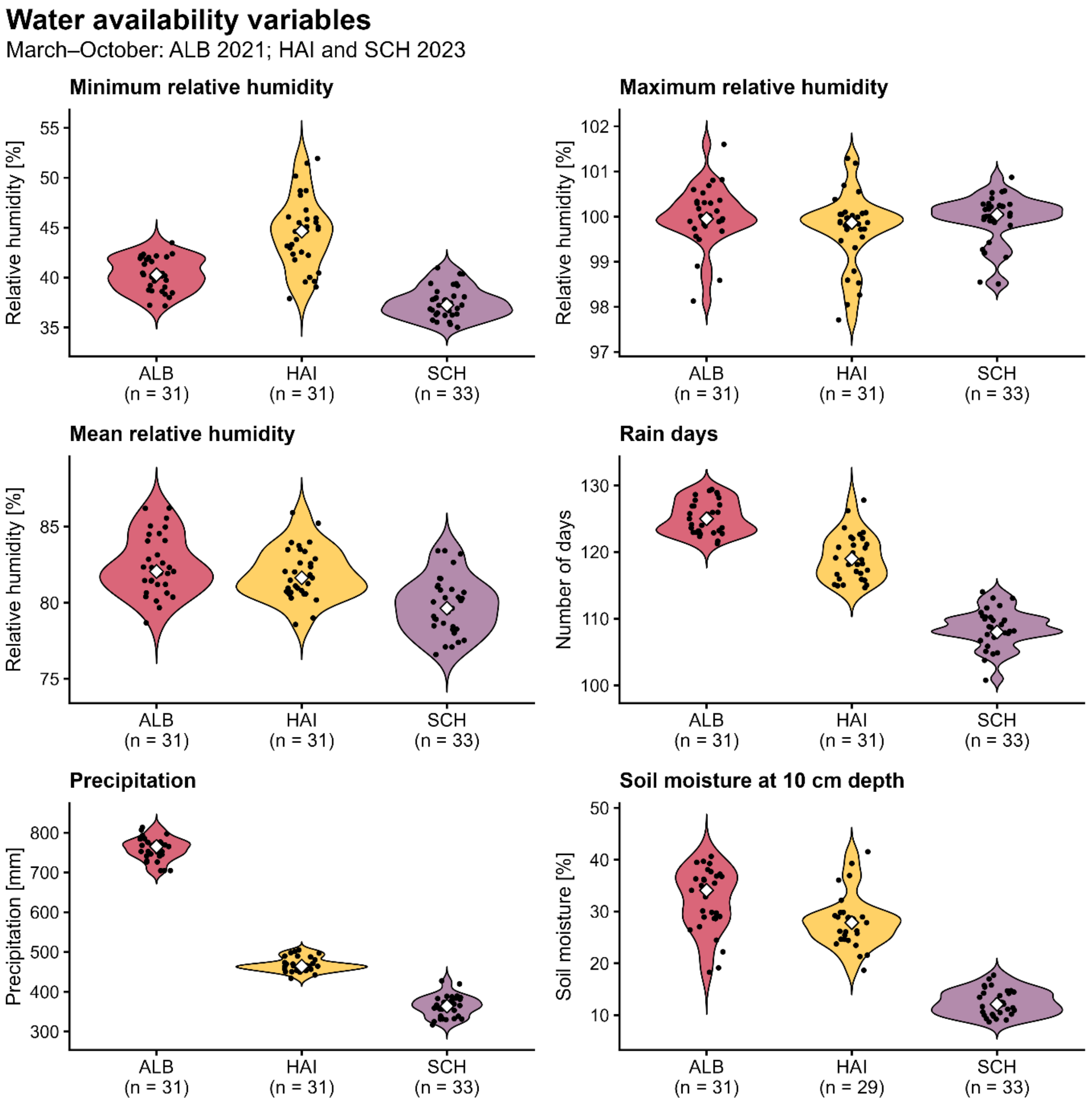


***Figure S23 | Violin plots show the distribution of water-related environmental variables during the year of seedling establishment across the 95 sampled plots for each study region.*** *The year of establishment corresponded to 2021 in the Swabian Alb (ALB) and 2023 in Hainich-Dün (HAI) and Schorfheide-Chorin (SCH). Data were obtained from BExIS (Dataset ID 19007).*

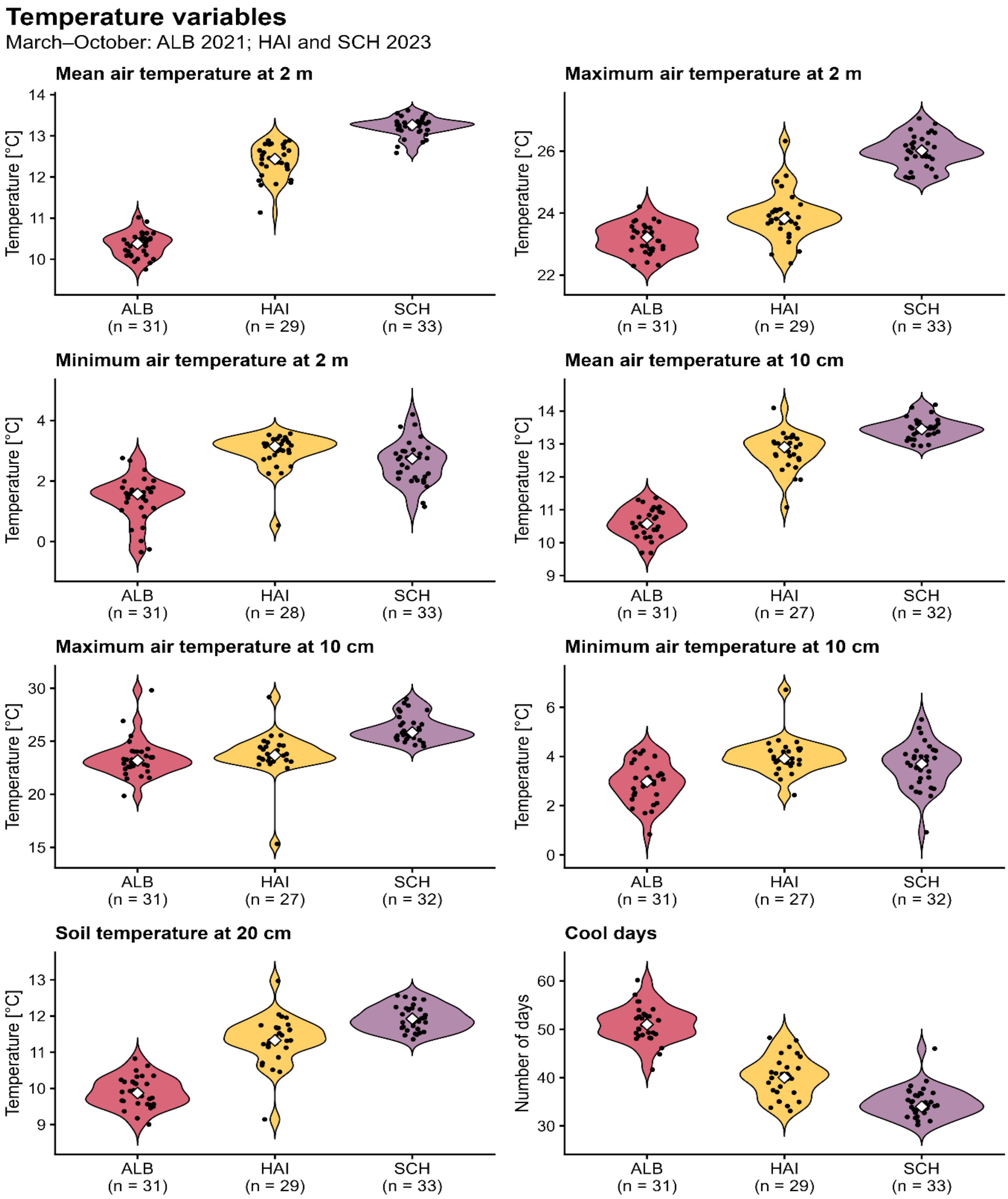


***Figure S24 | Violin plots show the distribution of temperature-related environmental variables during the year of seedling establishment across the 95 sampled plots for each study region.*** *The year of establishment corresponded to 2021 in the Swabian Alb (ALB) and 2023 in Hainich-Dün (HAI) and Schorfheide-Chorin (SCH). Data were obtained from BExIS (Dataset ID 19007).*

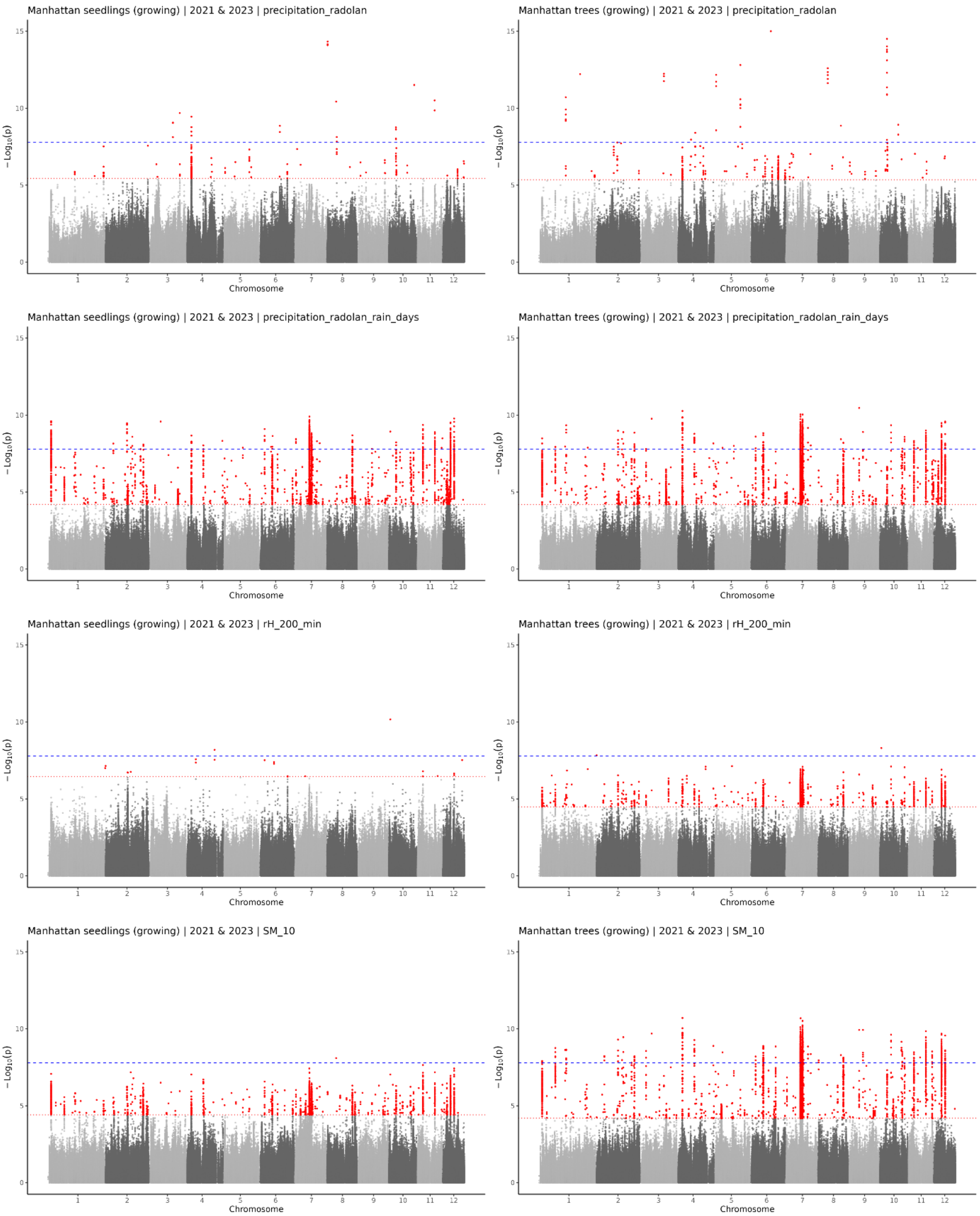


***Figure S25 | Gene–environment association (GEA) manhattan plots using water-related environmental variables from the year of seedling establishment (2021 in ALB; 2023 in HAI and SCH) during the growing season (march – october).***

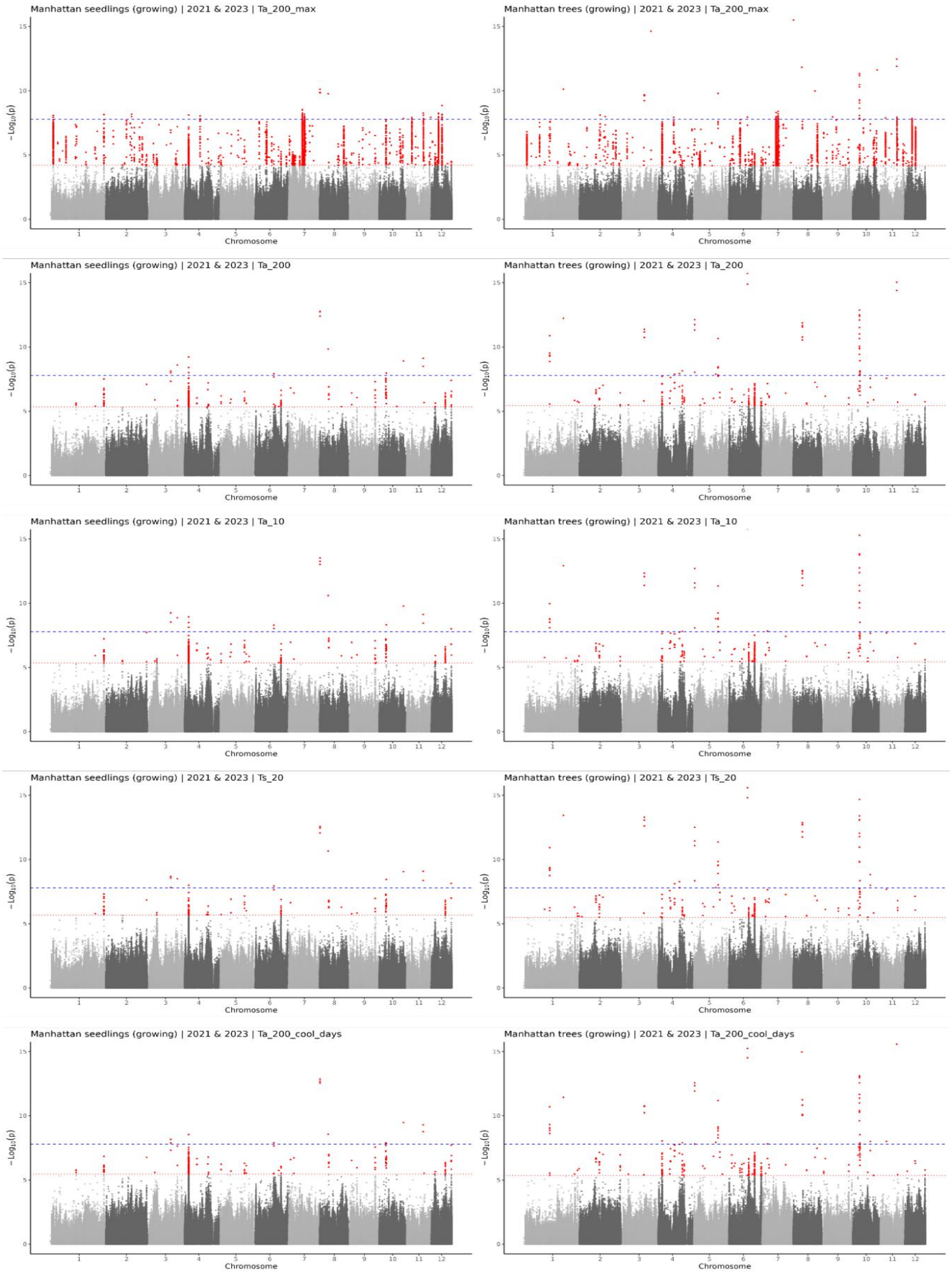


***Figure S26 | Gene–environment association (GEA) manhattan plots using temperature-related environmental variables from the year of seedling establishment (2021 in ALB; 2023 in HAI and SCH) during the growing season (march – october).***

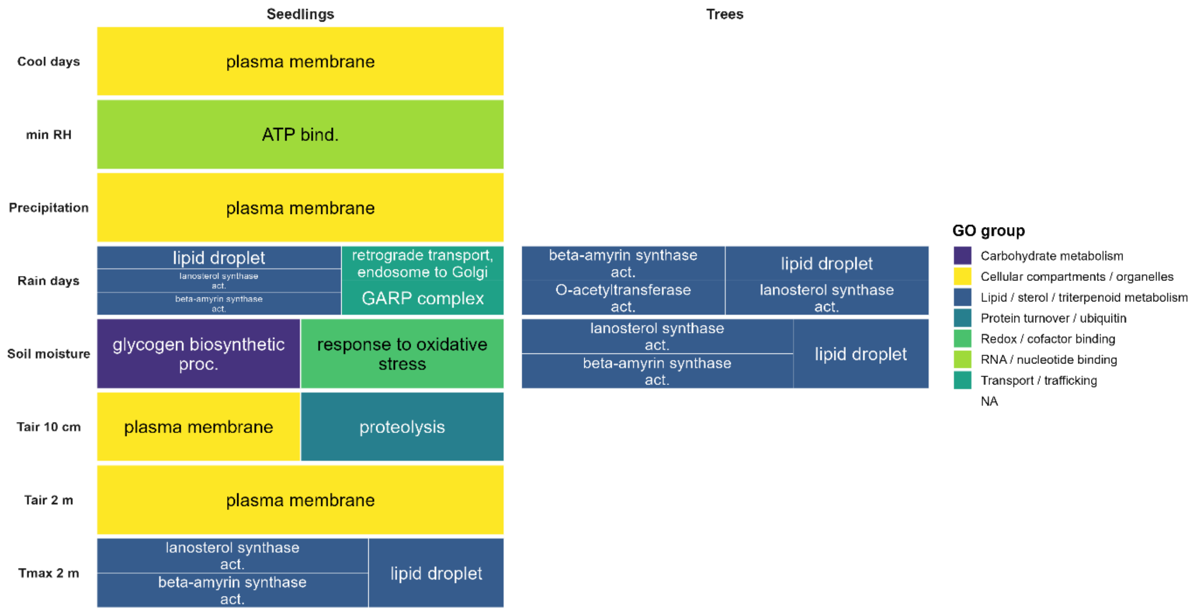


***Figure S27 | Gene Ontology enrichment of LFMM outlier SNPs (ORA) for environmental variables during the year of seedling establishment (growing season).*** *Tree map plot summarizes all significantly enriched GO terms identified by over-representation analysis (ORA) for each environmental variable. Rows correspond to environmental variables, and columns to life stages (seedlings, trees). Each rectangle represents a GO term, grouped into broader functional categories. Environmental variables correspond to the year of seedling establishment (2021 in ALB; 2023 in HAI and SCH). As in the analyses based on 2023 environmental data, seedlings exhibited a greater diversity of enriched GO terms than trees. T = temperature; RH = relative humidity.*

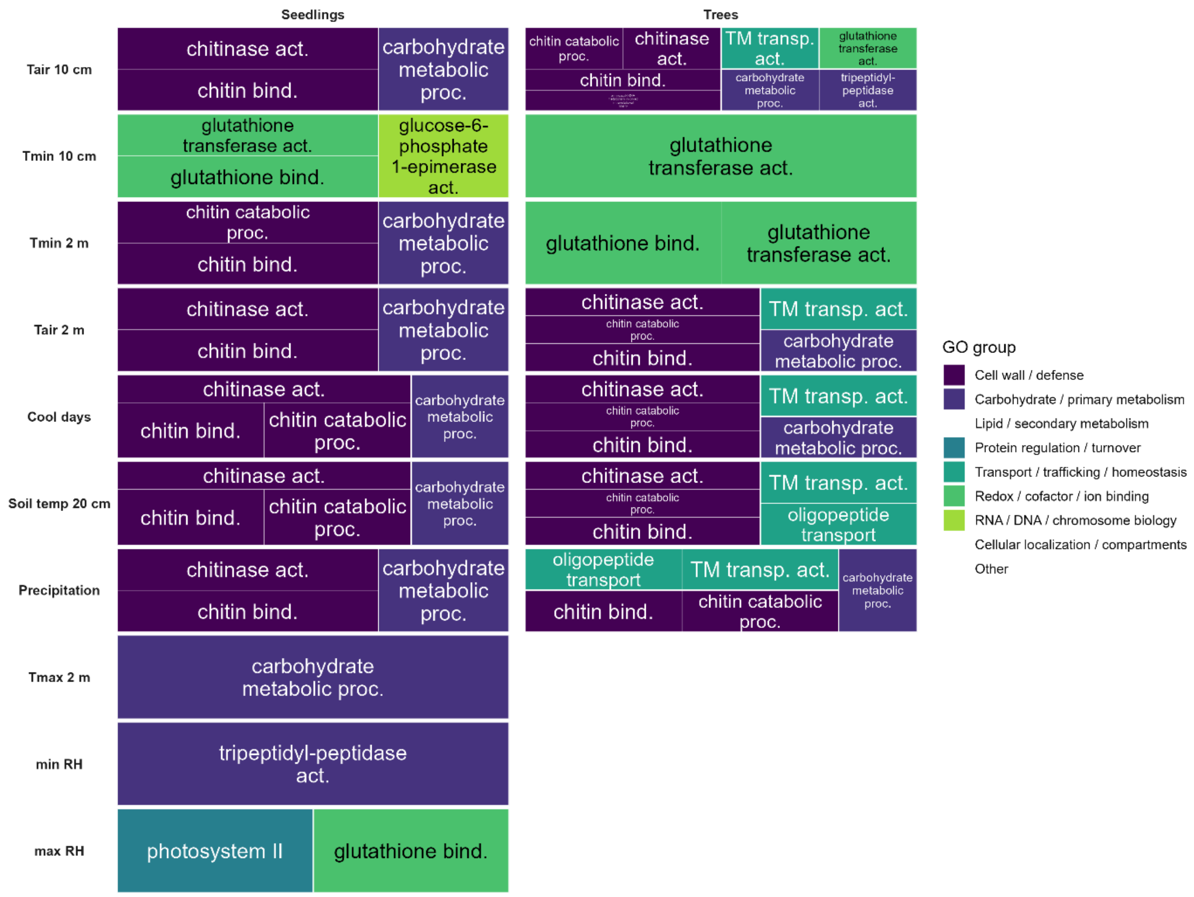


***Figure S28 | Gene Ontology (GO) terms identified by gene set enrichment analysis (GSEA) for seedlings (left) and trees (right) based on environmental variables from the year of seedling establishment during the growing season.*** *Tree map plot summarizes all significantly enriched GO terms identified by gene set enrichment analysis (GSEA) for each environmental variable. Rows correspond to environmental variables, and columns to life stages (seedlings, trees). Each rectangle represents a GO term, grouped into broader functional categories. Environmental variables correspond to the year of seedling establishment (2021 in ALB; 2023 in HAI and SCH). T = temperature; RH = relative humidity.*
